# Xylosyltransferase Bump-and-hole Engineering to Chemically Manipulate Proteoglycans in Mammalian Cells

**DOI:** 10.1101/2023.12.20.572522

**Authors:** Zhen Li, Lucia Di Vagno, Himanshi Chawla, Aisling Ni Cheallaigh, Meg Critcher, Douglas Sammon, David C. Briggs, Nara Chung, Vincent Chang, Keira E. Mahoney, Anna Cioce, Lloyd D. Murphy, Yen-Hsi Chen, Yoshiki Narimatsu, Rebecca L. Miller, Lianne I. Willems, Stacy A. Malaker, Mia L. Huang, Gavin J. Miller, Erhard Hohenester, Benjamin Schumann

## Abstract

Mammalian cells orchestrate signalling through interaction events on their surfaces. Proteoglycans are an intricate part of these interactions, carrying large glycosaminoglycan polysaccharides that recruit signalling molecules. Despite their importance in development, cancer and neurobiology, a relatively small number of proteoglycans have been identified. In addition to the complexity of glycan extension, biosynthetic redundancy in the first protein glycosylation step by two xylosyltransferase isoenzymes XT1 and XT2 complicates annotation of proteoglycans. Here, we develop a chemical genetic strategy that manipulates the glycan attachment site of cellular proteoglycans. By employing a tactic termed bump-and-hole engineering, we engineer the two isoenzymes XT1 and XT2 to specifically transfer a chemically modified xylose analogue to target proteins. The chemical modification contains a bioorthogonal tag, allowing the ability to visualise and profile target proteins modified by both transferases in mammalian cells. The versatility of our approach allows pinpointing glycosylation sites by tandem mass spectrometry, and exploiting the chemical handle to manufacture proteoglycans with defined GAG chains for cellular applications. Engineered XT enzymes permit a view into proteoglycan biology that is orthogonal to conventional techniques in biochemistry.

## Main

Proteoglycans are large biomolecules that consist of a core protein and one or more glycosaminoglycan (GAG) modifications. Ubiquitous on cell surfaces and within the extracellular matrix in metazoa, proteoglycans form a myriad of interactions with cellular receptors as well as soluble signalling molecules, and provide structural support in connective tissues such as cartilage.^1,2^ Growth factors, neurotrophic factors and chemokines can be recruited to target cells through GAG binding sites, rendering proteoglycans important determinants for development.^3,4^ Consequently, dysfunctions in GAG biosynthesis cause severe phenotypes from embryonic lethality to skeletal and muscular deficiencies.^5^ Binding events between proteoglycans and their receptors are impacted by the core protein as well as the identity of GAG polysaccharides that are classified into either heparan sulfate (HS), chondroitin sulfate (CS), dermatan sulfate (DS) or keratan sulfate.^6,7^ Biochemistry and genetic engineering have linked proteoglycan physiology to the GAG structures on particular cell types or even on distinct subcellular locations.^8–13^ Despite their relevance in physiology, only a relatively small number of less than 50 proteoglycans is known in humans.^14,15^ Furthermore, it is challenging to dissect the role of the proteoglycan backbone in physiology from the role of the GAG chain, necessitating strategies in chemistry to manipulate and alter proteoglycans.^16^

An impediment for profiling proteoglycans is the large size of GAG modifications that renders analysis by mass spectrometry (MS) challenging. While GAG-carrying glycopeptides contain common amino acid signatures such as acidic patches and a central O-glycosylated Ser with often flanking Gly or Ala residues, there is no consensus sequence to predict GAG glycosylation in the Golgi.^6,9,17,18^ Common strategies to identify proteoglycans feature enzymatic digestion of GAG chains either before or after isolation of glycopeptides.^9,19–24^ While powerful, such procedures make use of complex digestion and purification protocols and focus solely on the GAG-carrying glycopeptide, without the advantages of shotgun (glyco-)proteomics methods that employ the full MS peptide coverage of individual proteins for detection.

The biosynthesis of HS and CS commences via a common O-linked glycan “linker” modification consisting of a glucuronic acid (GlcA), two galactoses (Gal) and a xylose (Xyl) in the GlcA(β-3)Gal(β-3)Gal(β-4)Xyl(β-)Ser sequence (Fig. 1a), with optional modifications such as phosphorylation on the core Xyl.^7,25^ The first glycosylation step attaching Xyl to Ser is subject to biosynthetic redundancy by the xylosyltransferase isoenzymes XT1 and XT2 that utilize uridine diphosphate (UDP)-Xyl as a substrate. The isoenzymes share 60% amino acid identity but display tissue-specific expression patterns and dysfunctions are associated with different genetic disorders - Desbuquois Dysplasia type 2 and Spondylo-Ocular Syndrome for patient *XYLT1* and *XYLT2* mutations, respectively.^26–29^ Differential roles in physiology have been attributed to XT1 and XT2.^30,31^ Although XT2 appears to be the dominant isoenzyme in cell lines and serum,^32,33^ *Xylt1* and *Xylt2* knockout (KO) mice display differential defects in development.^30,31^ Despite their importance in physiology and the unanswered questions about substrate repertoires, it is currently not possible to directly profile the substrate proteins or even individual glycosylation sites of XT isoenzymes in cells or *in vivo*.

**Fig. 1:**
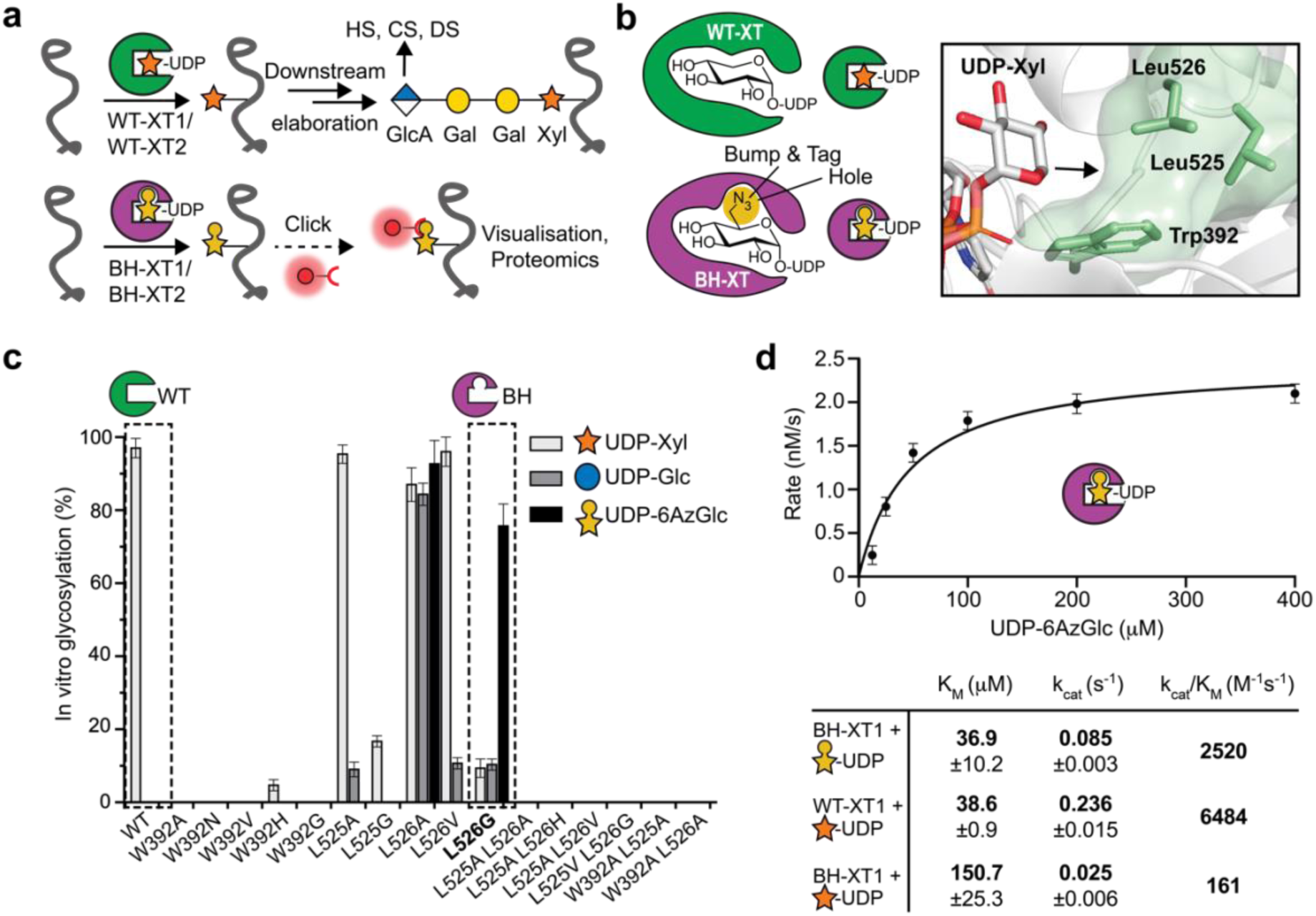
Design of a xylosyltransferase bump-and-hole system. **a**, principle of the BH approach. WT-XTs transfer Xyl to substrate proteins that can be extended to GAG chains. BH-XTs transfer a bioorthogonal Xyl analogue for visualisation and MS profiling. **b**, structural considerations of XT1 engineering to accept UDP-6AzGlc instead of UDP-Xyl. Insert: Gatekeeper residues in the XT1 crystal structure (PDB 6EJ7) and structural trajectory of 6-azidomethyl modification in UDP-6AzGlc. **c**, *in vitro* glycosylation of a fluorescently labelled bikunin substrate peptide by WT- or mutant XT1 and different UDP-sugars. Data are means ± SD from three technical replicates from one out of two independent experiments. **d**, Michaelis-Menten kinetics of *in vitro* peptide glycosylation by different XT1/UDP-sugar combinations. Data are one independent replicate with three technical replicates (top) and average ± range from two independent replicates each (bottom).

Here, we employ a chemical biology tactic termed bump-and-hole (BH) engineering to probe the substrates of human xylosyltransferases in living cells. Based on structural considerations, we mutate a bulky amino acid in the active site of XT1 to a smaller residue to accept the chemically modified substrate UDP-6AzGlc that is not accepted by the wildtype (WT) enzyme. The chemical modification contains an azide group for bioorthogonal incorporation of fluorophores or biotin (Fig. 1a). Judicious choice of the analogue as a derivative of UDP-glucose enables cytosolic delivery, circumventing the lack of a cellular salvage pathway for “direct” analogues of Xyl. After in-depth biochemical characterisation, we install the XT1 BH system in mammalian cells to directly visualise and probe proteoglycans. Using MS glycoproteomics, we find that BH-engineered XT1 modifies the native glycosylation site of a model proteoglycan in mammalian cells. We further show that BH engineering can be applied to the isoenzyme XT2 simply by harnessing sequence conservation in protein engineering, allowing differential substrate profiling of both isoenzymes in mammalian cells. Introduction of a chemical handle at the native glycosylation site enables attachment of a bioorthogonally tagged GAG chain, furnishing “designer” proteoglycans to modulate cellular behaviour. By developing an XT BH system, we chemically manipulate proteoglycans in mammalian cells for applications in MS and functional evaluation.

## Results

### Design of a xylosyltransferase bump-and-hole system

Our XT BH design was prompted by biosynthetic and structural considerations (Fig. 1b). In the absence of a functional group that allows facile chemical modification,^34,35^ the most common approach to develop bioorthogonal reporters of monosaccharides is the replacement of hydroxyl groups with azido groups.^36–38^ Beahm et al. developed a 4-azido-substituted Xyl analogue that is incorporated into proteoglycans by XT1,^39^ but the corresponding UDP-sugar cannot be biosynthesized in mammalian cells since there is no salvage pathway for Xyl.^40^ The bump-and-hole tactic uses substrate analogues that would normally not be accepted by GTs.^41–43^ We sought to employ this feature to our benefit and reprogram XTs to accept a UDP-sugar that is not accepted by WT-XT1 but can be biosynthesized in mammalian cells. WT-XT1 has been reported to use UDP-6AzGlc with approx. 20-fold lower enzymatic efficiency than UDP-Xyl.^44^ We opted to develop a mutant with reversed selectivity to accept UDP-6AzGlc over UDP-Xyl. As an analogue of glucose (Glc), 6AzGlc was projected to hijack parts of the UDP-Glc salvage pathway and therefore allow cellular biosynthesis unlike Xyl analogues.^36^

We recently reported the crystal structure of XT1, revealing a two-lobe architecture containing a catalytic glycosyltransferase domain.^17^ Since XT1 contains an unusually constricted UDP-Xyl binding site that prevents the use of larger UDP-sugars such as UDP-6AzGlc, we deemed it possible to generate additional space (a “hole”) in the active site by mutation. XT1 harbours several bulky “gatekeeper” amino acids in close proximity to C-5 of UDP-Xyl, namely Trp392, Leu525 and Leu526 (Fig. 1b). We designed, expressed and purified from Expi293 cells a total of 16 XT1 single and double mutant variants in which these residues were replaced with smaller amino acids (Fig. 1c, Supporting Fig. 1). *In vitro* glycosylation of a well-known bikunin substrate peptide in an HPLC assay served to assess glycosylation from the sugar donors UDP-Xyl, UDP-6AzGlc and, as a substrate of intermediate size of the “bump”, UDP-Glc.^17^

WT-XT1 displayed exclusive activity for UDP-Xyl in our hands, with no activity towards UDP-Glc or UDP-6AzGlc.^44^ Most engineered XT1 variants displayed either no activity at all or were still selective for UDP-Xyl, with some displaying activity toward UDP-Glc. Strikingly, the mutant Leu526Gly preferred UDP-6AzGlc as a substrate, with 7-8-fold higher turnover than using UDP-Xyl or UDP-Glc in an endpoint assay. Compared to the Leu526Gly mutant (henceforth termed “BH-XT1”), the construct Leu526Ala displayed no such selectivity, with equal activity on all three UDP-sugars (Fig. 1c). We determined optimal enzyme concentrations and kinetic constants for the native and BH enzyme-substrate pairs (Fig. 1d, Supporting Figs 2, 3). We found that the K_M_ of the BH pair (36.9 µM) was similar to the WT pair (38.6 µM), while the k_cat_ was 2.8-fold reduced in the BH pair. In contrast, BH-XT1 uses UDP-Xyl with an approx. 10-fold lower catalytic efficiency than UDP-6AzGlc, suggesting that the native substrate UDP-Xyl should not be able to outcompete UDP-6AzGlc in cellular applications. Taken together, we established a sensitive structure-activity relationship in the development of a XT1 bump-and-hole variant. BH-XT1 fulfilled the crucial pre-requisite of preferring a chemically modified substrate that is not used by the WT enzyme, with comparable kinetics to the WT enzyme-substrate pair.

**Fig. 2:**
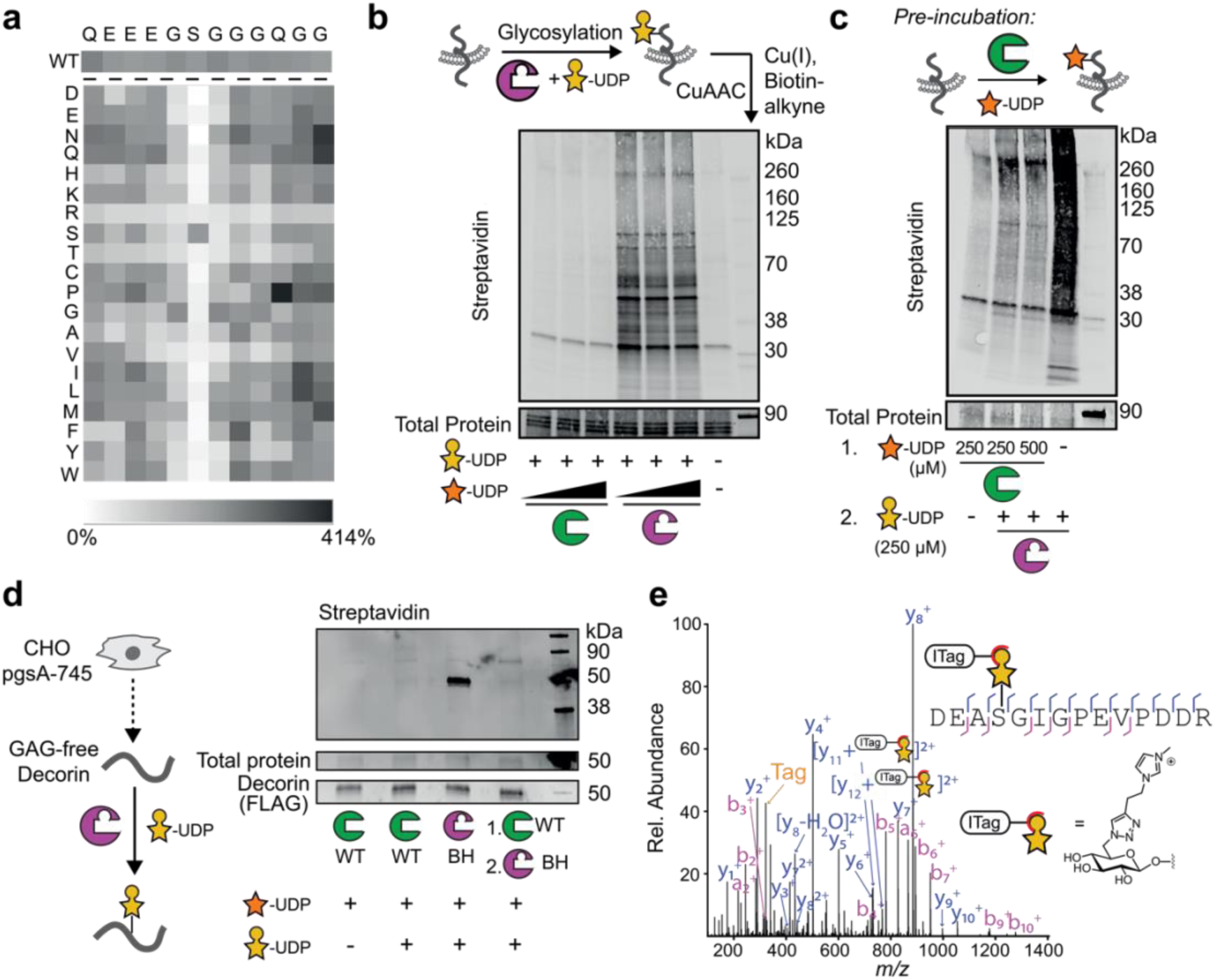
BH engineering preserves protein substrate specificity of XT1. **a**, *in vitro* glycosylation of a peptide substrate panel based on the bikunin peptide indicated at the top of the panel as assessed by a luminescence assay, with every position substituted for each of the 20 amino acids. Intensity of grey scale indicates % turnover. The WT bikunin peptide was present 12 times in the panel (copied in the top row) and all other peptide reactions were normalised on the average of these 12 data points. Data are from one out of three independent experiments. **b**, *in vitro* glycosylation of a membrane protein fraction of XT2-KO CHO^KO^ *^Xyl2t^* cells as assessed by streptavidin blot. Reactions contained 250 µM UDP-6AzGlc and 100, 200 or 300 µM UDP-Xyl, respectively, and were reacted with biotin-alkyne before blotting. Data are from one out of two independent experiments. **c**, glycosylation by BH-XT1/UDP-6AzGlc can be prevented by pre-incubation with WT-XT1/UDP-Xyl. Reactions were processed as in **b**. Data are from one experiment. **d**, *in vitro* glycosylation of a GAG-free preparation of human decorin purified from pgsA-745 CHO cells as assessed by streptavidin blot. Reactions were run with 250 µM UDP-sugars and processed as in **b** and **c**. Data are from one out of two independent experiments. **e**, analysis of the glycosylation site on decorin introduced by BH-XT1 by MS and HCD fragmentation. Decorin was *in vitro* glycosylated as in **c**, subjected to CuAAC with ITag-alkyne,^45^ digested and subjected to MS-glycoproteomics. Fragments are annotated on the tryptic peptide from mature decorin. “Tag” denotes the signature ion 322.1508 *m/z* from the depicted chemically modified sugar. Data are from one experiment.

**Fig. 3:**
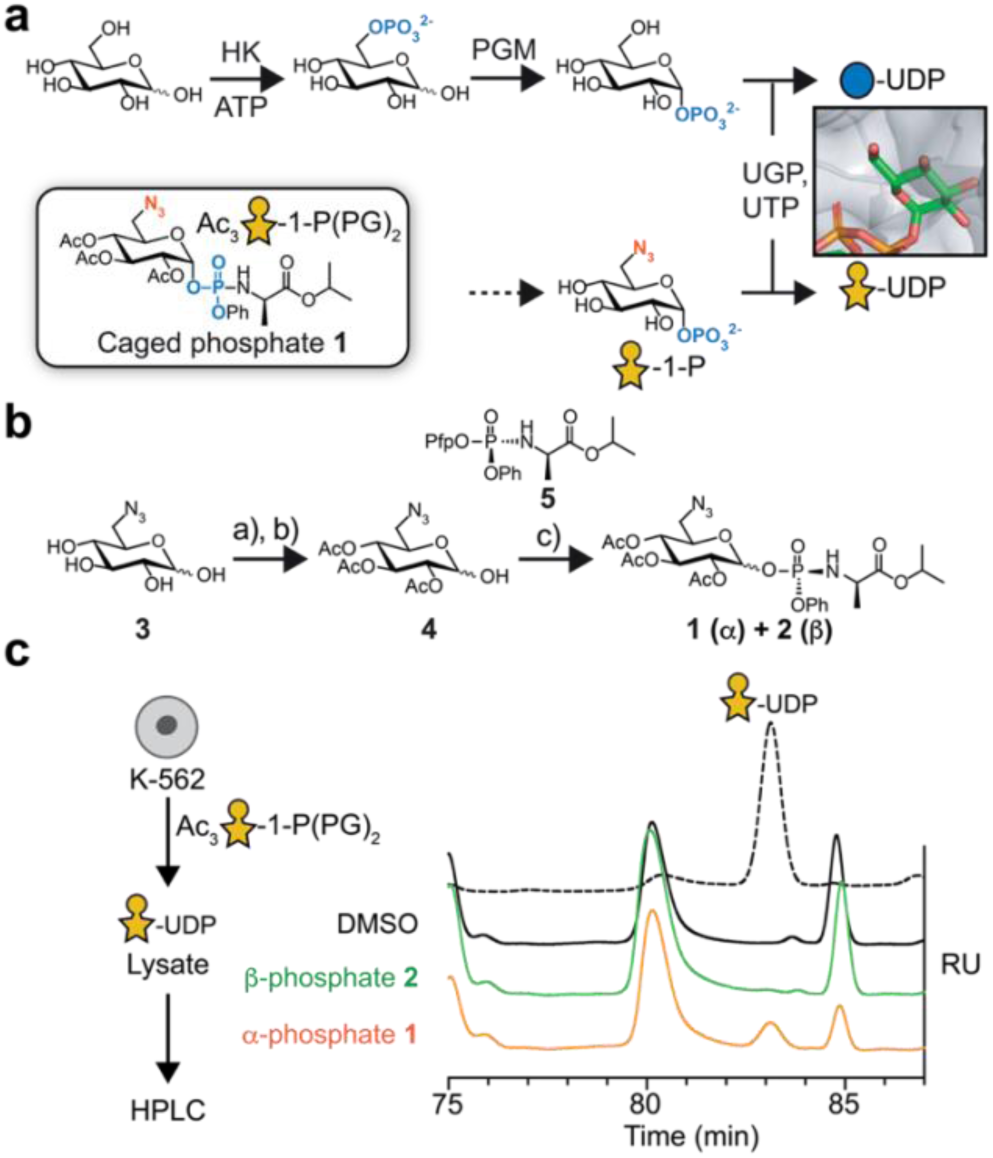
Biosynthesis of UDP-6AzGlc. **a**, biosynthetic pathway of UDP-Glc via hexokinase (HK), phosphoglucomutase (PGM) and UDP-Glc pyrophosphorylase (UGP). Biosynthesis of UDP-6AzGlc from caged phosphate **1** bypasses the HK and PGM steps. Insert: crystal structure of UGP with UDP-Glc indicating that the 6-OH group protrudes into an open cavity (PDB 4R7P). **b**, synthesis of caged sugar-1-phosphates **1** and **2** from 6AzGlc **3**. **c**, biosynthesis of UDP-6AzGlc in K562 cells as assessed by ion exchange HPLC of lysates fed with compounds **1** or **2**. Data are from one out of two independent experiment. Reagents and conditions: a) Ac_2_O, pyridine, DMAP, r.t., 90%; b) AcOH, ethylene diamine, r.t. 45-73%; c) 2 M LDA in THF, **5**, -78°C to -70°C, (**1**) 60% (**2**) 9.8%. DMAP = 4-Dimethylaminopyridine; LDA = lithium diisopropylamide. RU = relative units.

### Bump-and-hole engineering retains the peptide specificity of WT-XT1

To assess whether BH engineering retains substrate preference of WT-XT1 towards the proteoglycan backbone, we first tested the BH enzyme-substrate pair (BH-XT1, UDP-6AzGlc) with a panel of 240 substrate peptides in an *in vitro* glycosylation assay. The panel contained derivatives of the well-characterised bikunin XT1 substrate peptide in which each amino acid was substituted with each of the other 19 proteinogenic amino acids. We had previously used the same peptide panel to extract amino acid preferences of the native enzyme-substrate pair (WT-XT1, UDP-Xyl) in a luminescence-based assay.^17^ Employing the same assay, the peptide substrate preferences were found to be remarkably conserved between WT- and BH-XT1 (Fig. 2a, for WT peptide preference see Briggs and Hohenester^17^). For instance, introducing basic Lys or Arg residues anywhere in the substrate peptide lowered enzyme activity, while acidic Glu and Asp tended to increase activity. An exception was the substitution of Glu at -4 position with Asp that led to a decrease in turnover for both WT- and BH-XT1.^17^ As in WT-XT1, substitutions of glycine residues at positions -1 and +1 of the central Ser were not well-tolerated by BH-XT1. An exception was substitution of Gly at +1 position to hydrophobic amino acids Leu, Met or Phe, which led to residual activity in BH-but not WT-XT1. Since this Gly is in contact with Leu526 in WT-XT1, we reasoned that the Leu526Gly “hole” left space for substitutions to larger hydrophobic amino acids in substrate peptides. Since these were the only reproducible differences between WT and BH-XT1 (other replicates in Supporting Fig. 4) and low in number out of a 240-member peptide library, we concluded that BH engineering exhibits conservation of peptide substrate preference *in vitro*.

**Fig. 4:**
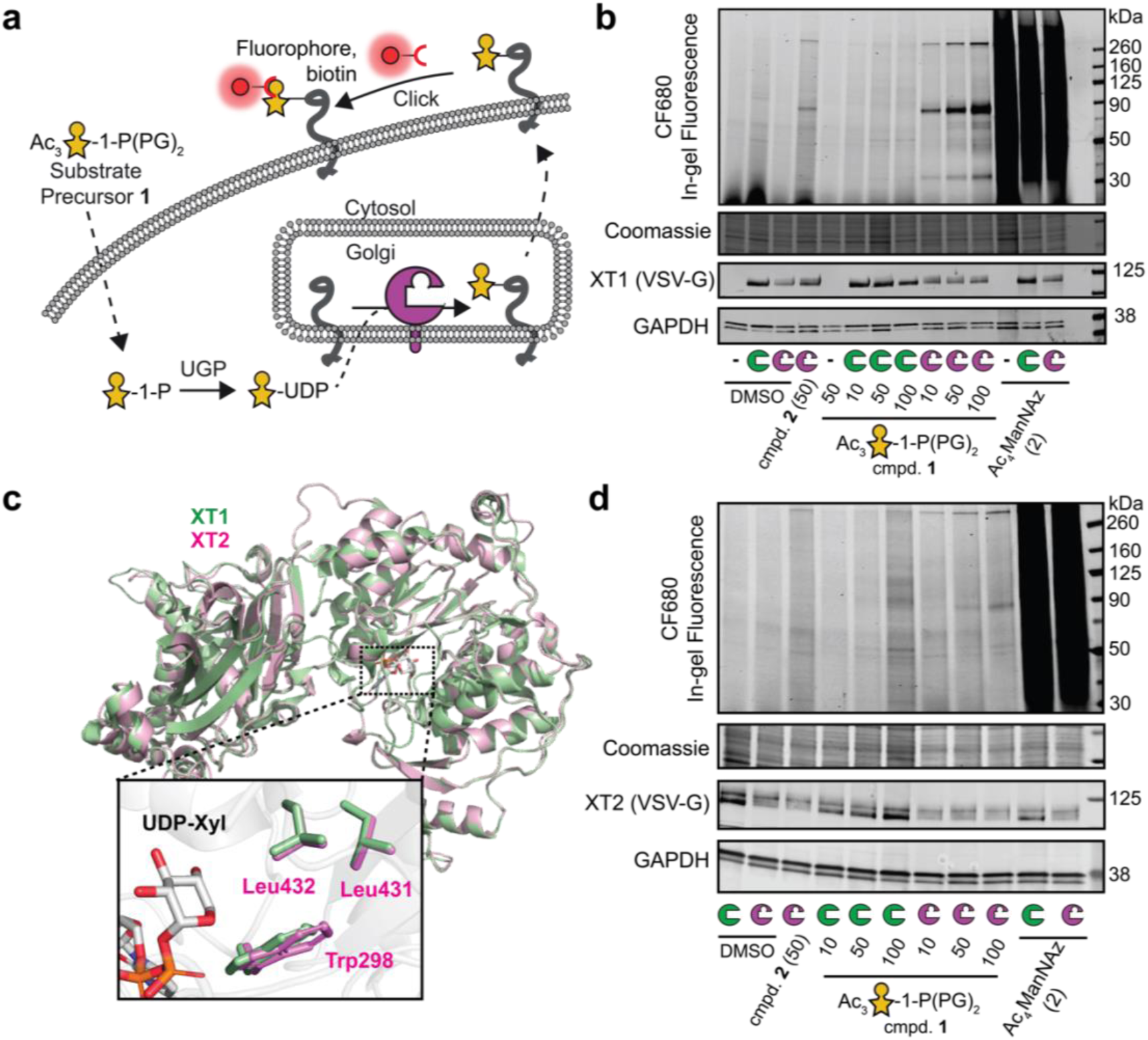
BH-engineered xylosyltransferases label glycoproteins in mammalian cells. **a**, overview of cellular BH engineering. UDP-6AzGlc is biosynthesized after delivery through compound **1**, and used by BH-engineered XTs. Cell surface CuAAC detects proteoglycans. **b**, chemical tagging of proteoglycans by BH-XT1 on pgsA-745 CHO cells as assessed by in-gel fluorescence. Cells stably expressing WT- or BH-XT1 or non-transfected were fed with compounds in the indicated concentrations in µM before on-cell CuAAC and in-gel fluorescence. Western Blots are from a separate gel using the same samples. Data are from one out of two independent replicates. **c**, structural alignment between the crystal structure of human XT1 (PDB 6EJ7, green) and the AlphaFold structure of human XT2 (accession no. Q9H1B5, purple) with aligned gatekeeper residues in the insert. **d**, chemical tagging of proteoglycans by BH-XT2 on pgsA-745 CHO cells as assessed by in-gel fluorescence. Cells stably transfected with WT- or BH-XT2 were fed and treated as in **b**. Data are from one out of two independent experiments.

### BH-XT1 glycosylates proteoglycans at GAG attachment sites *in vitro*

We next assessed whether BH-XT1 retains the activity of WT-XT1 to initiate GAG attachment sites on proteoglycan backbones. We prepared membrane fractions from Chinese Hamster Ovary (CHO) cells with or without KO for endogenous xylosyltransferase genes *Xylt1* and *Xylt2*.^10^ Membrane fractions were incubated with recombinant WT- or BH-XT1 as well as synthetic UDP-Xyl and UDP-6AzGlc, followed by reaction with alkyne-biotin under copper-catalysed azide-alkyne cycloaddtion (CuAAC) “click” conditions. Analysis by streptavidin blot suggested labelling of lysate proteins with 6AzGlc only when BH-XT1, but not WT-XT1 was present (Supporting Fig. 5). Since *Xylt2* is the major xylosyltransferase gene expressed in CHO cells,^10^ we used CHO^KO^ *^Xylt2^* cells for further *in vitro* glycosylation experiments. We established that labelling by BH-XT1 with UDP-6AzGlc could not be outcompeted with increasing concentrations of UDP-Xyl, suggesting that BH-XT1 specifically and potently recognises UDP-6AzGlc as a substrate (Fig. 2b). Pre-incubation of the membrane protein fraction with WT-XT1 and UDP-Xyl abrogated incorporation of 6AzGlc by BH-XT1, suggesting that the same glycosylation sites are introduced by both enzymes (Fig. 2c).

**Fig. 5:**
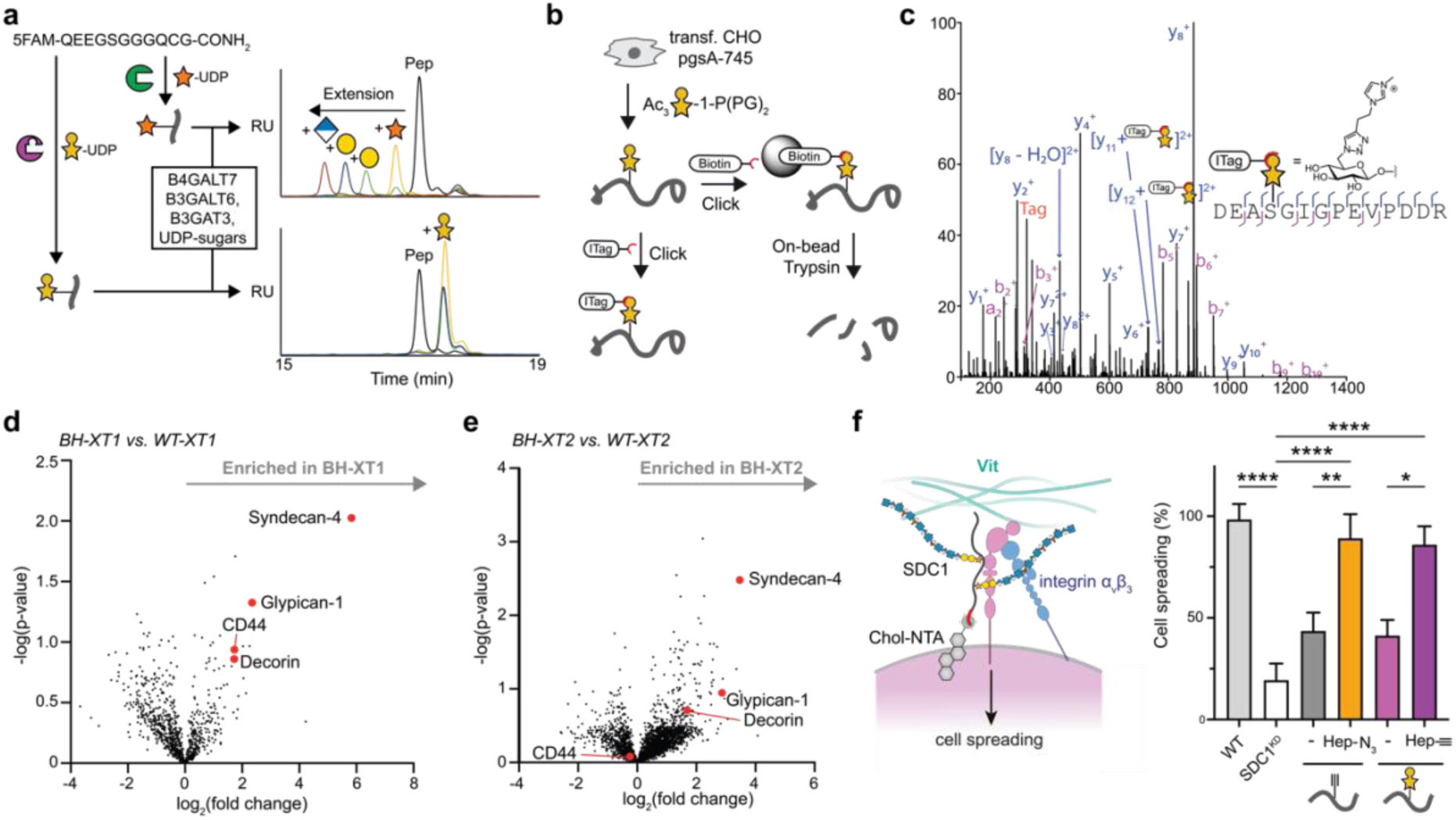
Chemical functionalization by BH-XT1/2 enables detection and manipulation of proteoglycans. **a**, 6AzGlc is not elongated by GAG linker enzymes. Fluorescently labelled bikunin-derived peptide was incubated with WT-XT1/UDP-Xyl or BH-XT1/UDP-6AzGlc and subsequently with the indicated soluble glycosyltransferases^7^ and corresponding UDP-sugars. Elongation was assessed on each step by HPLC. Data are from one experiment each with different combinations of transferases. **b**, using BH-XT1/2 to profile proteoglycans by MS. **c**, analysis of the glycosylation site on decorin introduced by BH-XT1 in living cells by MS and HCD fragmentation. Decorin was co-expressed with BH-XT1 in cells fed with **1**, then subjected to CuAAC with ITag-alkyne,^45^ digested and subjected to tandem MS. Fragments are annotated on the tryptic peptide from mature decorin. Not all fragments are shown. “Tag” denotes the signature ion 322.1508 *m/z* from chemically modified sugar. Data are from one experiment. **d**, differential MS experiment of peptide fractions enriched in BH- vs. WT-XT1-expressing pgsA-745 CHO cells fed with compound **1** from three independent replicates. The proteoglycans were assigned and processed from the *Cricetulus griseus* protein database search (Uniprot) followed by visualization of data on Perseus. **e**, differential MS experiment of peptide fractions enriched in BH- vs. WT-XT2-expressing pgsA-745 CHO cells fed with compound **1** from three independent replicates. **f**, analysis of cell spreading of MDA-MB-231 cells with SDC1-knockdown and rescue with clickable heparin introduced either into alkyne-tagged SDC1_37_ or azide-tagged SDC1-6AzGlc. Spreading relies on the interaction between HS/heparin-containing SDC1 and α_v_β_3_ integrin on MDA-MB-231 cells and vitronectin on coated wells. Statistical significance was assessed by one-way ANOVA, and asterisks denote p values: **** p < 0.0001; ** p < 0.01; * p < 0.05. Data are means + SEM from three independent replicates with cells counted in at least six different frames per replicate.

We next confirmed *in vitro* that BH-XT1 emulates the activity of WT-XT1 to glycosylate proteoglycans. Human decorin has a single site of GAG attachment. Recombinant expression in the CHO cell mutant pgsA-745 that lacks endogenous XT activity results in a GAG-free decorin preparation.^7,8^ We incubated this GAG-free decorin with either WT- or BH-XT1 in the presence of UDP-Xyl and/or UDP-6AzGlc, followed by CuAAC with alkyne-biotin and streptavidin blot. (Fig. 2d). While WT-XT1 activity did not lead to discernible streptavidin signal on decorin, BH-XT1 in the presence of UDP-6-AzGlc led to intense streptavidin signal that could be abrogated by pre-incubation of decorin with WT-XT1/UDP-Xyl. These data indicate that the single GAG attachment site was blocked with a xylose residue by WT-XT1, preventing BH-XT1 activity. We observed the same behaviour in GAG-free preparation of human glypican 1 (Supporting Fig. 6), suggesting that BH-XT1 recapitulates the activity of WT-XT1 across a range of proteoglycans.

We confirmed the glycosylation site modified by BH-XT1 on *in vitro* glycosylated recombinant decorin by tandem MS. Two fragmentation methods are routinely employed for O-glycopeptides. High energy collision-induced dissociation (HCD) primarily fragments the glycosidic bond to detect glycan oxonium ions while electron transfer dissociation (ETD) fragments the peptide backbone to allow glycan site annotation. The clickable azide tag was essential to improve sugar identification in mass spectra, allowing incorporation of functional groups that are beneficial to analysis. Specifically, we employed a clickable imidazolium tag (ITag) that carries a permanent positive charge and increased the charge state of glycopeptides, allowing direct glycosylation site annotation.^45^ We first employed a standard workflow in which HCD fragmentation led to an ITag-containing, 6AzGlc-derived signature ion that was used to trigger ETD on the same glycopeptide.^35,41,43,45^ This tandem strategy is used because O-linked glycan modifications are usually too labile to be detected within peptide fragments during HCD, hampering glycosylation site localisation. Surprisingly, ITag- modified 6AzGlc was detected on peptide fragments in HCD spectra on a tryptic glycopeptide derived from decorin, without the need for additional ETD (Fig. 2e).

Decorin is proteolytically processed during secretion to remove a propeptide and shorten the N-terminus.^46^ HCD fragmentation allowed for direct identification of Ser34 as the attachment site of 6AzGlc by BH-XT1 on this mature form, consistent with Ser34 being the site of cellular GAG attachment (Fig. 2e).^47^ Taken together, these results suggest that the XT1 bump-and-hole enzyme-substrate pair glycosylates native GAG attachment sites in proteoglycans *in vitro*. We also surmise that the 6AzGlc modification is stable towards HCD fragmentation, allowing for facile annotation of GAG attachment sites by MS.

### Biosynthesis of UDP-6AzGlc in mammalian cells via a caged sugar-1-phosphate

Application of a GT BH system in living cells requires biosynthesis of the nucleotide-sugar. In general, caged, membrane-permeable monosaccharide precursors are employed with ester modifications that are deprotected in the cytosol. Free monosaccharides can then be converted to UDP-sugars before transport to the Golgi.^42,43,48^ Although human cells are devoid of a salvage pathway for UDP-Xyl, our strategic use of UDP-6AzGlc provided an opportunity for cellular application by hijacking the biosynthetic pathway for UDP-Glc instead. Glc is activated in mammalian cells first by phosphorylation to Glc-6-phosphate and, subsequently, isomerization by phosphoglucomutase PGM to Glc-1-phosphate (Fig. 3a). Conversion to UDP-Glc then features the enzymes UDP-Glc pyrophosphorylase 1 or 2, UGP1/2. Since phosphorylation at the 6-position was prevented by the presence of an azido group, we sought to bypass the kinase and PGM steps and provide a sugar-1-phosphate as a direct substrate for UGP1/2. We were encouraged by analysis of the UGP1/UDP-Glc co-crystal structure in which the UDP-Glc 6-hydroxyl group is solvent-exposed, suggesting that an azido group at that position should be tolerated by the enzyme (Fig. 3a).^49^ While we and others have made sugar-1-phosphates caged as labile bis-*S*-acetylthioethyl (SATE) phosphotriesters, synthesis of SATE-caged 6AzGlc-1-phosphate failed in our hands.^43,50,51^ Instead, we took inspiration from the increasingly popular protide technology that has gained attention to cage phosphates in antiviral nucleotides,^52,53^ and has been recently used to cage sugar-phosphates.^54–56^ We synthesized phosphoramidate diester **1** as a caged sugar-1-phosphate to be deprotected by hydrolases in the cytosol of living mammalian cells (Fig. 3a).^52^ The synthesis proceeded from lactol mixture **3** via the intermediate triacetate **4.** Treatment of **4** with **5** under basic conditions yielded both α-phosphate **1** (60% yield) and β-phosphate **2** (9.8% yield) (Fig. 3b).^57^ Since UGP1/2 is naturally restricted to α-configured Glc-1-phosphate, preferential formation of the α-anomer was gratifying. In turn, β-phosphate **2** served as a negative control in feeding experiments. Feeding K-562 cells the α-phosphate **1** led to notable and reproducible biosynthesis of UDP-6AzGlc (Fig. 3c). In turn, β-configured phosphate **2** led to negligible UDP-6AzGlc levels, possibly arising from very small (4%) amounts of α-configured phosphate **1** as a contaminant of **2** (Supporting Information). These data suggest that α-phosphate **1** is a suitable precursor to deliver UDP-6AzGlc to mammalian cells by entering the UDP-Glc biosynthetic pathway.

### Development of a cellular XT1 bump-and-hole system

With a strategy for UDP-6AzGlc delivery in hand, we established a BH system to chemically tag XT1 substrate proteins in mammalian cells (Fig. 4a). The pgsA-745 CHO cell line was employed because of the lack of competition with internal xylosyltransferases. Cells were stably transfected with plasmids encoding WT- or BH-XT1 using a transposase-based genome insertion method,^43,58^ followed by feeding the 6AzGlc-1-phosphate precursor **1**. After overnight incubation, CuAAC was performed to attach clickable alkyne-CF680 to 6AzGlc on the cell surface while keeping cells alive.^35,43,48^ Surplus click reagents were washed away, cells lysed and fluorophore incorporation assessed by SDS-PAGE and in-gel fluorescence (Fig. 4b). Minimal background fluorescence was observed in cells fed with DMSO or only expressing WT-XT1, even when fed with increasing concentrations of 6AzGlc-1-phosphate precursor **1**. In the presence of BH-XT1, clear fluorescent proteins were observed at 38 kDa, 90 kA and 260 kDa. With increasing feeding concentration of **1**, a dose-dependent increase of fluorescence was observed, along with labelled protein bands of lower intensity, especially between 50 and 90 kDa. In accordance with biosynthetic experiments, the β-configured 6AzGlc-1-phosphate **2** yielded a weak and diffuse labelling signal when fed to cells, indicating that UDP-6AzGlc biosynthesis is a direct prerequisite for cellular chemical tagging of glycoproteins by BH-XT1. The fluorescent bands at 90 kDa and 260 kDa in the BH-XT1/compound **1** lanes were observed weakly when compound **2** was fed, which we attribute to small residual levels of compound **1** in the preparation (*see* Fig. 3c). Ac_4_ManNAz as a precursor to azide-tagged sialic acid yielded a strong and uniform fluorescent signal across all cell lines tested.^59^ Our data indicate that BH-XT1 specifically tags cell surface proteins with bioorthogonal 6AzGlc in the secretory pathway of mammalian cells.

The mammalian genome encodes two xylosyltransferase isoenzymes that have been differentially implicated in disease and proteoglycan function.^28,30,32,60^ To allow for substrate profiling of the second isoenzyme XT2, we extended the BH strategy by employing structural similarities between the two isoenzymes. A structural overlay between the XT1 crystal structure and the XT2 structure predicted by AlphaFold highlighted conservation of amino acids interacting with UDP-Xyl (Fig. 4d). We identified the gatekeeper residues Leu431, Leu432 and Trp298 in XT2 that occupied the same role as the respective residues in XT1, with Leu432 in XT2 overlaying with Leu526 in XT1. The BH-XT2 mutant L432G was thus stably expressed in pgsA-745 CHO cells. Feeding with 6AzGlc-1-phosphate precursor **1** and cell-surface CuAAC reaction with CF680-alkyne led to a similar band pattern by in-gel fluorescence, with glycoprotein bands at 90 kDa and 260 kDa identified in BH-XT2-expressing cells in a dose-dependent manner (Fig. 4d). We noted that the intensity of fluorescent bands specifically tagged by BH-XT2 was lower than by BH-XT1, indicating a lower activity of the second isoenzyme. WT-XT2 expression did not lead to the same fluorescent band pattern. Neither feeding DMSO nor compound **2** led to discernible signal over background, and Ac_4_ManNAz was included as a positive labelling control. Our data suggest that the BH approach is applicable to both xylosyltransferase isoenzymes in mammalian cells, allowing for the first time to assess their substrate profiles.

### XT bump-and-hole engineering enables profiling of cellular proteoglycans

Xylosyltransferase BH engineering is poised to allow the identification of proteoglycans, a feat that normally requires elaborate methods of glycopeptide enrichment and characterisation.^9,21,23^ A pre-requisite to glycoprotein enrichment and identification is biosynthetic simplicity – ideally, 6AzGlc would replace an entire GAG chain without added complexity from glycan elaboration. To establish an MS-glycoproteomics workflow, it was thus important to assess whether 6AzGlc, like Xyl, was extended to a functional GAG linker tetrasaccharide. We recently reported an enzymatic method for extension of xylosylated glycopeptides by recombinant preparations of the glycosyltransferases B4GALT7, B3GALT6 and B3GAT3 (termed linker enzymes) in the presence of UDP-galactose (UDP-Gal) and UDP-glucuronic acid (UDP-GlcA). Employing a fluorescently labelled bikunin-derived peptide, we first confirmed by HPLC that attachment of either Xyl (by WT-XT1) or 6AzGlc (by BH-XT1) led to a shift of peptide retention time (Fig. 5a). Upon addition of the linker enzymes, the Xyl moiety was sequentially elongated to the full tetrasaccharide. In contrast, a 6AzGlc-modified peptide did not shift in retention time upon incubation with the GTs and UDP-sugars. We concluded that 6AzGlc is a chain-terminating modification that is not extended to functional GAG chains. We interpreted this substantial decrease in glycan complexity as an advantage for MS and to manipulate the composition of proteoglycans (see below).

We next determined that a functional XT1 system chemically tags a model proteoglycan in mammalian cells by MS. FLAG-tagged human decorin was overexpressed as a secreted proteoglycan in pgsA-745 CHO cells that expressed BH-XT1. Cells were fed with 6AzGlc-1-phosphate precursor **1**, and decorin immunoprecipitated from conditioned supernatant. The preparation was subjected to CuAAC with ITag-alkyne, digested and subjected to MS with HCD fragmentation. We confirmed unambiguously that Ser34 was glycosylated by BH-XT1 inside mammalian cells, confirming the bump-and-hole approach as suitable to identify proteoglycans including native Xyl attachment sites (Fig. 5b, c).

Our in-gel fluorescence data suggested that the two isoenzymes XT1 and XT2 may exhibit an overlapping substrate glycoprotein profile. To test this notion, we compared the glycoproteome tagged by either BH-XT1 or BH-XT2 in cells by MS. Chemically tagged glycoproteins in conditioned supernatants of stably transfected pgsA-745 CHO cells were treated with alkyne-biotin, enriched on neutravidin resin and subjected to on-bead proteolytic digest. Peptide fractions were analysed by MS using data-independent acquisition over three replicates each, with WT-XT1/2-expressing cells fed with **1** as control conditions. Data were processed using the software DIA-NN to identify and quantify *Cricetulus griseus* proteins.^61^ Enriched peptide and protein fractions were analyzed with Perseus.^62^ Glycosylation by BH-XT1 led to a striking enrichment of cellular proteoglycans, including decorin, syndecan-4, glypican-1 and CD44, compared to the supernatant of cells expressing WT-XT1 (Fig. 5d, Supporting Fig. 7). BH-XT2 led to strong enrichment of the same glycoproteins with exception of CD44 (Fig. 5e, Supporting Fig. 7). In both cases, we analyzed the identity of detected peptides from these enriched proteins and found them to lie outside previously annotated glycosylation site(s) (Supporting Table 1). These findings suggests that cellular XT1/2 BH engineering allows unambiguous proteoglycan profiling without the need to annotate the GAG attachment site. When a less stringent comparison was made between cells expressing BH-XT1/2 fed with compound **1** against the same cell lines fed with DMSO as a control, the *bona fide* proteoglycans nidogen-1, chondroitin sulfate proteoglycan 4, agrin and aggrecan were additionally found to be enriched (Supporting Fig. 8). We conclude that BH engineering enables straightforward detectability of proteoglycans from mammalian cells.

### Chemical functionalization allows for functional annotation of the GAG modification on proteoglycans

The BH approach replaces the GAG chain of a proteoglycan with a bioorthogonal modification. We reasoned that this approach could be used to introduce suitably modified GAG polysaccharides, creating neo-glycoproteins that allow for functional dissection of the proteoglycan components in cell biology. We have previously used unnatural amino acid introduction via amber stop codon reassignment to produce syndecan-1 (SDC1) chemically tagged at position 37 (termed SDC1_37_) with a clickable alkyne.^16^ Through CuAAC, SDC_37_ could be furnished with azide-containing, clickable heparin to generate a neo-proteoglycan termed SDC1_37_-Hep. In the presence of either fully glycosylated, HS-containing SDC1 or SDC1_37_-Hep, MDA-MB-231 cells exhibit enhanced spreading on vitronectin-coated surfaces in an integrin α_v_β_3_-dependent fashion that can be analyzed by microscopy. Soluble SDC1 preparations can be deposited on cells by virtue of their hexa-His tags through the use of synthetic Ni^2+^-displaying, membrane-anchored cholesterol anchor.^16^ We used BH-XT1 to introduce 6AzGlc into a non-glycosylated WT-SDC1 preparation *in vitro*, to generate SDC1-6AzGlc. Heparin was furnished with an anomeric alkyne tag via hydrazide chemistry, and introduced into 6AzGlc-SDC1 via CuAAC to furnish SDC1-6AzGlc-Hep. Both SDC1-Hep conjugates displayed an increase in molecular weight (Supporting Fig. 9). MDA-MB-231 cells were depleted of endogenous SDC1 by siRNA-based knockdown, significantly impacting cell spreading (Fig. 5f). Deposition of non-glycosylated SDC1_37_ or SDC1-6AzGlc enhanced cell spreading by approx. 20%, suggesting a role of the SDC backbone alone for MDA-MB-231 spreading that has been noted before.^16^ Heparin-containing SDC1_37_-Hep and SDC1-6AzGlc-Hep fully rescued cell spreading, indicating that BH engineering and bioorthogonal chemistry restored the functional properties of a proteoglycan.

## Discussion

The importance of proteoglycans in physiology is undisputed, as the vast majority of signalling events between cells or with the extracellular matrix are modulated by the associated GAG chains. While great efforts are being made to understand the details of GAG polysaccharide sequence on biology,^10,14,22,63,64^ we still lack important information on the first step of glycosylation to the protein backbone. The two human xylosyltransferases display tissue selectivity and differences in attached GAG sequences but we do not yet have fundamental detail on their individual biological functions.^28,31–33,60^ Furthermore, only a relatively small number of annotated mammalian proteoglycans has been identified, with new annotations requiring considerable effort.^9^ A chemical tool to dissect XT1/2 biology must accurately report on XT1/2 activity while being orthogonal to other glycosylation events in the secretory pathway and deliverable to living mammalian cells. The use of a UDP-Glc analogue matched these pre-requisites. Both catalytic efficiency and peptide substrate preference of the XT1 bump-and-hole enzyme-substrate pair were remarkably conserved, which we attribute to the careful structure-based design of the pair. Incorporation of 6AzGlc into cell surface proteins was dependent on the presence of BH-XT1, indicating that the sugar does not enter other major glycosylation pathways. We note that this finding does not exclude incorporation into glycoconjugates that naturally contain D-Glc, as most of these are either glycolipids or intracellular glycoproteins, neither of which would be visible with our assays.

The finding that 6AzGlc is not extended to the common GAG linker tetrasaccharide was expected due to the restrictive active site architecture of the follow-on enzyme B4GALT7.^65^ However, we recognize this non-extension as a particular advantage for MS since the sugar 6AzGlc is structurally well-defined and does not require any further simplification by glycosidase treatment prior to analysis. In our MS experiments, annotation was further simplified by the availability of the ITag technology to facilitate MS.^45,66^ Furthermore, a chain-terminating, clickable inhibitor of chain extension has the potential to be employed to study GAG biology *in vitro* or *in vivo*,^16,39,67^ substantially expanding our toolbox.

Establishing a cellular XT bump-and-hole system required a biosynthetic entry point for UDP-6AzGlc. Previously used per-acetylated 6AzGlc was not a suitable precursor for glycosylation in our hands,^36^ but we note that cell lines from different organisms can vary in their biosynthetic potential.^68^ Nevertheless, a protide-based caged sugar-1-phosphate was a reliable precursor for UDP-6AzGlc to fashion a cellular bump-and-hole system. We did not assess whether UDP-6AzGlc was used by other glycosyltransferases, but cell surface labelling experiments suggested that BH-XT activity was necessary to introduce the chemical modification into glycoproteins.

While XT2 appears to be the dominant isoenzyme expressed in humans, dysfunctions in both enzymes lead to severe yet differential disorders in mouse models and in patients.^30–33^ After fully characterising an XT1 BH system, we designed a functional BH-XT2 mutant simply based on structural homology. BH-XT2 was directly applied in living cells without characterizing the corresponding soluble recombinant enzyme first, showcasing the reliability of the tactic as well as the importance of structural data for identification of nucleotide-sugar binding.

Proteoglycans can be perceived as a modular assembly between a protein backbone and one or more GAG chains. While biological function can be imparted by either component, it can be challenging to differentiate both. The biosynthetic details of GAG extension by either HS or CS/DS are beginning to be understood,^7^ but methods to reliably swap the GAG chain on a given proteoglycan are of fundamental importance for proteoglycan biology. We have recently employed stop codon reassignment to introduce an alkyne-tagged amino acid into a recombinant proteoglycan backbone,^16^ as a critical aspect of understanding the role of SDC1 for cell spreading. We applied BH engineering to underpin these findings, allowing to chemoenzymatically attach a chemical modification to the recombinant protein. We note the ability to employ this modular approach to generate “designer proteoglycans” with unnatural GAG chains to generate functional understanding.

Our work is setting the foundation to establish the fine differences between XT1 and XT2 and profile proteoglycans in a range of different model systems.

## Acknowledgements

We thank Ganka Bineva-Todd and Andrea Marchesi for help with chemical synthesis and analysis, and Khadija Babiker for help with cloning similar constructs. We thank Chloë Roustan, Svend Kjaer and the Francis Crick Institute Structural Biology Science Technology Platform for help with protein expression and purification. We further thank Tania Auchynnikava and Mark Skehel for help with mass spectrometry, and the Crick Proteomics, Chemical Biology and Cell Services Science Technology Platforms for valuable support. This work was supported by the Francis Crick Institute (to B. S.) which receives its core funding from Cancer Research UK (CC2127, CC2068), the UK Medical Research Council (CC2127, CC2068) and Wellcome Trust (CC2127, CC2068), the BBSRC (BB/T01279X/1 to E. H. and B. S., BB/V008439/1 to B. S.), the EPSRC (EP/T007397/1 to G.J.M) and the MRC (MR/T019522/1 to G.J.M). This work was supported in part by the NIH under award R35GM142462 (to M. L. H.). We also thank the EPSRC UK National Mass Spectrometry Facility (NMSF) at Swansea University. This work was supported by the Novo Nordisk Foundation grant No. NNF22OC0073736 (to R. L. M.). L. I. W. gratefully acknowledges funding from the European Research Council (ERC) under the European Union’s Horizon 2020 research and innovation programme (Grant agreement No 851448). For the purpose of Open Access, the author has applied a CC BY public copyright licence to any Author Accepted Manuscript version arising from this submission.

## Notes

### Competing Interest Statement

The authors have declared no competing interest.

### Summary of Updates

MS-proteomics and functional application data added.

