## Supporting Information for "Xylosyltransferase Bump-and-hole Engineering to Chemically Manipulate Proteoglycans in Mammalian Cells"

[a] Department of Chemistry, Imperial College London, W12 0BZ, London, United Kingdom.
[b] Chemical Glycobiology Laboratory, The Francis Crick Institute, NW1 1AT London, United Kingdom
[c] Proteomics Science Technology Platform, The Francis Crick Institute, London NW1 1AT, United Kingdom.
[d] Lennard-Jones Laboratory, School of Chemical & Physical Sciences, and Centre for Glycoscience, Keele University, Keele, Staffordshire, ST5 5BG United Kingdom.
[e] Skaggs Graduate School of Chemical and Biological Sciences, Scripps Research, La Jolla, California.
[f] Department of Chemistry, Scripps Research, La Jolla, California.
[g] Department of Life Sciences, Imperial College London, London SW7 2AZ, UK.
[h] Signalling and Structural Biology Laboratory, The Francis Crick Institute, London NW1 1AT, United Kingdom.
[i] Ddepartment of Chemistry, Yale University, CT 06511, New Haven, United States.
[j] York Structural Biology Laboratory and York Biomedical Research Institute, Department of Chemistry, University of York, York YO10 5DD, United Kingdom
[k] Copenhagen Center for Glycomics, Department of Cellular and Molecular Medicine, Faculty of Health Sciences, University of Copenhagen; Copenhagen-N, Denmark.
[%] Current address: Avacta, London, United Kingdom.
[&] Current address: GlycoDisplay ApS, Copenhagen, Denmark.
[#] These authors contributed equally.

#### Supporting Figures

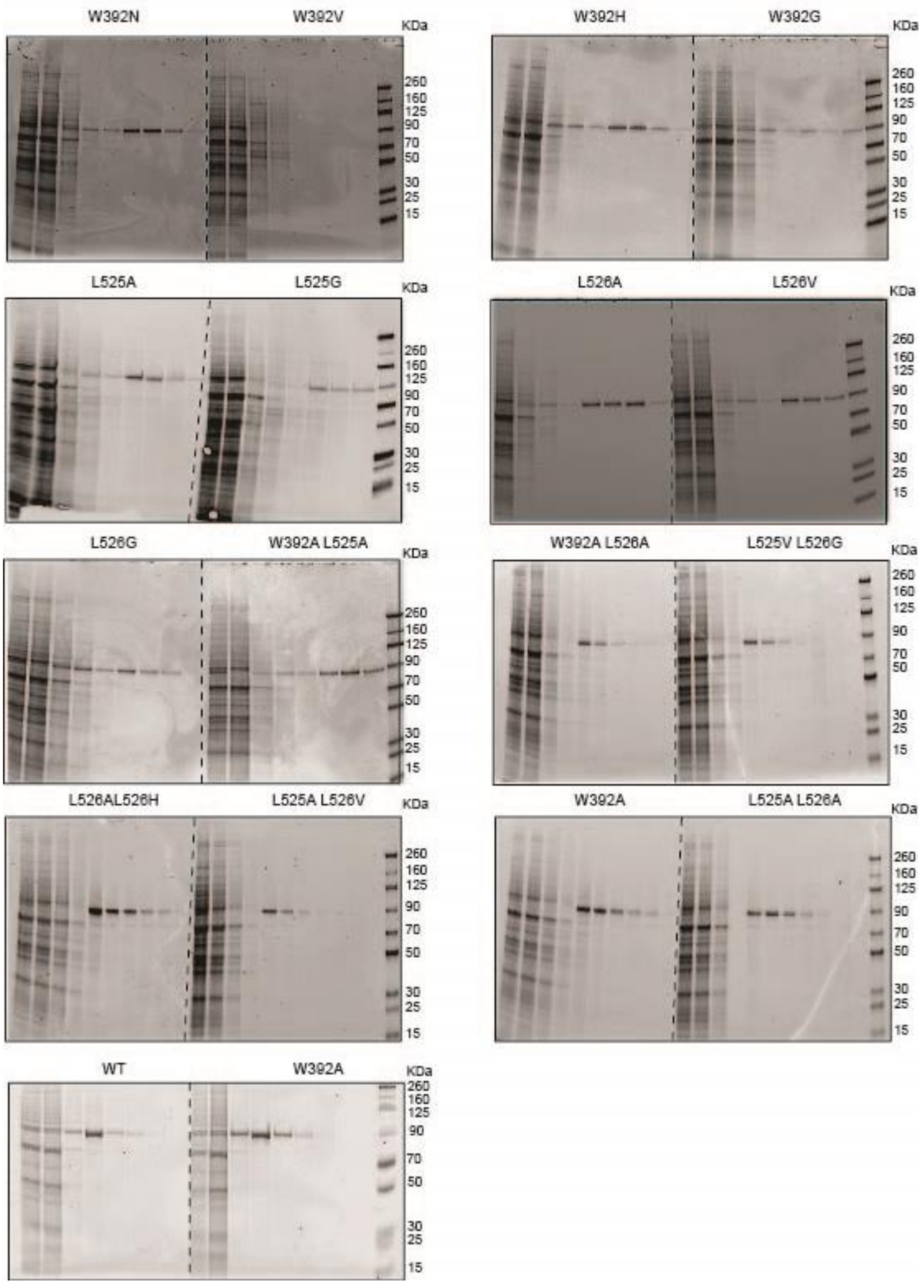


**Supporting Fig. 1:** Expression of XT1 constructs in Expi293F cells on 10 mL scale purified by Ni-NTA affinity chromatography and visualised by SDS-PAGE and Coomassie staining. Fractions are from left to right: Lane 1 Input, 2 flow-through, 3-5 20 mM imidazole, 6-7 50 mM imidazolde, 8-9 200 mM imidazole. Note that the final fraction was omitted for some constructs.

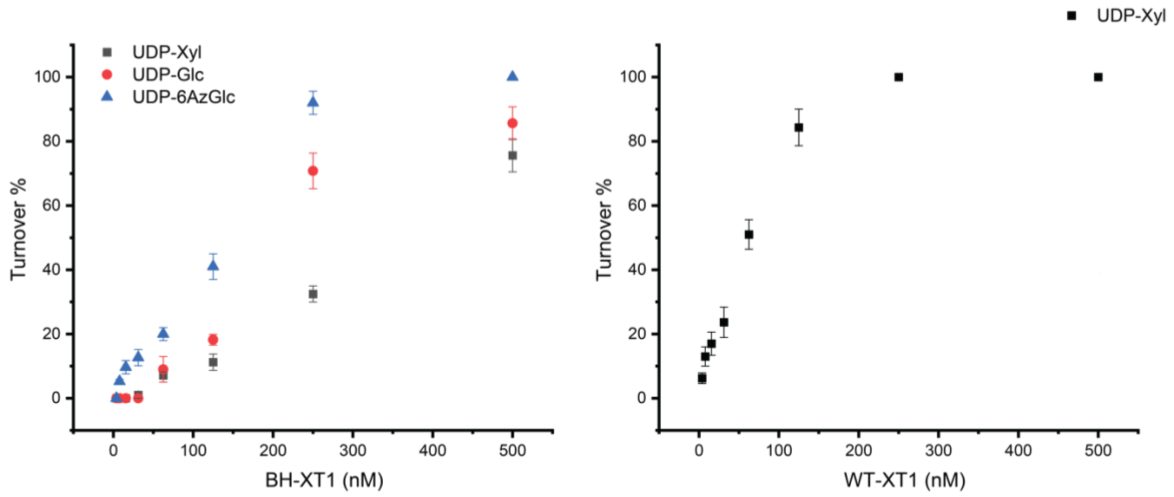


**Supporting Fig. 2:** Enzyme dependence of XT1 *in vitro* glycosylation reactions. Turnover of fluorescent bikunin substrate peptide was assessed in reactions containing constant concentrations of peptide and UDP-sugar. Enzyme concentration was varied. Analysis was performed by HPLC and plots were used to determine the enzyme concentration required to achieve 10-20% conversion. Data are means ± SD of three technical replicates.

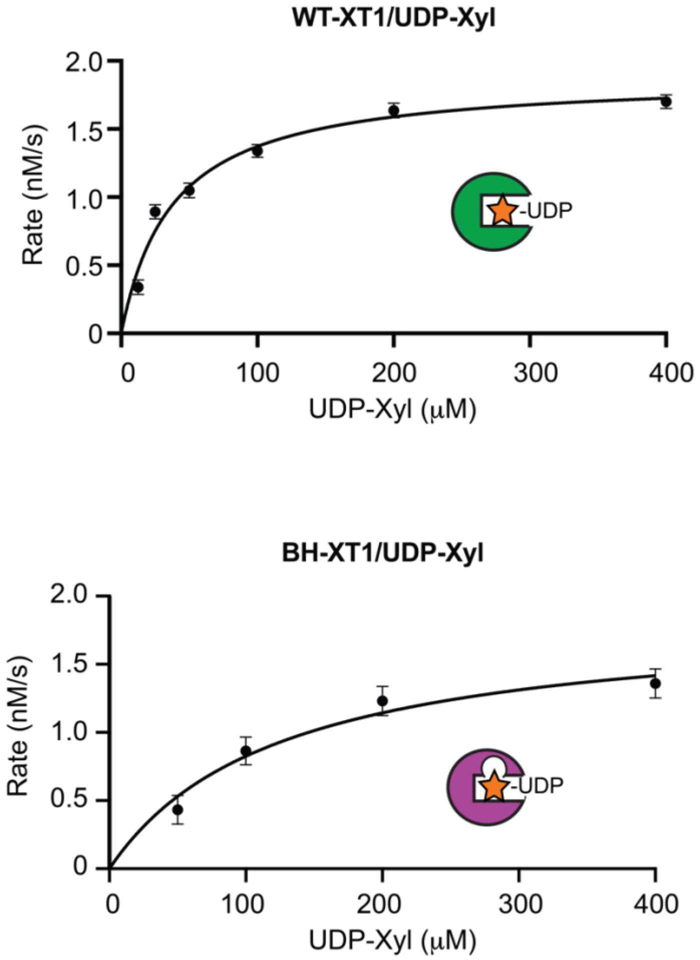


**Supporting Fig. 3:** Michaelis-Menten kinetics of the indicated enzyme-substrate pairs. Data are means ± SD of three technical replicates.

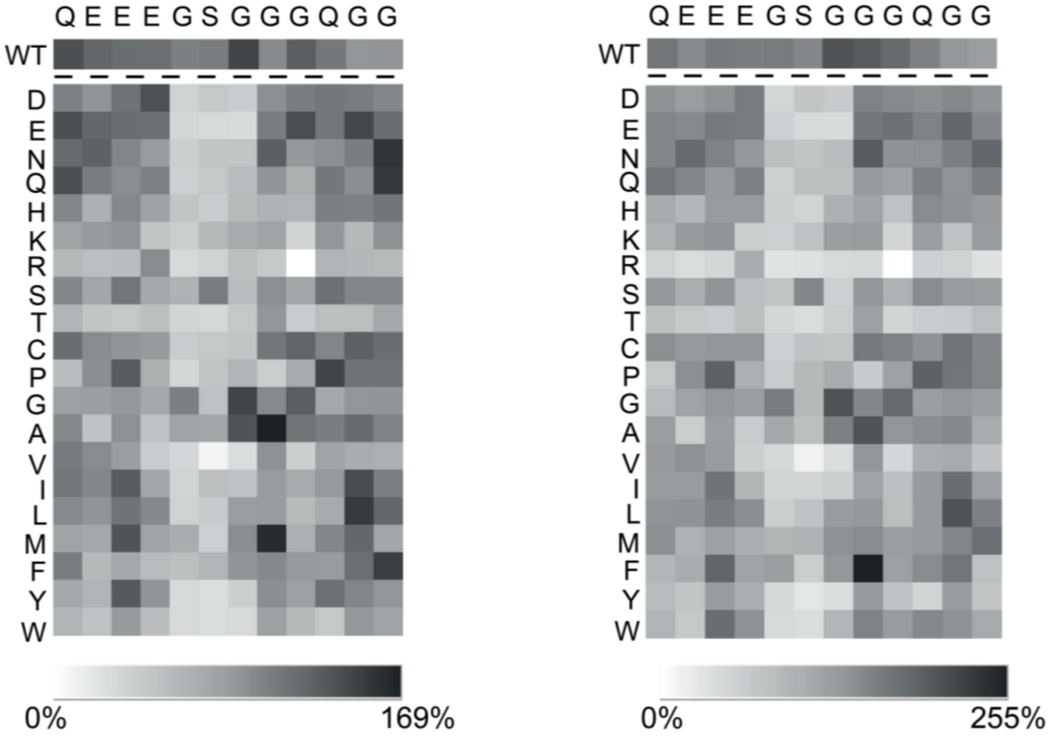


**Supporting Fig. 4**: Independent replicates of *in vitro* glycosylation with the bikunin peptide substrate panel. Data were generated as in Fig. 2a.

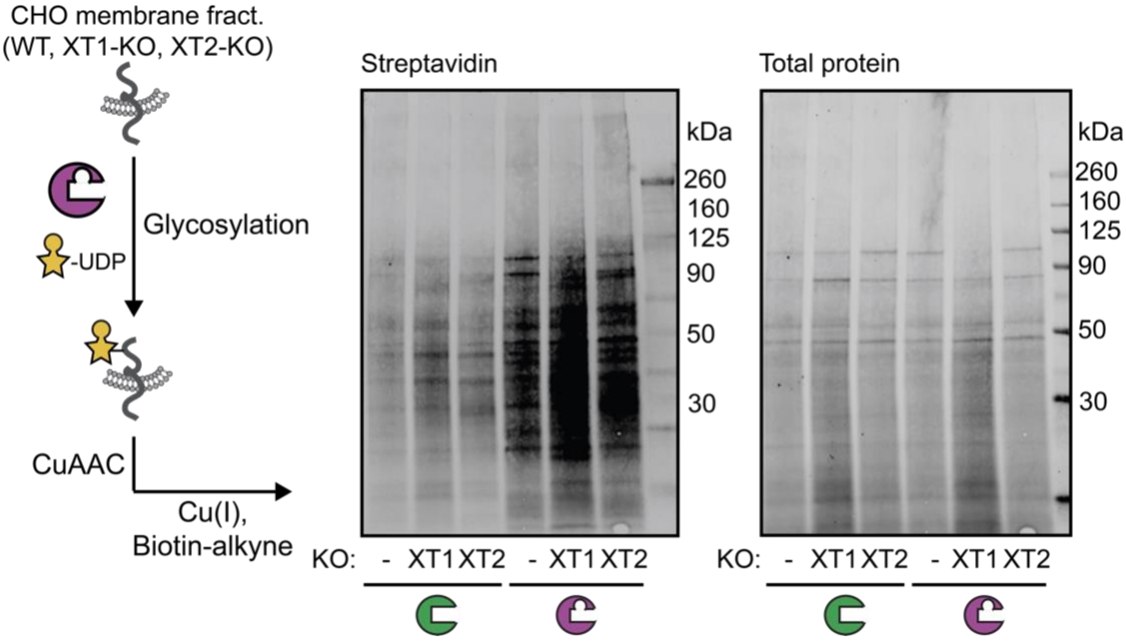


**Supporting Fig. 5**: *in vitro* glycosylation of a membrane protein preparation of parental, XT1-KO or XT2-KO CHO cells as assessed by streptavidin blot. Reactions contained 250 µM UDP-6AzGlc and were reacted with biotin-alkyne before blotting as in Fig. 2b. Data are from one experiment.

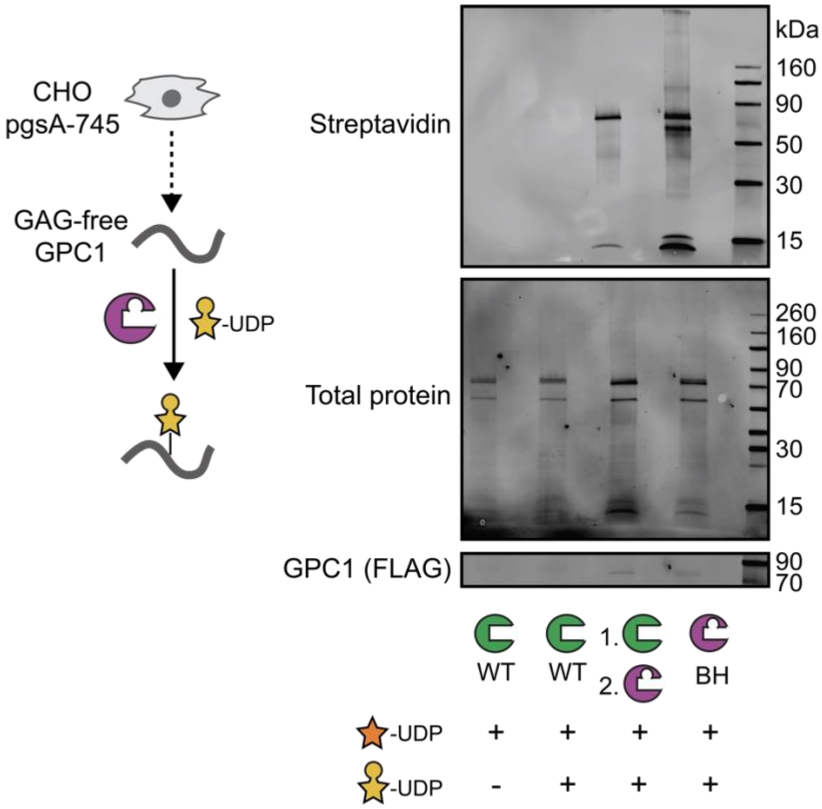


**Supporting Fig. 6**: *in vitro* glycosylation of a GAG-free preparation of FLAG-tagged human glypican-1 from pgsA-745 CHO cells as assessed by streptavidin blot. Reactions contained 250 µM UDP-sugars and were processed with or without pre-incubation with WT-XT1/UDP-Xyl as indicated. Data are from one out of two independent replicates.

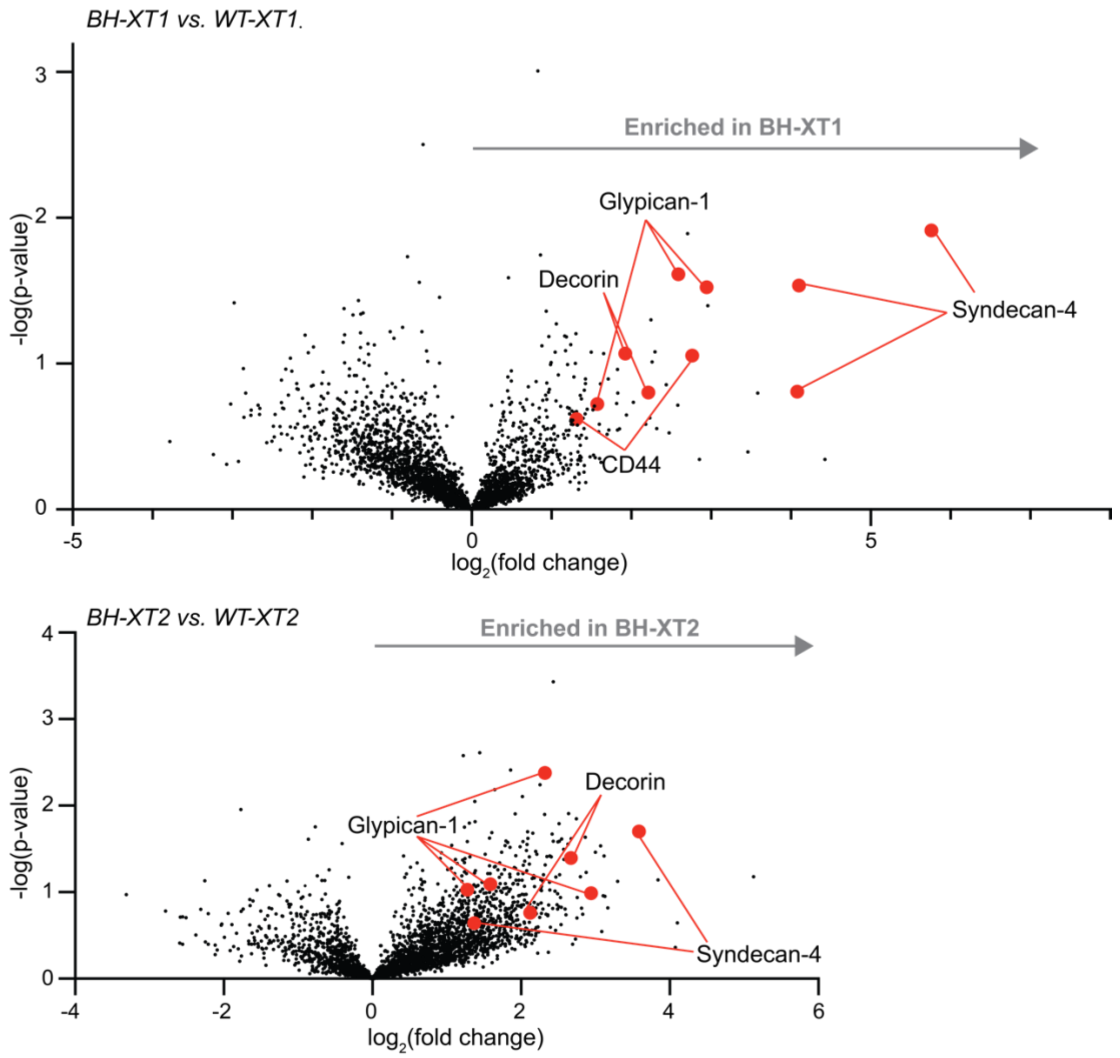


**Supporting Fig. 7:** Volcano plots of individual peptides enriched by in differential MS experiments of secretome from BH- vs. WT-XT1/2-expressing pgsA-745 cells fed with compound **1** from three independent replicates.

**Supporting Table 1:** Identified, enriched peptides from secretome samples in BH-XT1 and BH-XT2. Samples were processed according to Fig. 5d/e and analysis performed in Perseus according to Supporting Fig. 7. Only one charge state per peptide is shown. n/a = not analyzed.

| | | BH-XT1 vs. WT-XT1 | | | BH-XT2 vs. WT-XT2 | | |
|---|---|---|---|---|---|---|---|
| | **Peptide sequence** | **charge** | **-log(p value)** | **log2(fold change)** | **charge** | **-log(p value)** | **log2(fold change)** |
| **Syndecan-4** | EVEQNEVIPK | +2 | 1.9132256 | 5.76043749 | +2 | 1.70032027 | 3.58913064 |
| | VSPSEVDNDISNK | +2 | 0.80799931 | 4.0779624 | +2 | 0.64023339 | 1.36967818 |
| | RVSPSEVDNDISNK | +3 | 1.53655427 | 4.10091321 | +3 | 0.59440457 | 1.77871513 |
| | ETEVIDHQHFLEGR | +3 | 0.5158269 | 1.69561354 | +3 | 0.35888786 | 0.94391918 |
| **Glypican-1** | LALQEKPPTGSLEK | +3 | 0.28764139 | 1.1448005 | +3 | 2.37743205 | 2.32337936 |
| | SFVQGLGVASDVVR | +2 | 1.61215323 | 2.58823967 | +2 | 1.09004828 | 1.58631134 |
| | GCLANQADLDAEWR | +2 | 0.60544997 | 1.24314292 | +2 | 1.02892665 | 1.27857939 |
| | VAQVPLAPECSR | +2 | 1.5246964 | 2.94538943 | +2 | 0.98795528 | 2.9424003 |
| **Decorin** | NLHTLILVNNK | +3 | 0.79836121 | 1.82946014 | +3 | 1.39351802 | 2.67002153 |
| | ISSGAFTPLVK | +2 | 1.06943501 | 1.92464415 | +2 | 0.54719266 | 1.40408516 |
| | SSGIESGAFQGMK | +2 | 0.11169496 | 0.54547246 | +2 | 0.49434041 | 1.94164133 |
| **CD44** | YAGVFHVEK | +2 | 1.056109 | 2.76455148 | +2 | 0.27005457 | -0.9179918 |
| | YGFIEGQVVIPR | +2 | 0.72334206 | 1.57499822 | +2 | 0.08397354 | -0.3173734 |

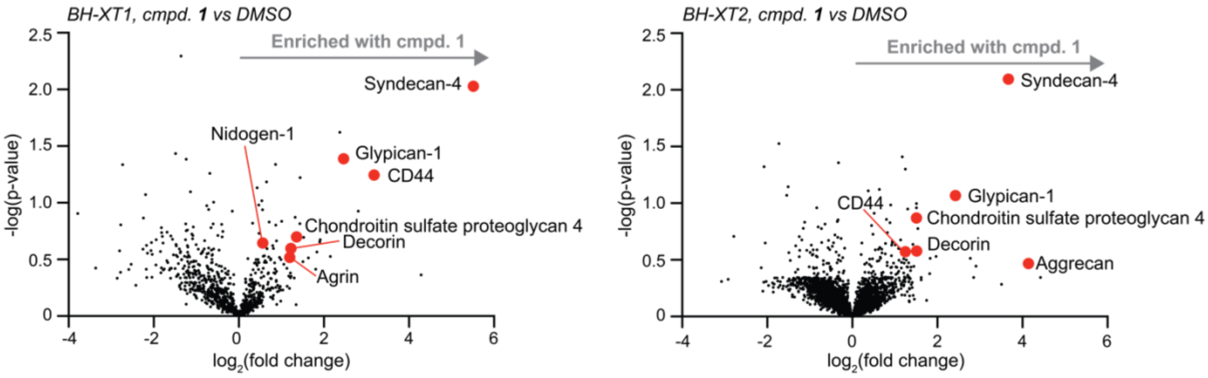


**Supporting Fig. 8:** Volcano plots of proteoglycans enriched by in differential MS experiments of secretome from BH-XT1/2-expressing pgsA-745 cells fed with compound **1** or DMSO from three independent replicates.

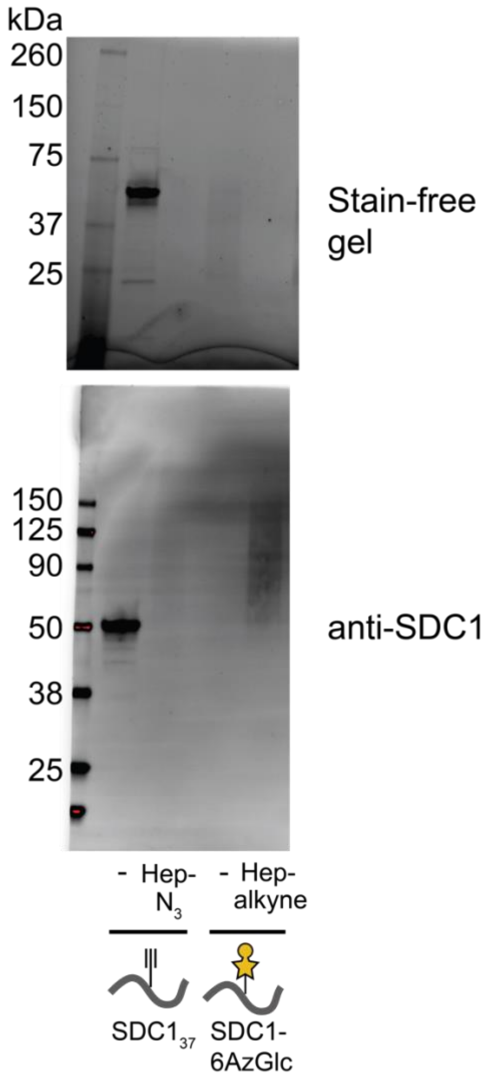


**Supporting Fig. 9:** Generation of SDC1-heparin conjugates. Recombinant $SDC1_{37}$ and SDC1 were expressed and purified as described,[1] and incubated with clickable heparin under CuAAC conditions. Conjugates were analysed by stain-free gel and Western Blot with anti-SDC1 detection. Data are from one experiment.

### Experimentals

#### Cloning of XT1 mutants for *in vitro* assays

**Supporting Table 2.** Primers used for generation of XT1 mutants

| Primer names | Sequence |
|---|---|
| XT1 W392N rev | 5'-ggaggctggctcctccgttgatggtggccattctc-3' |
| XT1 W392N for | 5'-gagaatggccaccatcaacggaggagccagcctcc-3' |
| XT1 W392V rev | 5'-aggctggctcctcccacgatggtggccattc-3' |
| XT1 W392V for | 5'-gaatggccaccatcgtgggaggagccagcct-3' |
| XT1 W392H rev | 5'-ggaggctggctcctccgtggatggtggccattctc-3' |
| XT1 W392H for | 5'-gagaatggccaccatccacggaggagccagcctcc-3' |
| XT1 W392G rev | 5'-ggctggctcctccgccgatggtggccatt-3' |
| XT1 W392G for | 5'-aatggccaccatcggcggaggagccagcc-3' |
| XT1 L525A rev | 5'-gaaggactcagcaggaagggcggtgtaggagtagaactg-3' |
| XT1 L525A for | 5'-cagttctactcctacaccgcccttcctgctgagtccttc-3' |
| XT1 L525G rev | 5'-gaaggactcagcaggaaggccggtgtaggagtagaactg-3' |
| XT1 L525G for | 5'-cagttctactcctacaccggccttcctgctgagtccttc-3' |
| XT1 L526A rev | 5'-ggaagaaggactcagcaggggccagggtgtaggagtagaac-3' |
| XT1 L526A for | 5'-gttctactcctacaccctggcccctgctgagtccttcttcc-3' |
| XT1 L526V rev | 5'-agaaggactcagcaggcaccagggtgtaggagtag-3' |
| XT1 L526V for | 5'-ctactcctacaccctggtgcctgctgagtccttct-3' |
| XT1 L526G rev | 5'-ggaagaaggactcagcagggcccagggtgtaggagtagaac-3' |
| XT1 L526G for | 5'-gttctactcctacaccctgggccctgctgagtccttcttcc-3' |
| XT1 L525A L526A rev | 5'-gtatggaagaaggactcagcaggagcggcggtgtaggagtagaactgtttca-3' |
| XT1 L525A L526A for | 5'-tgaaacagttctactcctacaccgccgctcctgctgagtccttcttccatac-3' |
| XT1 L525A L526V rev | 5'-gtatggaagaaggactcagcaggcacggcggtgtaggagtagaactgtttca-3' |
| XT1 L525A L526V for | 5'-tgaaacagttctactcctacaccgccgtgcctgctgagtccttcttccatac-3' |
| XT1 L525A L526H rev | 5'-gtatggaagaaggactcagcagggtgggcggtgtaggagtagaactgtttca-3' |
| XT1 L525A L526H for | 5'-tgaaacagttctactcctacaccgcccaccctgctgagtccttcttccatac-3' |
| XT1 L525A L526G rev | 5'-ccgtatggaagaaggactcagcagggccggcggtgtaggagtagaactgtttcatc-3' |
| XT1 L525A L526G for | 5'-gatgaaacagttctactcctacaccgccggccctgctgagtccttcttccatacgg-3' |
| XT1 W392A L526A rev | Use XT1 L526A rev on XT1 W392A pCEP plasmid |
| XT1 W392A L526A for | Use XT1 L526A for on XT1 W392A pCEP plasmid |
| XT1 pOPING Infusion rev | 5'-gtgatggtgatgtttaaactcacctgagccggccat-3' |
| XT1 pOPING Infusion for | 5'-gcgtagctgaaaccggtaccatgagggcctggatctt-3' |

A pCEP-Pu vector containing DNA coding residues of 232-959 of WT and W392A human xylosyltransferase 1 (original cDNA from Dharmacon, identifier BC045778, clone ID 4791553) with a TEV protease-cleavable His-tag at the N-terminus of the secreted protein was used previously.[2] The plasmid was used as a template to generate genes of single and double mutants of XT1 by overlap extension PCR using the Q5 HiFi polymerase (New England Biolabs) and bespoke primers (Table S1) designed with the Agilent QuikChange tool (https://www.agilent.com/store/primerDesignProgram.jsp, accessed on 4th Dec 2023), according to manufacturer's instructions. In PCR step 1, two 25 µL reactions were set up individually with 10 ng XT1 template DNA, 12.5 µL Q5 HiFi mastermix and either 0.5 µM primer mix of "XT1 pOPING for" and the "rev" primer for the targeted XT1 mutation, or "XT1 pOPING infusion rev" and the "for" primer for the targeted XT1 mutation. Conditions featured 1 min 98 °C denaturation, 25 cycles of 10 s at 98 °C, 30 s at 68 °C and 1 min at 72 °C, and a final extension of 15 min at 72 °C. PCR products from the two reactions were purified by agarose gel using a Macherey-Nagel ™ NucleoSpin Gel and PCR Clean-up kit (Thermo Fisher Scientific, Waltham, USA), concentration measured by nanodrop ranging from 10-50 ng/µL and then 2.5 µL of the purified PCR products were mixed with 0.5 µM primer mix of infusion for and infusion rev primers following the same PCR protocols as above. 100 ng of purified final PCR product was then mixed with 50 ng KpnI-HF and PmeI (New England Biolabs, Ipswich, USA). pOPING plasmid was a gift from Ray Owens (Addgene plasmid # 26046 ; http://n2t.net/addgene:26046 ; RRID:Addgene_26046)[3], digested with the same restriction enzymes, and inserts were introduced using an infusion HD cloning kit (Takara Bio, Kusatsu, Japan) according to the manufacturer's instructions. Stellar competent cells (Takara Bio) were transformed by heat-shock and used for plasmid amplification. All plasmids were confirmed by sanger sequencing (Genewiz, Leipzig, Germany) and nanopore sequencing (Plasmidsaurus, Eugene, USA) before use.

**Expression and enrichment of XT1 mutants**

Expi293F ™ cells (Thermo Fisher Scientific) were cultured in 10 mL Gibco® FreeStyle™ Medium (Thermo Fisher Scientific) in 50 mL cell culture flasks at 37 °C, 8% $CO_2$, and 125 rpm. The cells were sub-cultured to $0.8x10^6$ cells/mL two days before transfection at the exponential growth phase with a viability of at least 95%. Transfection of the Expi293F ™ cells was done at a density of $3x10^6$ cells/mL using the ExpiFectamine™ 293 transfection kit (Thermo Fisher Scientific). For each 10 mL cell culture, 10 µg DNA and 30 µL of ExpiFectamine™ 293 were diluted in 500 µL Opti-MEM® I (Thermo Fisher Scientific), respectively, with the ratio of DNA/Expifectamine™ 293 at 1:3 (w/v) and incubated at room temperature for five min. They were then mixed and incubated for 20 min at room temperature to allow the DNA-transfection reagent complexes to form before being added drop by drop to the cells. The cells were then incubated at the same condition for another 20-24 h before adding 50 µL ExpiFectamine™ 293 Transfection Enhancer 1 and 500 µL of Enhancer 2. The cell culture supernatants were harvested on the fifth day in 15 mL tubes by centrifuging at 500 g for 5 min and then 100x Halt™ protease inhibitor cocktail (Thermo Fisher Scientific) was added to a 1x final concentration.

HisPur™ Ni-NTA RA resin (Thermo Fisher Scientific) (100 µL beads slurry = 0.01 cell culture medium volume slurry) were washed with 10 column volumes (CV) of water and Equilibration Buffer (25 mM Tris-HCl pH 7.5, 150 mM NaCl, 20 mM imidazole) twice each before being added to the cell culture supernatant and incubated at 4 °C on a roller overnight. The cell culture supernatants were then centrifuged at 500 g for 10 min at 4 °C. The resulting resin was re-suspended and eluted sequentially with 10 CV Equilibration Buffer, 10 CV 25 mM Tris-HCl pH 7.5 with 150 mM NaCl and 50 mM imidazole, and 10 CV 25 mM Tris-HCl pH 7.5 with 150 mM NaCl and 200 mM imidazole twice sequentially and centrifugation at 380 g, 4 °C for 5 min after each wash to collect the supernatant. All fractions were checked by SDS-PAGE and those containing XT1 were pooled, concentrated to 600 µL by 30 KDa MWCO viva spin column (Cytiva, Marlborough, USA) and buffer exchanged with 50 mM Tris-HCl pH 7.5 with 150 mM NaCl and 20% (v/v) glycerol. Protein concentration was measured by NanoDrop One (Thermo

Fisher Scientific) as 0.05 to 0.12 mg/mL (yield: 30 ug- 72 ug). Protein was aliquoted into 50 µL per aliquot, flash-frozen in liquid nitrogen and stored at -80 °C.

**Analyses of enzyme specificities of XT1 mutants to UDP-Xylose, UDP-Glucose and UDP-6-Azido-Glucose**

A bikunin derived peptide described before[2] was conjugated to 5-carboxyfluorescein (FAM) through an ε-aminohexoic acid (ε-AHx) linker resulting in 5FAM-ε-Ahx-GQEEEGSGGGQGG-$CONH_2$ (**5FAM-bik**) as described.[4] Glycosylation assays with a total volume of 25 µL were carried out using 100 µM 5FAM-bik peptide, 0.2 µM purified XT1 (WT or mutants), and 200 µM UDP-xylose, UDP-glucose or UDP-6-azido-6-deoxyglucose (Biosynth, Staad, Switzerland) and incubated in Reaction Buffer (50 mM Tris-HCl pH 7.0 with 50 mM NaCl) at 37 °C overnight. The reaction mixtures were then boiled at 95 °C for 2 min to stop the glycosylation reaction and were briefly centrifuged. Of each reaction mixture, 5 µL were then run on a Acquity H-Class PLUS qDA UPLC-MS (Waters, Milford, USA) equipped with a ACQUITY UPLC® Glycan BEH Amide column (130 Å, 1.7 µm, 2.1 x 100 mm). Samples were run at flow rate of 0.35 mL/min using Buffer A: 10 mM Ammonium Formate at pH4.5; Buffer B: 10 mM Ammonium Formate in ACN: water 90:10. The percentage of turnover of **5FAM-bik** peptide into the corresponding glycopeptides was calculated by integration of the HPLC UV trace (260 nm absorption) of the peptide and the glycopeptide and calculated as Turnover % = Peak Area of Glycopeptide / Peak Area of (**5FAM-bik** + glycopeptide)%.

**Enzyme kinetics of WT- and BH-XT1**

Enzyme concentration titration was carried out to identify the concentration of WT- and BH-XT1 required for a turnover of maximum 10-20%. For purified XT1 preparations, a serial dilution from 500 nM to 3.9 nM was used in Reaction Buffer. Following this, 200 µM UDP-sugar and 100 µM synthetic fluorescent peptide **5FAM-bik** were added with a total volume of 25 µL. The reaction mixtures were incubated at 37 °C for 1.5 h, stopped by incubation at 95 °C for 2 min and then briefly centrifuged. Aliquots of 5 µL of the reaction mixtures were checked by HPLC and the turnover rate was calculated to give 16 nM WT-XT1 for incubation with UDP-Xyl, 30 nM $XT1^{L526G}$ (BH-XT1) for incubation with UDP-6AzGlc and 100 nM BH-XT1 for incubation with UDP-Xyl as suitable concentrations for Michaelis-Menten kinetics. A serial dilution of UDP-sugar from 400 to 3.125 µM was run in glycosylation reactions in 20 µL Reaction Buffer with 100 µM synthetic fluorescent peptide **5FAM-bik** and the optimal concentration of WT- or BH-XT1. The reaction mixtures were incubated at 37 °C for 1.5 hours, reactions stopped by incubation at 95 °C for 2 min and briefly centrifuged. Aliquots of 5 µL of the reaction mixtures were checked by HPLC, the turnover calculated and transformed into rate with the formula 1000x{[Peak Area of Glycopeptide/Peak Area of (**5FAM-bik** + glycopeptide)]x100 µM}/5400 s to give the rate v in nM/s. The kinetics curve was then plotted with Prism 10 (GraphPad, San Diego, USA) and fitted with a Michaelis Menten function to calculate $k_{cat}$, $K_M$ and $v_{max}$.

**Peptide specificity analyses**

BH-XT1 activity toward a collection of substrate peptides was assessed by using a UDP-Glo™ Glycosyltransferase assay kit (Promega, Madison, USA) following the manufacturer's instructions. The collection of 240 peptides was described before.[2] In brief, 25 µL reactions contained 25 nM BH-XT1, 100 µM UDP-6AzGlc and 25 µM peptide in 50 mM Tris-HCl pH 7.0 with 50 mM NaCl in 96 well plates. Reactions were shaken at 350 rpm for 30 s on a thermomixer (Thermo Fisher Scientific) at room temperature and then incubated at room temperature for 1 h followed by adding 25 µL freshly prepared UDP-Glo reagent for detection. A series of diluted UDP standards from 0-25 µM were included in each set of experiments to plot the standard curve, and blank UDP-6AzGlc was run as background subtraction. Luminescence was read using a plate-reader (Tecan, Männedorf, Switzerland) with luminescence read-out at 1000 ns integration. All luminescence data across different 96-well plates were normalized and presented as heat map in three independent experiments.

***In vitro* glycosylation of membrane fractions**

CHO cells with a CRISPR-KO for either *Xylt1* or *Xylt2* or parental CHOZN GS-/- (WT CHO cell with glutamine synthetase KO) were cultured in EX-CELL® CD CHO fusion medium (Sigma-Aldrich, St Louis, USA) supplemented with 2% Glutamax (Thermo Fisher Scientific) as described.[5] Supernatants (15 mL) were collected when cells were at a density of $1x10^6$ cells/mL, concentrated with 30 kDa MWCO Amicon ultra centrifugal filters (Merck, Darmstadt, Germany), aliquoted and stored at -80 °C. Cell pellets were treated with the subcellular fractionation kit for cultured cells (Thermo Fisher Scientific) following the manufacturer's instructions. Membrane fraction protein concentrations were measured by Pierce BCA protein assay (Thermo Fisher Scientific). Glycosylation reactions were run in 25 µL 50 mM Tris-HCl pH 7.0 with 50 mM NaCl with 16 µg membrane protein, 200 nM WT- or BH-XT1, 250 µM UDP-6AzGlc and 100/200/300 µM UDP-Xyl. The reaction mixtures were incubated at 37 °C overnight, treated with 7.5 µL of a click reaction mastermix (final concentration 1200 µM BTTAA, 600 µM CuSO4, 100 µM biotin-alkyne, 5 mM sodium ascorbate and 5 mM aminoguanidinium chloride), incubated at room temperature overnight and quenched by the addition of 3 µL 50 mM EDTA. Reaction mixtures were then subjected to SDS-PAGE and Western Blot. Total protein and streptavidin signal was recorded as described above.
Competition glycosylation reactions were run as described above, except that membrane fractions were pre-incubated with WT-XT1 and UDP-Xyl before addition of BH-XT1/UDP-6AZGlc in the indicated concentrations and incubating for another 16 h. Reactions were treated as described above.

***In vitro* glycosylation of recombinant, GAG-free Glypican-1**

GAG-free glypican-1 was prepared as reported.[4] For each sample, a 25 µL reaction mixture contained 200 nM WT-/BH-XT1, 250 µM UDP-Xyl and/or UDP-6AzGlc and 16 µM glypican-1 in 50 mM Tris-HCl pH 7.0 and 50 mM NaCl. Reactions contained WT-XT1 and UDP-Xyl (A and D), WT-XT1, UDP-Xyl and UDP-6Az-Glc (B), BH-XT1, UDP-Xyl and UDP-6AzGlc (C). Reactions A-D were incubated at 37 °C overnight, when 200 nM BH-XT1 and 250 µM UDP-6AzGlc were added to reaction D, which was then incubated for an additional 16 h. Following this, 7.5 µL of a click reaction mastermix was added to give a final concentration of 1200 µM BTTAA (Jena Bioscience, Jena, Germany), 600 µM $CuSO_4$ (Sigma-Aldrich), 100 µM biotin-alkyne (Biotium, Fremont, USA); 5 mM sodium ascorbate (Thermo Fisher Scientific) and 5 mM aminoguanidinium chloride (Cayman Chemical, Ann Arbor, USA). Click reactions were carried out at room temperature overnight at 350 rpm and later quenched by the addition of 3 µL 50 mM EDTA. Reaction mixtures were subjected to SDS-PAGE and Western Blot. Total protein content was measured on and Odyssey CLx (LI-COR Biosciences, Lincoln, USA) using the Revert Total Protein kit (LI-COR Biosciences) and biotinylation was assessed with IRDye 800CW Streptavidin (LI-COR Biosciences). FLAG-tagged decorin was visualised with a rabbit polyclonal antibody (invitrogen, PA1-984b) and a 700CW anti-rabbit antibody (LI-COR Biosciences).

***In vitro* glycosylation of recombinant, GAG-free decorin**

GAG-free decorin in pgsA-745 CHO cells was prepared before.[4] For each sample, a 25 µL reaction mixture contained 200 nM WT-/BH-XT1, 250 µM UDP-Xyl and/or UDP-6AzGlc and 0.64 µM decorin in 50 mM Tris-HCl pH 7.0 and 50 mM NaCl. Reactions contained WT-XT1 and UDP-Xyl (A and D), WT-XT1, UDP-Xyl and UDP-6Az-Glc (B), BH-XT1, UDP-Xyl and UDP-6AzGlc (C). Reactions A-D were incubated at 37 °C overnight, when 200 nM BH-XT1 and 250 µM UDP-6AzGlc were added to reaction D, which was then incubated for an additional 16 hours. Following this, 7.5 µL of a click reaction mastermix was added to give a final concentration of 1200 µM BTTAA, 600 µM $CuSO_4$, 100 µM biotin-alkyne; 5 mM sodium ascorbate and 5 mM aminoguanidinium chloride. Click reactions were carried out at room temperature overnight at 350 rpm and later quenched by the addition of 3 µL 50 mM EDTA. Reaction mixtures were subjected to SDS-PAGE and Western Blot. Total protein content was measured on and Odyssey CLx (LI-COR Biosciences) using the Revert Total Protein kit (LI-

COR Biosciences) and biotinylation was assessed with IRDye 800CW Streptavidin (LI-COR Biosciences). FLAG-tagged decorin was visualised with rabbit polyclonal anti-FLAG and a 700CW anti-rabbit antibody (LI-COR Biosciences).

**Sample prep for mass spectrometry analyses of glycosylated decorin**
GAG-free decorin protein (10 µg) was incubated with 250 µM UDP-6-AzGlc and 200 nM BH-XT1 in 50 mM Tris-HCl pH 7.0 with 50 mM NaCl in a 30 µL total reaction volume at 37 °C overnight. The glycosylation reaction was treated with 7.5 µL of a click reagent mastermix containing 1-(3-Butyn-1-yl)-3-methylimidazolium tetrafluoroborate described before[6] and termed ITag-alkyne (final concentration 1200 µM BTTAA, 600 µM $CuSO_4$, 100 µM Itag-alkyne, 5 mM sodium ascorbate and 5 mM aminoguanidinium chloride). The reaction was incubated by shaking at 350 rpm on a thermomixer at room temperature overnight. The clicked decorin sample was digested by trypsin in-solution with S-trap (Protifi, Fairport, USA) following the manufacturers' instructions.

**Mass spectrometry data acquisition for glycopeptide analysis**
Samples were analysed by online nanoflow liquid chromatography-tandem mass spectrometry using an Orbitrap Eclipse Tribrid mass spectrometer (Thermo Fisher Scientific) coupled to a Dionex UltiMate 3000 HPLC (Thermo Fisher Scientific). For each analysis, 14 µL were injected onto a trap column (Acclaim PepMap 100, 75 µm x 2 cm NanoViper) with loading buffer (2% Acetonitrile, 0.05% TFA) at 7 µl min−1 for 6 min (40 °C). Glycopeptides were then separated on an analytical column (PepMap RSLC C18, 75 µm x 50 cm, 2 µm particle size, 100 Å pore size, reversed-phase EASY) using a gradient 2-40% solvent B (5% DMSO, 0.1% formic acid, 75% acetonitrile, 20% water) over 140 min at 275 nL/min.
Full scan MS1 spectra acquired in the Orbitrap were collected at a resolution of 120,000 at FWHM (full width at half-maximum peak) and a mass range from 300 to 1500 m/z.
Dynamic exclusion was enabled with a repeat count of 3, repeat duration of 10 s, and exclusion duration of 10 s. Only charge states 2 to 6 were selected for fragmentation. MS2 scans were generated at top speed for 3 s. Higher-energy collisional dissociation (HCD) was performed on all selected precursor masses with the following parameters: isolation window of 2 m/z, 28% normalized collision energy, orbitrap detection (resolution of 30,000), maximum inject time of 54 ms, and a standard AGC target. An additional electron transfer dissociation (ETD) fragmentation of the same precursor was triggered if 1) the precursor mass was between 300 and 1,500 m/z and 2) the fingerprint ion generated by the specific tag (322.1507 for ITag-Alkyne) was present at ± 0.1 m/z and greater than 10% relative intensity.

**Mass spectrometry data analysis of glycopeptides**
Raw files were searched using Byonic™ (Protein Metrics, Cupertino, USA, version 4.6.1).
For glycopeptide analysis, search parameters included semi-specific cleavage specificity at the C-terminal site of R and K, with two missed cleavages allowed. Mass tolerance was set at 10 ppm for MS1s, 20 ppm for HCD MS2s, and 0.2 Da for ETD MS2s. Carbamidomethyl cysteine was set as a fixed modification. Variable modifications included methionine oxidation (common 1), asparagine deamidation (common 1). O-glycans modification was set to HexNAc with an additional 118.0643 m/z to account for the chemical modification. A maximum of two variable modifications were allowed per peptide. For each sample, variable modifications were searched against a focused FASTA file that exclusively contains protein sequences find in that sample. Identifications that contained chemically modified glycans on the peptide of interest were manually validated and localised using a combination of HCD and ETD information.

**Analysis of UDP-sugar biosynthesis**
This protocol was adapted from previously published procedures.[7–9] PgsA-745 CHO cells were cultured with F12K medium (Thermo Fisher Scientific) supplemented with 10% (v/v) FBS, penicillin (100 U/mL) and streptomycin (100 µg/mL). K-562 cells (ATCC CCL-243) were cultured in RPMI (Thermo Fisher Scientific) with 10% (v/v) FBS, penicillin (100 U/mL) and streptomycin (100 µg/mL). Approx. 5-10 million cells (comparable numbers between treatment

conditions) were fed with DMSO or 250 µM caged sugar-1-phosphates **1** or **2**. After 16 hours incubation, K-562 cells were harvested by centrifugation (500g, 5 min, 4 °C) and CHO psgA-745 cells were harvested using cell scrapers. Cells were washed with ice cold PBS twice and Zirconia/silica beads (0.1 mm, Biospec, Bertilesville, USA) were added at a volume to packed cell pellets with a 1/1 ratio. 1 mL of 1/1 acetonitrile/water was added to lyse the cells with a bead beater at 6m/s for 30 s, and the cell lysates was cooled at 4 °C for 10 min. They were then centrifuged at 14000 g, for 10 min at 4 °C and the supernatant was transferred to a protein low bind Eppendorf. Supernatants were dried down with a SpeedVac and the residue was dissolved in 0.3 mL Milli-Q water. The supernatant was passed through a centrifuge filter (30 min, 14000 g) using a 3 kDa MWCO Amicon ultra centrifugal filter. The flow-through was dried by SpeedVac and the residue resuspended in 50 µL of MQ water. High Performance Ion Exchange Chromatography was used to analyse lysates, employing a Waters Arc Premier HPLC with Photodiode Array Detector equipped with a 2mm x 150mm Dionex CarboPac PA1 column and matching PA1 guard column at a flow rate of 0.25 mL/min. The gradient was Buffer A (1 M NaOAc/1 mM NaOH), B (1 mM NaOH), C (1 M NaOH); 0 min, 5% A, 95% B; 20 min 40% A, 60% B; 60 min, 40% A, 60% B; 63 min, 50% A, 50% B; 87 min, 80% A, 20% B; 95 min, 80% A, 20% B; 96 min, 5% A, 95% B; 101 min, 5% A, 95% B.

**Site-directed mutagenesis and cloning of full-length XT1**
A full length XT1 plasmid in pDonor221 vector was purchased from DNASU (HsCD00744626). Site directed mutagenesis was performed using Q5 HIFI mutagenesis kit following manufacturer's instructions using the two mutagenesis primers CTACACCCTCGGCCCCGCTGAGTC and GAGTAGAACTGTTTCATCTTG. Full-length mutant DNA was then prepared using the two primers ATGGTGGCGGCGCCAT and CCGTAGCCGGCCATCAG. To add a VSV-G tag, a third PCR was done using the primers ACCCCAAGCTGGCCTCTGAGGCCATGGTGGCGGCGCCATGCGCCCG and CCCCAAGCTTGGCCTGACAGGCCCTACTTACCCAGGCGGTTCATTTCGATATCAGTGTA CCGTAGCCGGCCATCAG. pSBbi-GH was a gift from Eric Kowarz (Addgene plasmid # 60514 ; http://n2t.net/addgene:60514 ; RRID:Addgene_60514).[10] The plasmid was linearized using restriction enzyme SfiI (NEB) following manufacturer's instructions and purified from agarose gel. Full length WT- and BH-XT1 were then inserted into pSBbi-GH using infusion cloning kit following the manufacturer's instructions. Genes of interest were sequenced by sanger sequencing and the full plasmids were sequenced by nanopore sequencing before use.

**Stable transfection of pgs745 cells with full-length WT- and BH-XT1**
The plasmid pCMV(CAT)T7-SB100 was a gift from Zsuzsanna Izsvak (Addgene plasmid # 34879 ; http://n2t.net/addgene:34879 ; RRID:Addgene_34879).[11] PgsA-745 CHO cells were cultured in growth medium to $0.5x10^6$ cells/mL 24-48 hours before transfection. 2.5 µg pSBbi plasmid containing full length WT- or BH-XT1 were transfected together with 125 ng pCMV(CAT)T7-SB100 plasmid per well of a 6-well plate using Lipofectamine LTX (Thermo Fisher Scientific) according to manufacturer's instructions. After 24 hours, cell culture medium was aspirated, and cells were treated with fresh growth medium containing 200 µg/mL hygromycin B. Cells were cultured under these conditions for two weeks to obtain stable cells. Following selection, cells were propagated with 150 µg /mL hygromycin B in growth medium.

**Metabolic cell surface labelling and in-gel fluorescence**
Stably transfected pgsA-745 CHO cells with full length WT- or BH-XT1 were plated in 6-well plates at a density of $0.4x10^6$ cells/mL in growth medium without hygromycin B and then treated with the indicated concentration of caged sugar-1-phosphates **1** or **2**, $Ac_4ManNAz$ (Jena Bioscience) or DMSO. Cells were grown for 16 h. The cell culture medium was aspirated and cells washed with cold PBS without $Ca^{2+}/Mg^{2+}$. The cells were then detached with 1 mL ice-cold 8 mM EDTA for 20 min at 4 °C. Cells were transferred to a 1.5 mL cup and harvested by centrifugation at 500 g for 5 min at 4 °C.
Cells were then resuspended in 200 µL Cell buffer (2% FBS in PBS), transferred to a V-shaped 96 well plate (Thermo Fisher) and harvested by centrifugation at 500 g for 5 min at 4

°C. The cells were re-suspended in 35 µL Cell buffer and treated with 35 µL click solution mastermix (200 µM CuSO4, 1200 µM BTTAA, 10 mM sodium ascorbate, 10 mM aminoguanidinium chloride and 50 mM CF680 alkyne in Cell Buffer). Cells were briefly mixed and incubated for 7 min at room temperature on an orbital shaker. The reaction was quenched by addition of 35 µL 3 mM bathocuproinedisulfonic acid in PBS. Cells were then harvested, washed twice with 200 µL cell buffer and once with PBS and then resuspended in 100 µL ice-cold Lysis Buffer [50 mM Tris-HCl pH 8 with150 mM NaCl, 1 mM $MgCl_2$, 0.5% (w/v) sodium deoxycholate, 0.1% (w/v) SDS, 1% (w/v) Triton X-100, 1x Halt™ protease inhibitor, and 100 mU/µL benzonase (Merck)]. Cells were lysed for 20 min at 4 °C on an orbital shaker and centrifuged at 1500 g, 20 min at 4 °C. The supernatants were then transferred to 1.5 mL protein LoBind® Tubes (Eppendorf). The protein concentration was measured by Pierce BCA Protein Assay kits before in-gel fluorescence. Samples were then analysed after SDS-PAGE by in-gel fluorescence on an Odyssey CLx (LI-COR Bosciences). Total protein content was then assessed by Coomassie staining on the same gel using SafeBLUE Protein Stain (NBS Biologicals, Huntingdon, UK). Another gel was prepared and transferred to a nitrocellulose membrane for Western blot, using antibodies against Rabbit VSV-G tag (Abcam, ab50549), GAPDH (Abcam, ab181602) and secondary antibodies IRDye 800CW donkey anti-mouse (LI-COR) and IRDye 680RD Donkey anti-rabbit IgG (LI-COR). Image background was adjusted by LI-COR software.

**Chemoenzymatic GAG linker synthesis on 5FAM-bik peptide**

Xyl- or 6-AzGlc-containing glycopeptides were generated enzymatically in a 100 µL reaction with 100 µM **5FAM-bik** peptide, 16 nM WT-XT1 and 200 µM UDP-Xyl or 30 nM BH-XT1 and 200 µM BH-XT1, respectively. When full conversion was reached, glycopeptides were desalted with A Strata-X (60 mg /mL) Phenomenex solid phase extraction column following manufacturer's instructions and dried with Genevac™ miVac centrifugal concentrator (Fisher Scientific) before downstream one-pot enzyme reactions. These were carried out in 50 mM Na-HEPES pH 7.5, 25 mM MnCl2 and 50 mM NaCl in a total reaction volume of 20-30 µL as reported.[4] Soluble enzymes for extension (B4GALT7, MBP-B3GALT6, B3GAT3) were prepared before.[4] These enzymes were added at 0.025 µg/mL in different combinations to glycopeptides at 500 µM, and the UDP-sugars UDP-Gal and UDP-GlcA were added as needed at a two-fold molar excess to the acceptor. The reaction mixture was left at 30 °C overnight and the reaction progress was monitored by HPLC as reported before.[4]

**Expression and cellular glycosylation by BH-XT1 of decorin in pgsA-745 cells**

An expression construct of human decorin containing a C-terminal FLAG tag was prepared previously.[4] Cultured pgsA-745 cells stably transfected with full length BH-XT1 were plated at $1x10^6$ cells/mL in 5 mL growth medium in a T25 flask and grown to 100% confluency. The cells were then detached using 0.05% (v/v) Trypsin-EDTA (Thermo Fisher Scientific) and re-suspended in 5 mL fresh medium. While the cells were still in suspension, 5 µg plasmid DNA and 15 µg polyethylenimine (PEI) MAX 40 kDa (Polysciences, Warrington, USA) were diluted separately in 250 µL OptiMEM (Thermo Fisher Scientific) and incubated at room temperature for 5 min. Both solutions were mixed and incubated at room temperature for another 20 minutes before added to the cell culture dropwise. The cells were incubated overnight to attach, and 50 µM caged sugar-1-phosphate **1** was added. On the fourth day after transfection, the culture supernatant was collected. A 100 µL slurry of Pierce™ anti-DYKDDDDK affinity resin (Thermo Fisher Scientific) was washed with 30 CV FLAG buffer (25 mM HEPES with 150 mM NaCl) and added to the cell culture medium. The suspension was incubated at 4 °C overnight, then centrifuged at 4 °C at 1000 g for 5 minutes. The supernatant was removed and the beads were incubated with FLAG buffer containing 100 µg/mL FLAG® peptide (Sigma-Aldrich) for 1 h at room temperature. A 3 kDa MWCO Amicon ultra centrifugal filter was used to concentrate the protein to 300 µL and remove the FLAG peptide. The protein concentration was measured by BCA and the final yield of the protein estimated as 120 µg.

An aliquot of 20 µg purified decorin in 60 µL FLAG buffer was treated with 30 µL CuAAC mastermix (final concentration 1200 µM BTTAA, 600 µM $CuSO_4$, 100 µM Itag-alkyne, 5 mM sodium ascorbate and 5 mM aminoguanidinium chloride) overnight at room temperature.
The clicked decorin sample was digested by trypsin in-solution with S-trap (Protifi) following manufacturer's instructions. Mass spectrometry data acquisition and analysis was performed as described above.

**Site-directed mutagenesis of full length XT2 and cloning into pSBbi plasmids**
A full length XT2-pJF7_nHalo vector was purchased from DNASU (accession number sCD00866744). Site-directed mutagenesis was performed by overlap extension (see above) using the mutagenesis primers AGAAGGACTCGGCTGGGCCCAGTGTGTATGTGTAG and CTACACATACACACTGGGCCCAGCCGAGTCCTTCT as well as the two ATGGTGGCGAGCGCGCGAG and CAACCTGAGTCGCCCGTCTG. PCR reaction conditions are as stated above. Following assembly of the full-length XT2 gene, another PCR was performed using the primers AACTACCCCAAGCTGGCCTCTGAGGCCATGGTGGCGAGCGCGCGAG and CCCCAAGCTTGGCCTGACAGGCCCTACTTACCCAGGCGGTTCATTTCGATATCAGTGTA CAACCTGAGTCGCCCGTC. The PCR product was inserted into pSBi-GH by infusion cloning. Genes of interest were sequenced by sanger sequencing and the full plasmids were sequenced by nanopore sequencing before use.
Cell-surface labelling and in-gel fluorescence were performed as described above.

**Sample prep for proteomics analysis**
Stably transfected pgsA-745 CHO cells with full length WT- or BH-XT1 were seeded in T-75 flasks at a density of $1x10^6$ cells/mL in growth medium. After 6 h of incubation, cells were fed with either 50 µM caged sugar-1-phosphate **1** or DMSO. The cell culture medium used for feeding was BalanCD CHO growth A medium (Fujifilm Irvine Scientific). Cells were grown for 16 h. The cell culture medium (secretome) was harvested and centrifuged (350 g, 5 min) to pellet debris. The secretome samples were concentrated using 3 kDa MWCO Amicon ultra centrifugal filters. The buffer was exchanged with PBS twice. To prepare lysates, cells were washed with cold PBS without Ca2+/Mg2+ and were then detached with 5 mL ice-cold 8 mM EDTA for 20 min at 4 °C. Cells were transferred to a 15 mL tube and harvested by centrifugation at 500 g for 5 min at 4 °C. The cells were lysed with 200 µL of ice-cold Lysis Buffer. Cells were lysed for 20 min at 4 °C on an orbital shaker and centrifuged (1500 g, 20 min, 4 °C). Supernatant was transferred to a new plate and Pierce™ BCA Protein Assay kit was used to measure protein concentration of both lysates and secretome.
Both lysates (0.20 mg each) and secretome (0.50 mg each) samples were normalised up to 250 µL with PBS and incubated for 1 h at RT with 300 µL of Neutravidin beads slurry (Sera-Mag SpeedBeads Neutravidin-Coated Magnetic Beads, cytiva, Marlborough, USA), previously washed with PBS (2x200 µL), to remove endogenous biotinylated proteins. The supernatant was collected and then incubated with PNGase F overnight at 37 °C to remove N-glycans. The reaction was then quenched by heating to 95 °C for 10 s with subsequent cooling at 4 °C. The samples were then treated with a 10x click solution mastermix (6 mM CuSO4, 12 mM BTTAA, 1 mM biotin-DADPS-alkyne (Vector Laboratories, Newark, USA), 50 mM sodium ascorbate and 50 mM aminoguanidinium chloride) to a 1x final concentration. The click reaction was incubated overnight at rt under shaking (400 rpm). The reaction was passed through 3 kDa MWCO Amicon ultra centrifugal filters Amicon to exchange the buffer with PBS. The samples were then incubated with 350 µL of dimethylated Neutravidin Beads slurry (previously washed twice with 200 µL of PBS) for 1 h at rt.[12] Supernatant was discarded and beads were washed with 1% (w/v) SDS (3x350 µL), 6 M urea in PBS (3x350 µL), AmBic (50 mM ammonium bicarbonate, 3x350 µL) and 40% (v/v) LCMS-grade acetonitrile (4x100 µL). Beads were resuspended in 100 µL of AmBic containing 10 mM DTT and then incubated at 50 °C for 15 min. Beads were washed with AmBic (2x350 µL) and 100 µL of 20 mM iodoacetamide in AmBic was then added. Samples were kept for 30 min in the dark.

Iodoacetamide was then quenched by adding DTT 10 mM (final concentration). The beads were washed with AmBic (3x350 µL), then resuspended in 100 µL of AmBic and 300 ng of Lys-C Mass Spec Grade (Promega) were added to beads followed by overnight incubation at 37 °C. The supernatant was transferred to a new tube and 200 ng of Trypsin gold Mass Spec Grade (Promega) were added. The digestion was left for 8 h at 37 °C. Peptides were desalted by UltraMicroSpin™ (The Nest group Inc., Ipswich, USA) according to the manufacturer's protocol and vacuum-dried by SpeedVac.

Dried peptides were resuspended in 16 µL of 0.1% (v/v) formic acid in LCMS-grade water, sonicated for 15 min in a water bath, vortexed briefly and harvested for 5 min at 18000 g. The peptides were then loaded on Evotips (Evosep, Odense, Denmark) according to manufacturer's protocol. The data was acquired on TIMS TOF Pro2 (Bruker) coupled to Evosep One LC system. For the LC separation, a standard 60SPD 2.3 method was used, separation was performed using an EV-1109 column, and the column was heated to 40 °C during analyses. TIMS TOF Pro2 was operated in DIA PASEF mode, scan width was set to 100-1700m/z with the IM 1K/0 0.6 - 1.6, ramp and accumulation time was locked at 100 ms.

Raw MS files were loaded into DIANN 1.8.1 for quantification and identification by using the *Cricetulus griseus* FASTA protein sequences database from UniProt for database search. The DIANN output files, both protein groups and peptide groups were uploaded into Perseus (version 2.0.11)[13] to allow for data transformation and visualization, and into graph pad for statistical analysis. Briefly, to visualize the results on Perseus, the proteingrougs.txt file or peptidegroups.txt file were uploaded, followed by transformation of all the values to $\log_2(x)$. Then the data was imputed to replace missing values from normal distribution. Data from three independent replicate experiments of each sample in a row were categorically annotated with the same name. Once annotated, a two sample Welsh t-test was performed to statistically analyse the data. The Welch t-test was performed between samples from BH XT1 vs WT XT1-expressing cells and BH XT2 vs WT XT2-expressing cells, respectively. Multiple-testing corrections were performed using the Benjamini-Hochberg procedure to calculate p values (False Detection Rate). Protein hits were filtered using a p value of 0.5. The scatter plot function was used to visualize the volcano plots.

**Preparation of alkyne-heparin**

A previously published procedure was modified to produce alkyne-modified heparin.[1,14] 20 mg heparin (Iduron, HEP001) was dissolved in 94 µL 100 mM sodium acetate, 100 mM aniline buffer (pH 5.5) and prewarmed to 55°C. Warmed heparin was mixed with 6 µL of alkyne hydrazide (BroadPharm BP-28990, 1.68 mg in DMSO, 80 eq.), and DMSO was added to bring the aqueous buffer:DMSO ratio to 1:1. The mixture was protected from light and incubated at 55°C for 72 hr before dilution into 10 mL PBS, 0.45 µm filtered and dialyzed into MQ $H_2O$ (48 hr, buffer changed 3 times). The sample was then lyophilized. $^1$H NMR verified the conjugation of alkyne-hydrazide to heparin.

**Production of SDC1 and SDC1$_{37}$**

The human syndecan-1 ectodomains were cloned into pET28a expression vectors for production in BL21 (DE3) *E. coli*. SDC1$_{37}$ refers to a variant ectodomain wherein the canonical GAG attachment site S37 is replaced by the unnatural amino acid p-propargyltyrosine (pPY) with an alkyne handle for click chemistry. This variant was produced as previously described,[1] wherein the expression plasmid was co-transformed into BL21 with pULTRA-CNF which permits incorporation of unnatural amino acids. pPY (400 mg/L) was added to the bacterial culture during induction. Protein was purified using HisPur cobalt resin (ThermoFisher). SDC1$_{37}$ was treated with azide-containing heparin as previously described.[1]

**In vitro glycosylation of SDC1 with BH-XT1/UDP-6AzGlc**

Recombinant SDC1 and $SDC1_{37}$ were *in vitro* glycosylated with 6AzGlc using the procedure established for decorin (see above). In brief, two 78.5 µL reaction mixtures containing 15 µM of SDC1 or $SDC1_{37}$, 200 nM BH-XT1, 250 µM UDP-Xyl and UDP-6AzGlc in 50 mM Tris-HCl pH 7.0 with 50 mM NaCl were incubated at 37 °C overnight. The enzymes were then deactivated by incubation of the reaction mixtures at 95 degrees for 2 minutes. The glycosylation reaction mixtures were then buffer exchanged with the reaction buffer using 3 kDa MWCO Amicon ultra centrifugal filters to remove excess UDP sugars. The supernatants containing glycosylated SDC1 and SDC137 were then collected and dried under SpeedVac before proceeding to the next step to be clicked with the alkyne-heparin.

**SDC1 click reactions**

6AzGlc-modified hSDC1 (100 µM) and alkyne-heparin (20 mol eq.) were dissolved in PBS with aminoguanidine (5 mM) in Protein LoBind Eppendorf tubes. Click reagents (320 µM $CuSO_4$ 1600 µM tris(3-hydroxypropyltriazolylmethyl)amine (THPTA), 21 mM sodium ascorbate) were added and the reactions were incubated at 37°C. Reactions were monitored using an Ultimate 3000 UHPLC system and WAX-10 (4x250 mm) column at 1.0 mL/min in 20 mM Tris (pH 7.5) buffer. After reaction completion (~16h), 5 µL of HisPur cobalt resin beads (ThermoFisher, #89966) and an equal volume of wash buffer (10 mM imidazole/PBS, 40 µL) were added to each reaction. This mixture was incubated on a VortexGenie with the Multiple Sample attachment for 1h at RT. The supernatant was removed, and beads were washed twice with wash buffer (5 mins, orbital shaker), before incubation with elution buffer (150 mM imidazole/PBS, 5 mins, orbital shaker). Eluate was concentrated using a 3K MWCO filter and buffer exchanged into PBS. The concentration of glycoconjugate products was measured on a NanoDrop One using absorbance at 205 nm.

**Cell spreading assay**

MDA-MB-231 cells were treated with 200 nM pooled SDC1 TriFECTA Dicer substrate RNAs (hs.Ri.SDC1.13) using Lipofectamine RNAiMAX. 24 hours after transfection, 24-well plates were coated with 1X poly-D-lysine (15 mins, RT) before incubation with 10 µg/ml vitronectin (4°C, O/N, rocking). The following day, the plate was washed twice with PBS and blocked in 2% BSA/DMEM (1 hr, 37°C). Cells were harvested with non-enzymatic dissociation buffer and remodeled in 96-well round bottom plates with 10 µM cholPEGNTA (1h, 37°C) followed by SDC constructs (2 µM, 1h, 37°C). Cells were resuspended in DMEM + 10% FBS and allowed to adhere to VN coated plates (O/N, 37°C). Cells were then fixed with 4% PFA/PBS and stained with rhodamine-phalloidin (2 U/mL, 1h, RT, rocking), followed by Hoechst staining. Cells were imaged on an EVOS M500 fluorescence microscope. The extent of cell spreading and cell number were counted by ImageJ macros using previously published methods.[1]

**Supporting Table 3.** siRNA locations used for SDC1 knock-down.

| DsiRNA | Cross-reacting transcript | Location | Exon |
|---|---|---|---|
| 1 | NM_002997 | 3' UTR | 5 |
| | NM_001006946 | 3' UTR | 6 |
| 2 | NM_001006946 | CDS | 4 |
| | NM_002997 | CDS | 3 |
| 3 | NM_002997 | CDS | 3 |
| | NM_001006946 | CDS | 4 |

**Synthetic Chemistry**
All chemicals were from commercial vendors. (N-[(*S*)-(2,3,4,5,6-Pentafluorophenoxy)phenoxyphosphinyl]-L-alanine 1-methylethyl ester was purchased from Biosynth (Staad, Switzerland). Anhydrous THF was obtained by passing solvent through activated alumina columns and dispensed from a PureSolv MD ASNA solvent purification system and stored over 4 Å molecular sieves. Unless otherwise stated, all reactions were conducted using anhydrous solvents, under an atmosphere of $N_2$ which was passed through a Drierite® drying column. HRMS (ESI, NSI) were obtained on Agilent 6530 Q-TOF, LQT Orbitrap XL1 or Waters (Xevo, G2-XS TOF or G2-S ASAP) Micromass LCT spectrometers using a methanol mobile phase in positive/ negative ionisation modes as appropriate. Analytical thin layer chromatography (TLC) was carried out on pre-coated 0.25 mm Merck KgaA 60 F254 silica gel plates. HPLC was performed using an Agilent 1260 Infinity II preparative HPLC system equipped with a variable wavelength detector and a fraction collector, on a reverse phase column (Polaris 180 Å C18-A. 21.2 × 250 mm, 5 μm) to achieve a purity level >95% and to determine percent purity. HPLC method: Polaris 180 Å C18-A. 21.2 × 250 mm, 5 μm, Solvent A: acetonitrile, solvent B: water, a gradient elution from 5% to 95% B over 20 mins, held for 7 mins, elution reversed to 5%B over 2 mins then held for 7 mins. Flow rate 3mL/min and UV detection at 254nm. Visualisation was achieved using UV detection at 254 nm. Visualisation was by adsorption of UV light, thermal development or thermal development after dipping in a methanolic solution of sulfuric acid (5% v/v). Column chromatography was carried out on 40–63 μm silica gel (Sigma-Aldrich) under a positive pressure of compressed air. NMR spectra were recorded on an Avance 400 spectrometer (Billerica, US). The chemical shift data for each signal are given as δ in units of parts per million (ppm) relative to tetramethylsilane, where δ = 0.00 ppm and the standardised deuterated solvent peak (CDCl3 δ = 7.26 ppm). The number of protons (n) for a given resonance is indicated by nH. The multiplicity of each signal is indicated by s (singlet), br s (broad singlet), d (doublet), t (triplet), q (quartet), p (pentet), sep (septet), dd (doublet of doublets), ddd (doublet of doublet of doublets), dddd (doublet of doublet of doublet of doublets), dt (doublet of triplets), tt (triplet of triplets), dqd (doublet of quartets of doublets) or m (multiplet). Coupling constants (*J*) are quoted in Hz and calculated to the nearest 0.1 Hz. Additional experiments were performed to aid complete characterisation including: $^1$H-$^1$H homonuclear correlation spectroscopy (COSY), $^1$H-$^{13}$C heteronuclear correlation spectroscopy (HSQC) which was either proton coupled or proton decoupled, $^{31}$P proton coupled and $^{31}$P proton decoupled. Assignment of $^1$H and $^{13}$C atoms in NMR follows standard pyranose ring numbering.

Compound **4** was made from commercially available 6-azido-6-deoxyglucose as described before.[15]

**(N-(*R*)-[phenoxyphosphinyl]-L-alanine 1-methylethyl ester)-2,3,4-tri-O-acetyl-6-azido-6-deoxy-α-D-glucopyranoside (1) and (N-(*R*)-[phenoxyphosphinyl]-L-alanine 1-methylethyl ester)-2,3,4-tri-O-acetyl-6-azido-6-deoxy-β-D-glucopyranoside (2)**: Compound **4**[16] (180 mg, 0.54 mmol) was dissolved in anhydrous THF (3.2 mL) and cooled to -78 °C. A solution of 2 M LDA in THF (270 μL, 0.54 mmol) was added and the reaction was stirred for 15 mins before addition of N-[(*S*)-(2,3,4,5,6-Pentafluorophenoxy)phenoxyphosphinyl]-L-alanine 1-methylethyl ester (300 mg, 0.57 mmol) as a solution in THF (2.3 mL). The reaction was kept at -70 °C for 40 mins, then quenched by the addition of methanol (300 μL). After stirring for 5 mins, the solution was directly dry loaded on to silica for column chromatography using a gradient elution of 10% - 30% EtOAc in hexane with 1% $Et_3N$. The alpha anomer **1** was isolated (197 mg, 60% as a waxy white solid). Successive purifications of the beta anomer did not produce anomerically pure material and so preparative TLC was performed using 60% ethyl acetate in hexane with 1% $Et_3N$, yielding the beta anomer **2** (32 mg, 9.8%) as an oil with approx. 4% of the alpha-anomer **1** as a remaining impurity, as assessed by NMR. Alpha **1**: Rf = 0.61 (60% EtOAc in hexane with 1%

$Et_3N$) $^1$H NMR (400 MHz, $CDCl_3$) δ 7.37 - 7.32 (m, 2H ArH), 7.27 - 7.24 (m, 2H, ArH), 7.22 - 7.16 (m, 1H, ArH), 5.97 - 5.94 (dd, 1H, H1, *J* = 7.4, 3.4 Hz), 5.45 (t, 1H, H3, *J* = 10.3, 9.4 Hz), 5.11- 5.02 (m, 2H, H4, OC*H*($CH_3$)$_2$), 5.01 - 4.98 (ddd, 1H, H2, *J* = 10.3, 3.4, 2.5 Hz), 4.08 - 4.00 (m, 2H, H5, NHC*H*$CH_3$), 3.83 - 3.77 (m, 1H, NH), 3.30 - 3.26 (dd, 1H, H6a, *J* = 13.5, 2.9 Hz), 3.22 - 3.18 (dd, 1H, H6b, *J* = 13.5, 4.9 Hz) , 2.06 (s, 3H, $CH_3$), 2.04 (s, 3H, $CH_3$), 2.02 (s, 3H, $CH_3$), 1.45 - 1.43 (d,3H, NHCHC*H*$_3$, *J* = 7.0 Hz), 1.26 (d, 3H, OCH(*C*$H_3$)$_2$, *J* = 3.9 Hz), 1.25 (d, 3H, OCH(*C*$H_3$)$_2$, *J* = 3.9 Hz) .$^{13}$C NMR (101 MHz, $CDCl_3$) δ 172.8 (d, *C*OOCH($CH_3$)$_2$, *J* = 9.6 Hz), 170.2, 169.9, 169.5 (C=O), 150.62 (d, qArC, *J* = 7.0 Hz) , 130.0 (ArC), 125.3 (d, ArC, *J* = 1.3 Hz) , 120.3 (d, ArC, *J*= 5.0 Hz), 93.40 (d, C1, *J* = 5.6 Hz), 70.4 (C5), 69.9 (C2, *J* = 7.5 Hz), 69.7 (O*C*H($CH_3$)$_2$), 69.5 (C3), 68.9 (C4), 50.7 (C6), 50.4 (d, N*C*H$CH_3$, *J* = 2.4 Hz), 21.8 (d, CH(*C*$H_3$)$_2$, *J* = 3.2 Hz), 21.3, (d, NHCH*C*$H_3$, *J* = 3.7 Hz), 20.8 ($CH_3$), 20.7 ($CH_3$), 20.7 ($CH_3$). $^1$H-$^{13}$C-HSQC coupled: $^1J_{C\text{-}H}$ 180Hz. $^{31}$P NMR (162 MHz, $CDCl_3$) δ 1.11 - 0.94 (q, *J* = 8.8 Hz). HRMS $C_{24}H_{34}O_{12}N_4P_1$ [M+H$^+$] calc 601.1905 found: 601.1913 $[\alpha]_D$ = +91.9 (c = 1.0, $CHCl_3$), Reverse phase C18 HPLC retention time: 21.874 mins, 95.5% purity Beta **2**: Rf = 0.5 (60% ethyl acetate in hexane with 1% $Et_3N$) $^1$H NMR (400 MHz, $CDCl_3$) δ 7.33 - 7.29 (app t, 2H, ArH), 7.20 - 7.14 (m, 3H, ArH), 5.37 - 5.33 (t, H1, 1H, *J* = 7.8 Hz), 5.19 - 5.14 (m, 1H, H3), 5.09 - 5.02 (m, 3H, H2, H4, OC*H*($CH_3$)$_2$) , 4.08 - 4.00 (m, 1H, NHC*H*$CH_3$), 3.84 (dd, 1H, NH, *J* = 12.1, 8.6 Hz), 3.76 (ddd, 1H, H5, *J* = 9.9, 5.4, 2.8 Hz, 1H), 3.42 (dd, 1H, H6a, *J* = 13.5, 2.9 Hz), 3.33 (dd, 1H, H6b, *J* = 13.5, 5.4 Hz), 2.03 (s, 3H, COC*H*$_3$), 1.97 (s, 3H, COC*H*$_3$), 1.74 (s, 3H, COC*H*$_3$),1.42 (d, 3H, $CH_3$ *J* = 7.1 Hz,), 1.26 (d, 3H, OCH(C*H*$_3$)$_2$, *J* = 6.3 Hz), 1.24 (d, 3H, OCH(C*H*$_3$)$_2$, *J* = 6.2 Hz).$^{13}$C NMR (101 MHz, $CDCl_3$) δ 172.7 (d, *C*OOCH($CH_3$)$_2$, *J* = 10.1 Hz), 170.1, 169.5, 169.3 (CO), 150.6 (d, qArC, *J* = 7.3 Hz), 129.8 (ArCH), 125.3 (ArCH), 120.5 (d, ArCH, *J* = 4.8 Hz), 95.8 (d, C1, *J* = 4.5 Hz), 73.7 (C5), 72.4 (d, C3, *J* = 1.8 Hz), 71.0 (d, C2, *J* = 8.9 Hz), 69.5, 68.9 (C4, *C*H($CH_3$)$_2$), 50.8 (C6), 50.4 (d, NHCH, *J* = 2.2 Hz), 21.8 (OCH(*C*$H_3$)$_2$), 20.9 (d, NHCH*C*$H_3$, *J* = 3.6 Hz), 20.7, 20.6, 20.5 (CO*C*$H_3$). $^1$H-$^{13}$C-HSQC coupled: $^1J_{C\text{-}H}$ 168Hz. $^{31}$P NMR (162 MHz, $CDCl_3$) δ 1.38 - 1.20 (dt, *J* = 8.1Hz). HRMS: $C_{24}H_{34}O_{12}N_4P_1$ [M+H$^+$] calc 601.1905 found: 601.1909. $[\alpha]_D$ = +4 (c = 1.0, $CHCl_3$). Reverse phase C18 HPLC retention time: 21.858 mins, 98.4% purity.

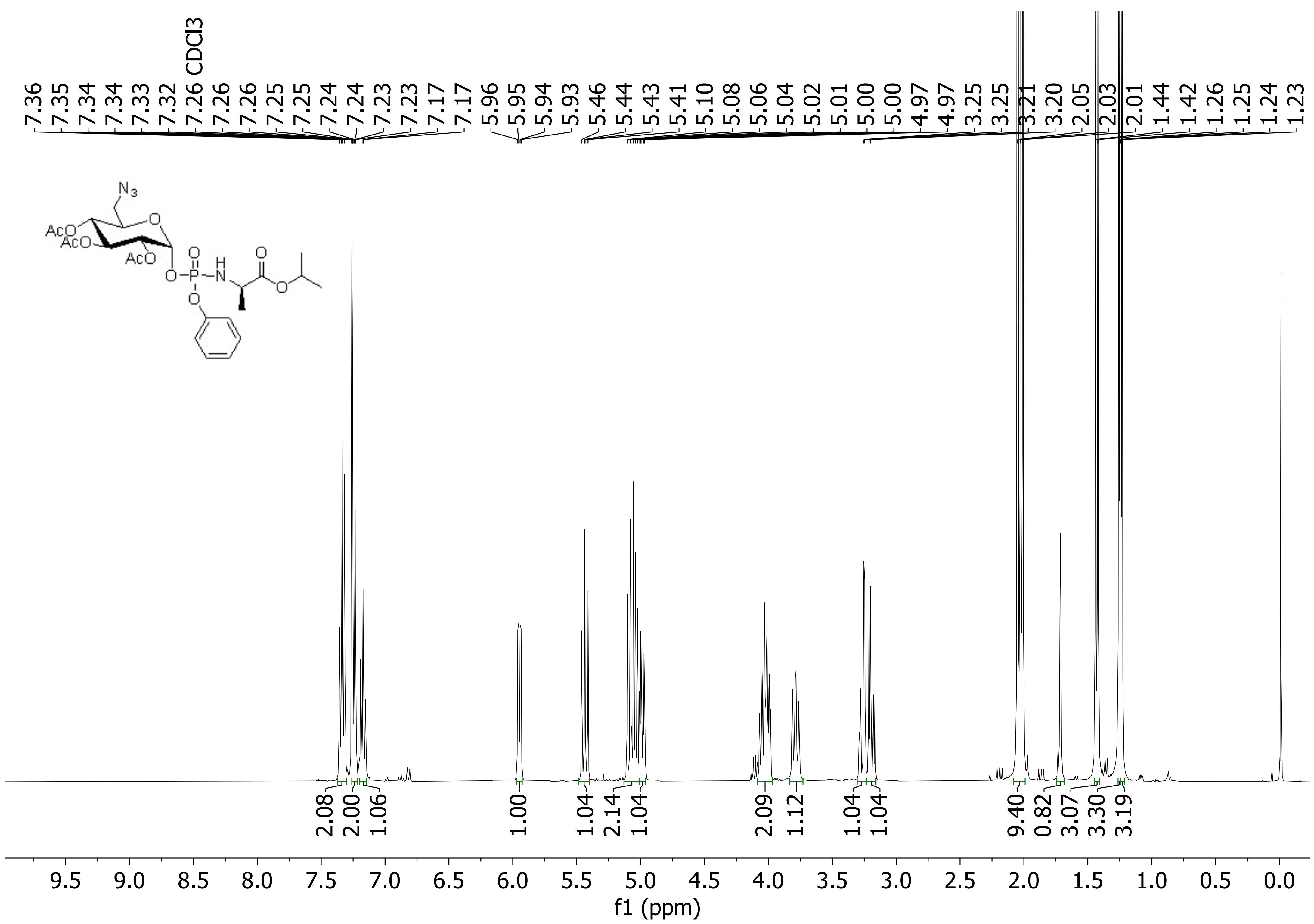
CDCl3
7.36 7.35 7.34 7.34 7.33 7.32 7.26 7.26 7.26 7.25 7.25 7.24 7.24 7.23 7.23 7.17 7.17 5.96 5.95 5.94 5.93 5.46 5.44 5.43 5.41 5.10 5.08 5.06 5.04 5.02 5.01 5.00 5.00 4.97 4.97 3.25 3.25 3.21 3.20 2.05 2.03 2.01 1.44 1.42 1.26 1.25 1.24 1.23
N3
AcO
AcO
AcO
O
O
P
H
N
2.08 2.00 1.06 1.00 1.04 2.14 1.04 2.09 1.12 1.04 1.04 9.40 0.82 3.07 3.30 3.19
9.5 9.0 8.5 8.0 7.5 7.0 6.5 6.0 5.5 5.0 4.5 4.0 3.5 3.0 2.5 2.0 1.5 1.0 0.5 0.0
f1 (ppm)

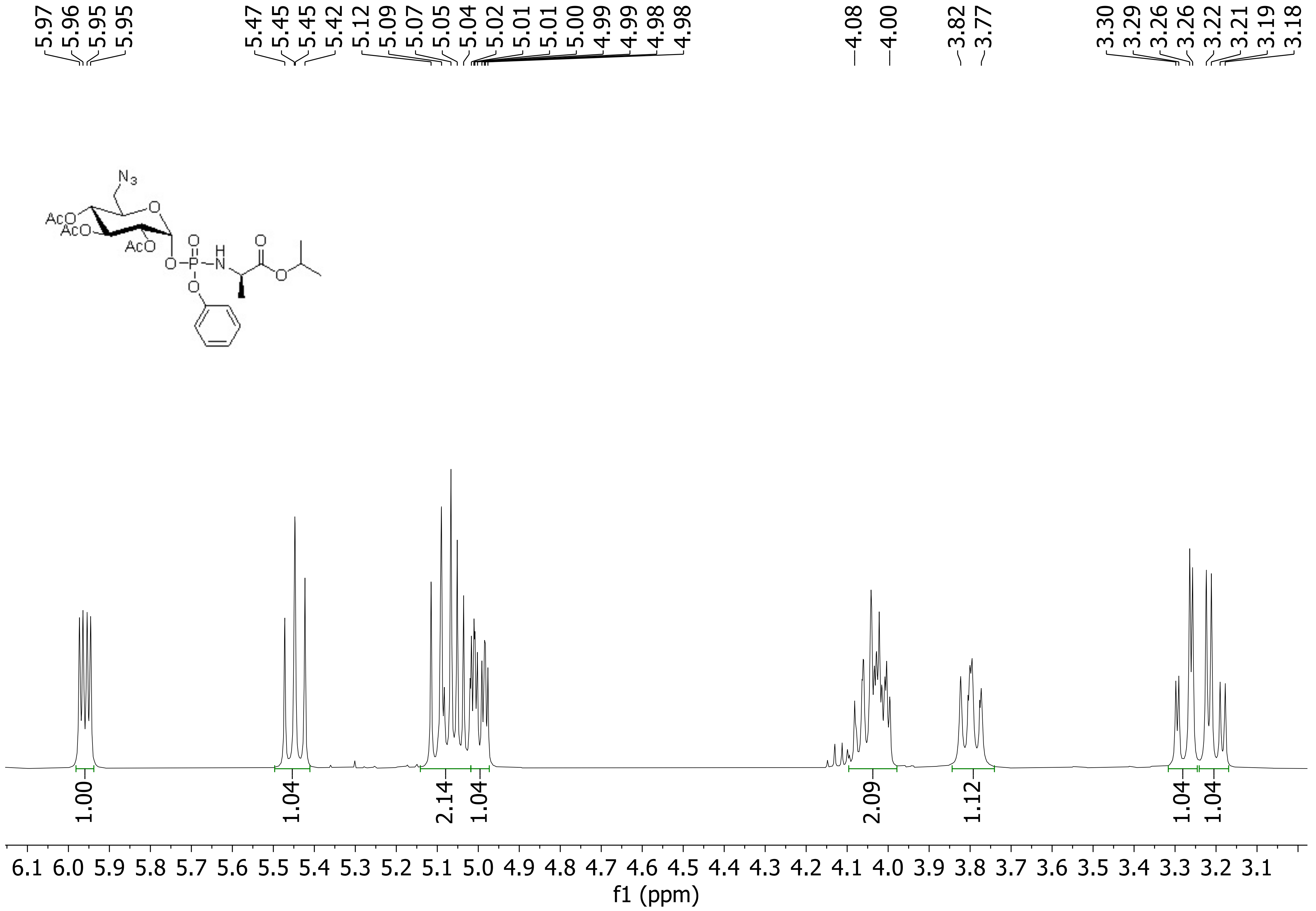
5.97 5.96 5.95 5.95
5.47 5.45 5.45 5.42 5.12 5.09 5.07 5.05 5.04 5.02 5.01 5.01 5.00 4.99 4.99 4.98 4.98
4.08 4.00
3.82 3.77
3.30 3.29 3.26 3.26 3.22 3.21 3.19 3.18
N3
AcO
AcO
AcO
O
O
O
P
H
N
O
O
O
O
1.00
1.04
2.14
1.04
2.09
1.12
1.04
1.04
6.1 6.0 5.9 5.8 5.7 5.6 5.5 5.4 5.3 5.2 5.1 5.0 4.9 4.8 4.7 4.6 4.5 4.4 4.3 4.2 4.1 4.0 3.9 3.8 3.7 3.6 3.5 3.4 3.3 3.2 3.1
f1 (ppm)

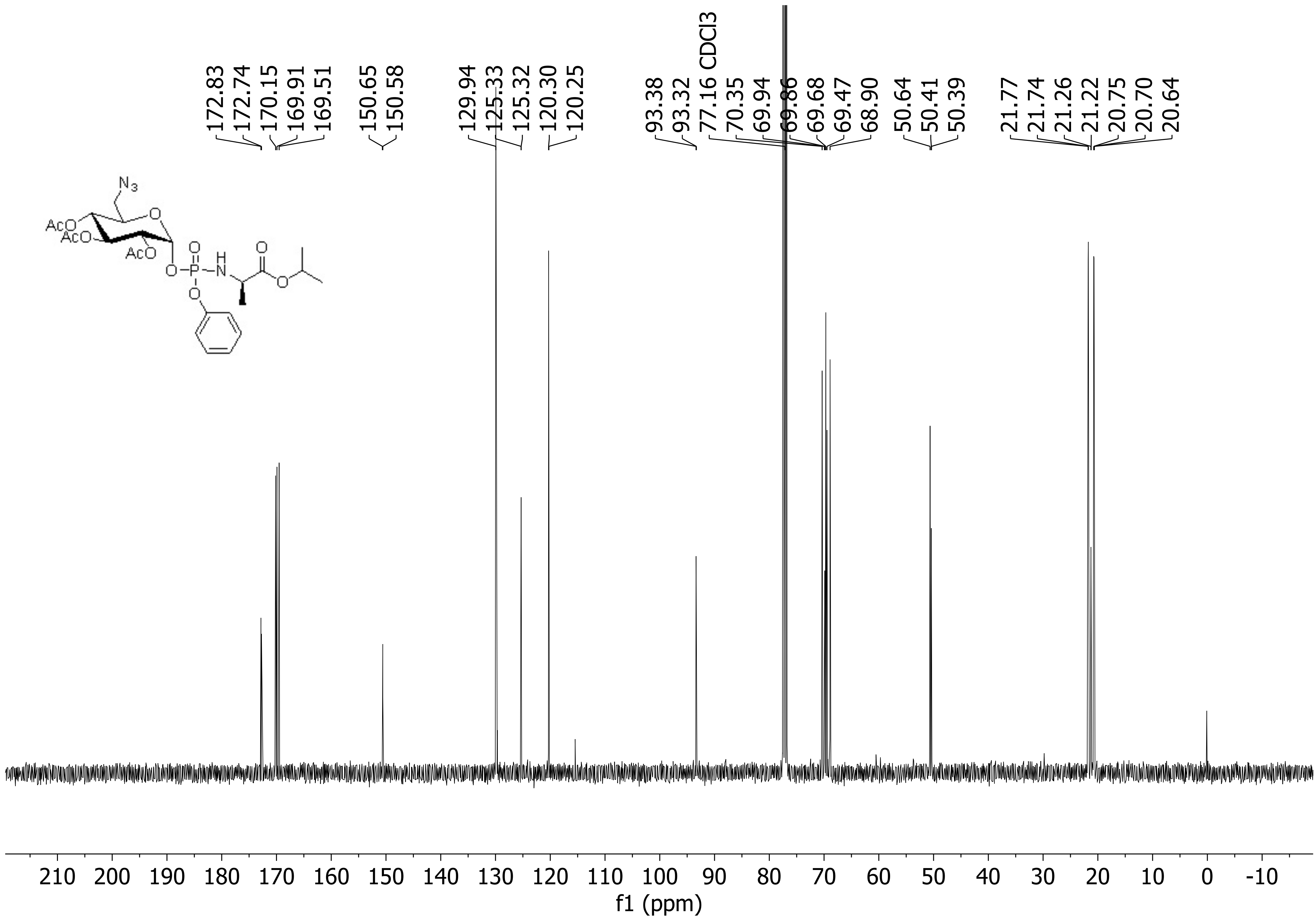

172.83
172.74
170.15
169.91
169.51
150.65
150.58
129.94
125.33
125.32
120.30
120.25
93.38
93.32
77.16 CDCl3
70.35
69.94
69.86
69.68
69.47
68.90
50.64
50.41
50.39
21.77
21.74
21.26
21.22
20.75
20.70
20.64
210 200 190 180 170 160 150 140 130 120 110 100 90 80 70 60 50 40 30 20 10 0 -10
f1 (ppm)

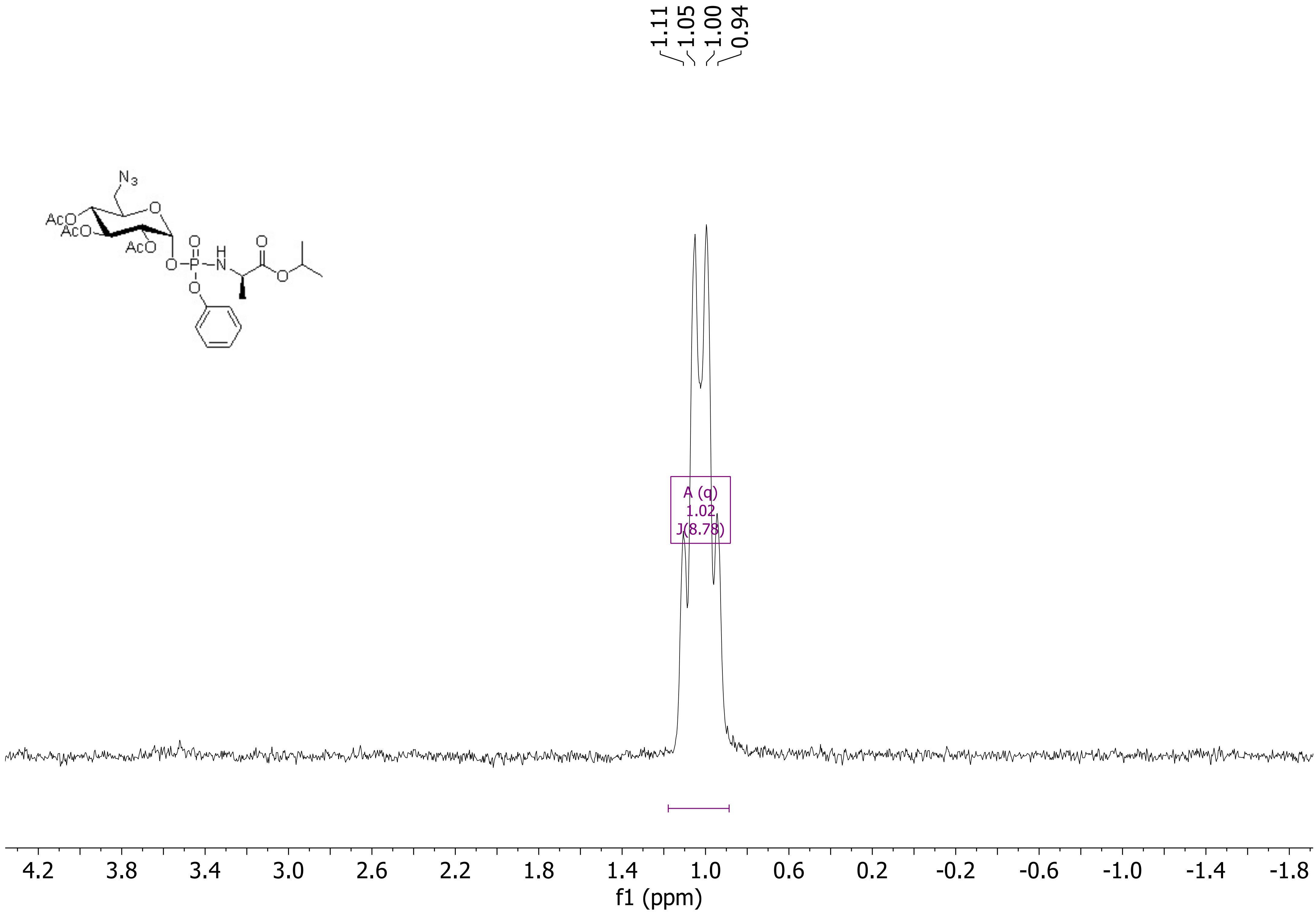
1.11
1.05
1.00
0.94
N3
AcO
AcO
AcO
O
O
H
O
P
N
O
O
O
A (q)
1.02
J(8.78)
4.2
3.8
3.4
3.0
2.6
2.2
1.8
1.4
1.0
0.6
0.2
-0.2
-0.6
-1.0
-1.4
-1.8
f1 (ppm)

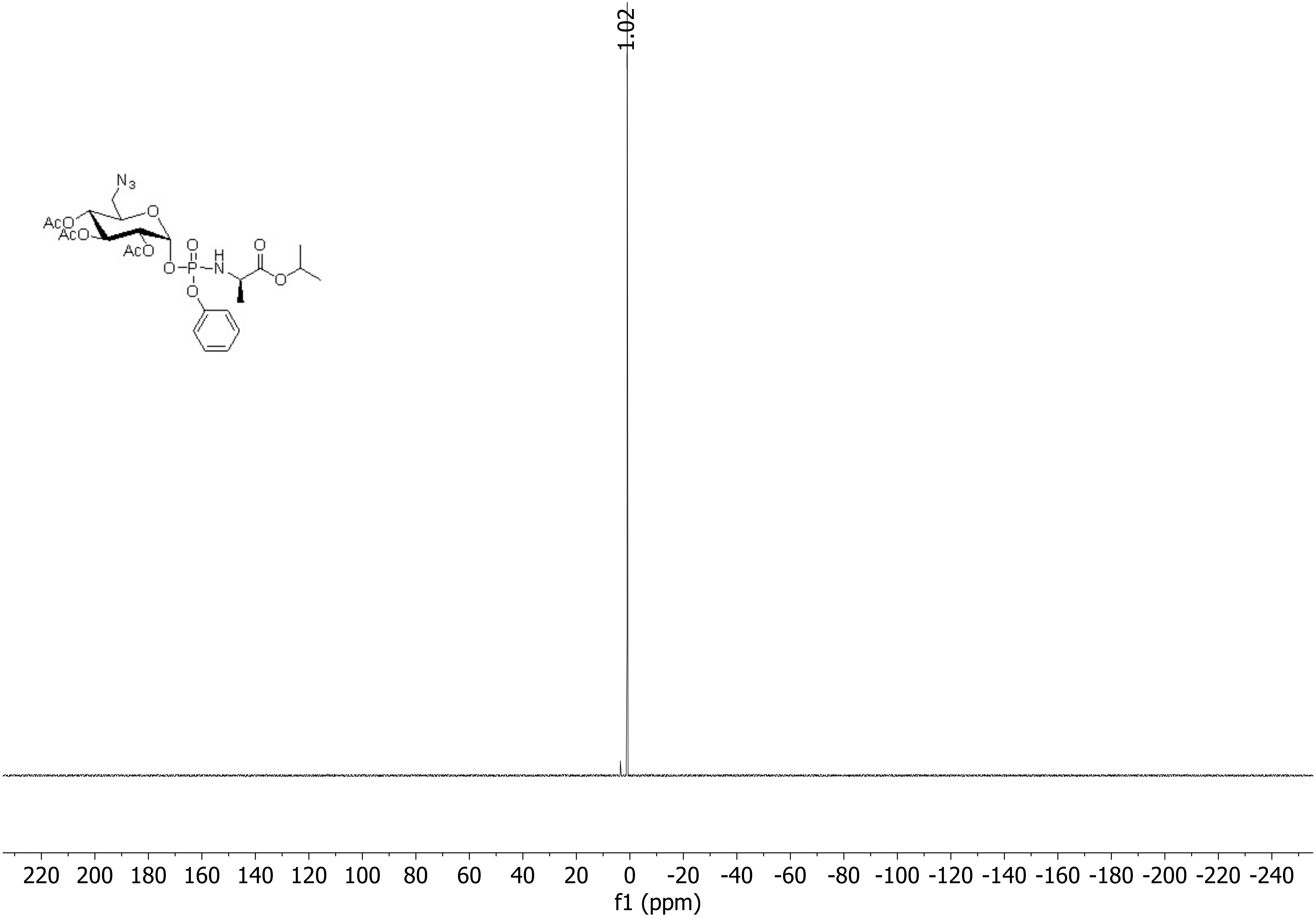

1.02
N3
AcO
AcO
AcO
O
O
P
H
N
O
O
O
O
220 200 180 160 140 120 100 80 60 40 20 0 -20 -40 -60 -80 -100 -120 -140 -160 -180 -200 -220 -240
f1 (ppm)

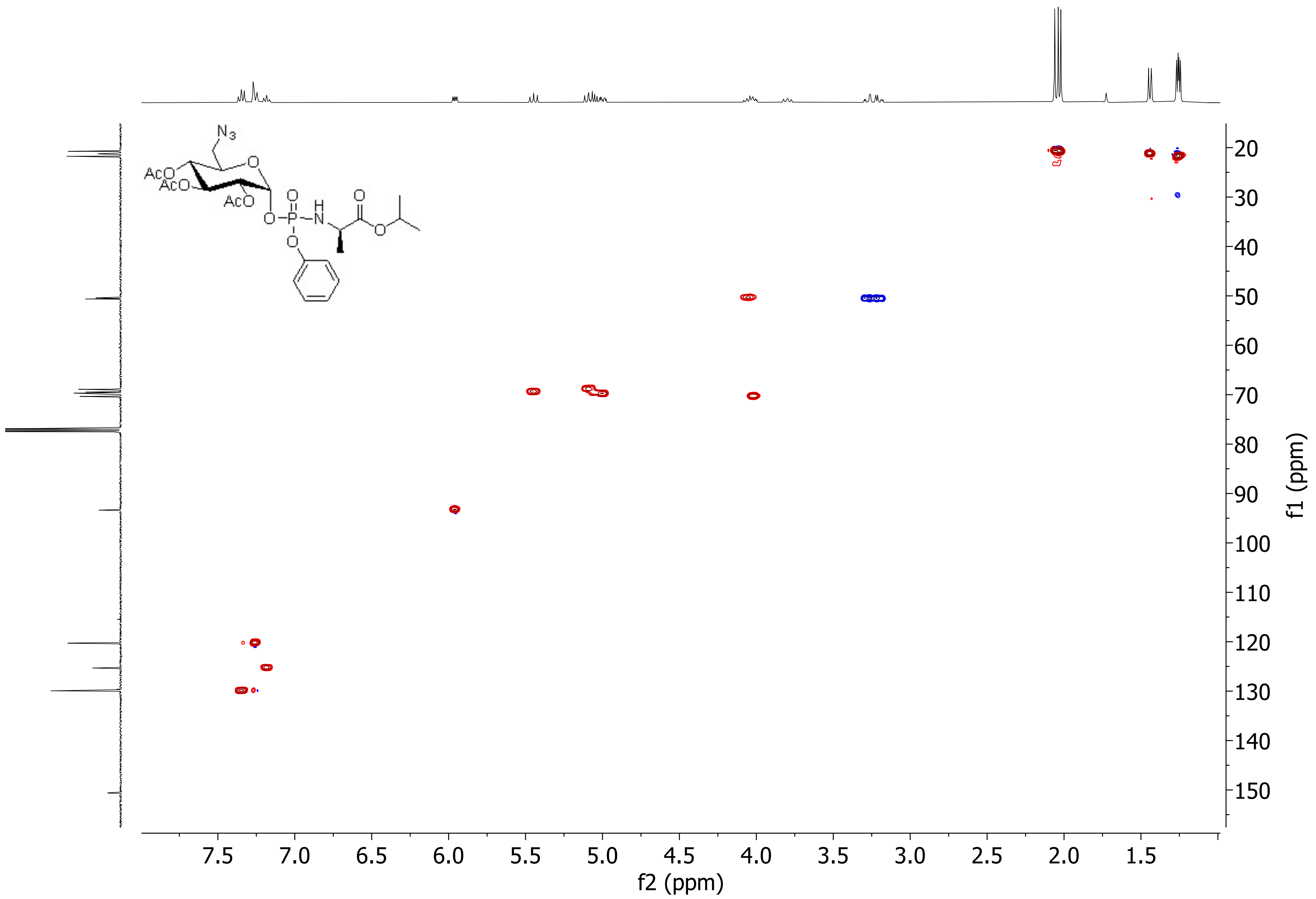
N3
AcO
AcO
AcO
O
O
P
H
N
O
O
O
O
20
30
40
50
60
70
80
90
100
110
120
130
140
150
f1 (ppm)
7.5
7.0
6.5
6.0
5.5
5.0
4.5
4.0
3.5
3.0
2.5
2.0
1.5
f2 (ppm)

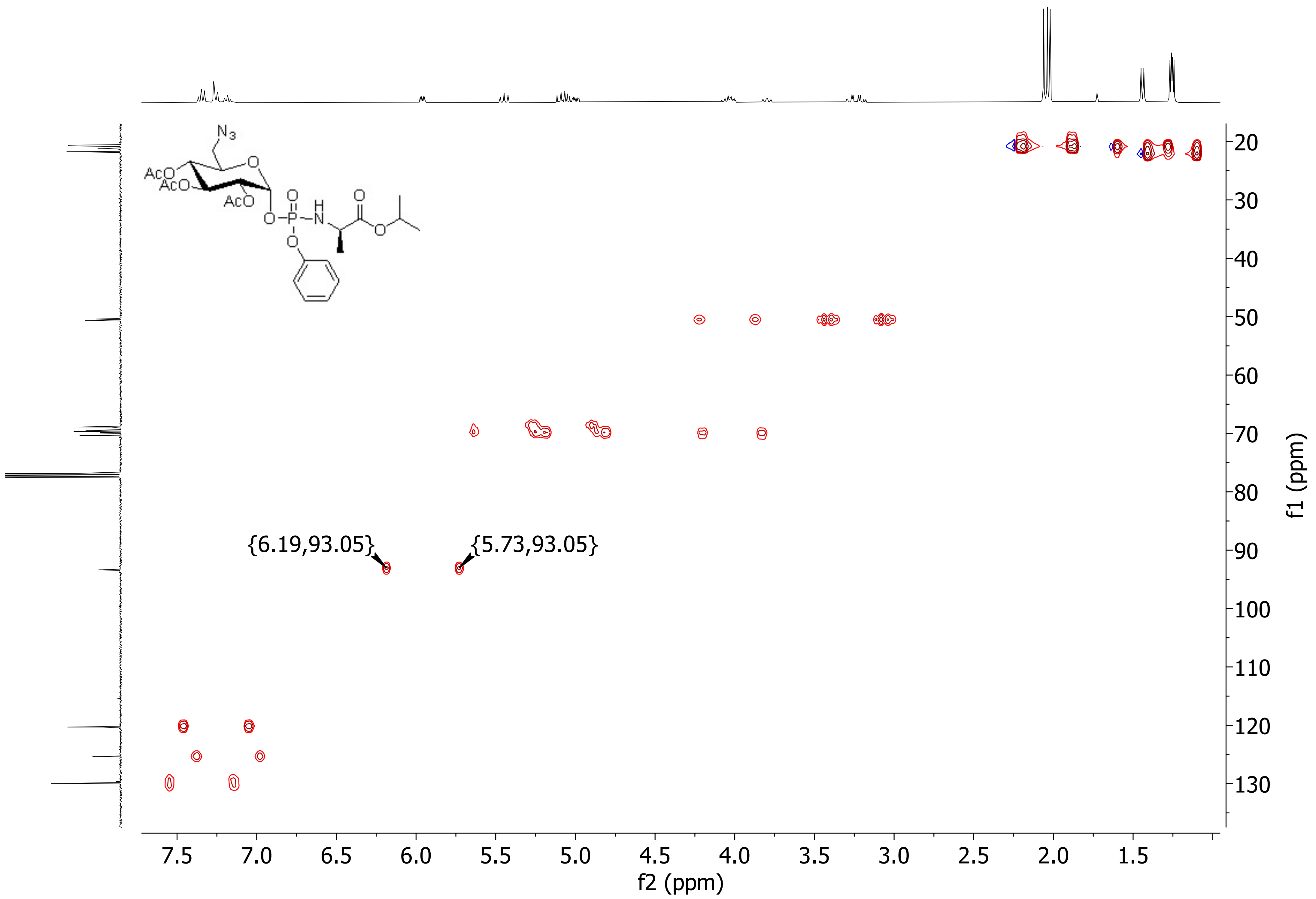
N3
AcO
AcO
AcO
O
O
P
H
N
O
O
O
O
{6.19,93.05}
{5.73,93.05}
20
30
40
50
60
70
80
90
100
110
120
130
f1 (ppm)
7.5
7.0
6.5
6.0
5.5
5.0
4.5
4.0
3.5
3.0
2.5
2.0
1.5
f2 (ppm)

ANC BS 5 F2 MW=600? C24H33N4O12P
(DCM)/MeOH+NH4OAc

NMSF, Swansea University
LTQ Orbitrap XL

Gavin Miller
01/03/2023 10:55:02

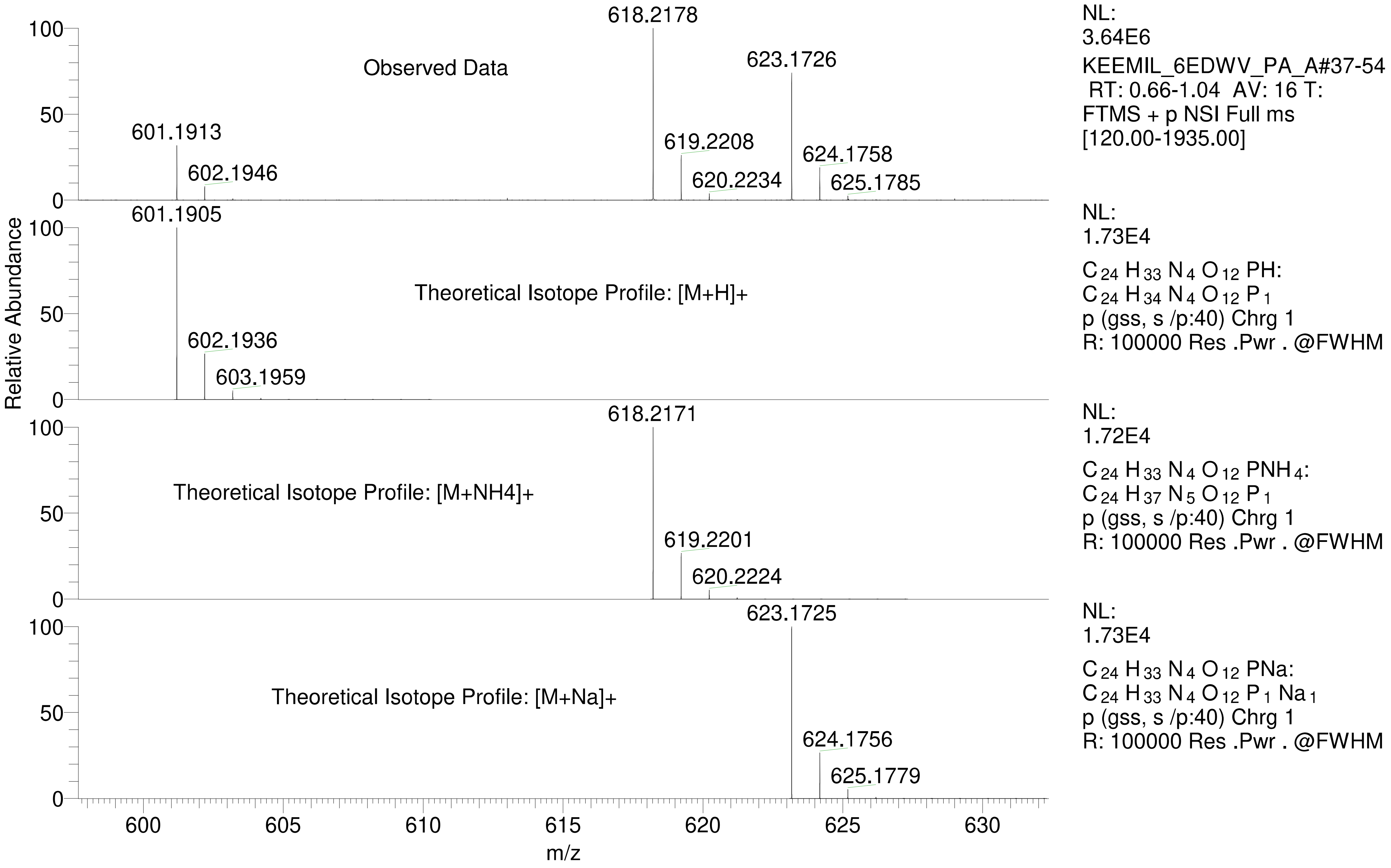

| Isotope: | Min. .. Max. | | |
|---|---|---|---|
| 14 N | 0....16 | | |
| 16 O | 0....20 | | |
| 12 C | 0....100 | | |
| 1 H | 0....120 | | |
| 23 Na | 1....1 | | |
| 31 P | 0....2 | | |
| Tolerance Window: | +- 5.00 ppm | | |
| Db/Ring Equiv: | -10.. 500 | N-Rule: | Do not use |
| Fits: | 500 | Charge: | 1 |

| Mass | Theoretical Mass | Delta [ppm] | RDB | Composition |
|---|---|---|---|---|
| 601.1913 | 601.1914 | -0.1 | -6.5 | $C_6 H_{39} O_{18} N_{10} P_2$ |
| | 601.1914 | -0.1 | 28.0 | $C_{39} H_{28} O_2 N_3 P_1$ |
| | 601.1911 | 0.4 | 32.5 | $C_{43} H_{25} O_2 N_2$ |
| | 601.1910 | 0.4 | -2.0 | $C_{10} H_{36} O_{18} N_9 P_1$ |
| | 601.1916 | -0.4 | 25.5 | $C_{28} H_{21} O_3 N_{14}$ |
| | 601.1910 | 0.4 | 3.5 | $C_9 H_{30} O_{13} N_{16} P_1$ |
| | 601.1916 | -0.4 | 20.0 | $C_{29} H_{27} O_8 N_7$ |
| | 601.1916 | -0.4 | 14.5 | $C_{30} H_{33} O_{13}$ |
| | 601.1917 | -0.6 | 23.5 | $C_{35} H_{31} O_2 N_4 P_2$ |
| | 601.1908 | 0.8 | 6.0 | $C_{20} H_{37} O_{12} N_5 P_2$ |
| | 601.1908 | 0.8 | 11.5 | $C_{19} H_{31} O_7 N_{12} P_2$ |
| | 601.1907 | 0.9 | 2.5 | $C_{14} H_{33} O_{18} N_8$ |
| | 601.1919 | -0.9 | 21.0 | $C_{24} H_{24} O_3 N_{15} P_1$ |
| | 601.1907 | 1.0 | 8.0 | $C_{13} H_{27} O_{13} N_{15}$ |
| | 601.1919 | -1.0 | 15.5 | $C_{25} H_{30} O_8 N_8 P_1$ |
| | 601.1919 | -1.0 | 10.0 | $C_{26} H_{36} O_{13} N_1 P_1$ |
| | 601.1905 | 1.3 | 10.5 | $C_{24} H_{34} O_{12} N_4 P_1$ |
| | 601.1905 | 1.3 | 16.0 | $C_{23} H_{28} O_7 N_{11} P_1$ |
| | 601.1921 | -1.3 | 7.5 | $C_{15} H_{29} O_{14} N_{12}$ |
| | 601.1921 | -1.3 | 2.0 | $C_{16} H_{35} O_{19} N_5$ |
| | 601.1922 | -1.5 | 16.5 | $C_{20} H_{27} O_3 N_{16} P_2$ |
| | 601.1922 | -1.5 | 11.0 | $C_{21} H_{33} O_8 N_9 P_2$ |
| | 601.1922 | -1.5 | 5.5 | $C_{22} H_{39} O_{13} N_2 P_2$ |
| | 601.1903 | 1.6 | 18.5 | $C_{34} H_{35} O_6 P_2$ |
| | 601.1903 | 1.6 | 24.0 | $C_{33} H_{29} O_1 N_7 P_2$ |
| | 601.1902 | 1.8 | 15.0 | $C_{28} H_{31} O_{12} N_3$ |
| | 601.1902 | 1.8 | 20.5 | $C_{27} H_{25} O_7 N_{10}$ |
| | 601.1924 | -1.8 | 3.0 | $C_{11} H_{32} O_{14} N_{13} P_1$ |
| | 601.1924 | -1.8 | -2.5 | $C_{12} H_{38} O_{19} N_6 P_1$ |
| | 601.1900 | 2.1 | 28.5 | $C_{37} H_{26} O_1 N_6 P_1$ |
| | 601.1900 | 2.1 | -6.0 | $C_4 H_{37} O_{17} N_{13} P_2$ |

=====================================================================

Acq. Operator : SYSTEM Seq. Line : 1
Sample Operator : SYSTEM
Acq. Instrument : Prep LC Location : P2-A-02
Injection Date : 10/3/2023 2:47:58 PM Inj : 1
Inj Volume : 200.000 µl
Method : C:\Users\Public\Documents\ChemStation\1\Data\Aisling\ANC BS6 iii percent purity repeat 2023-10-03 14-46-55\ANC-BS-6iii Percent Purity.M (Sequence Method)
Last changed : 10/3/2023 2:46:52 PM by SYSTEM
Method Info : Polaris 5 C15-A 250 x10mm SN 593740

Additional Info : Peak(s) manually integrated

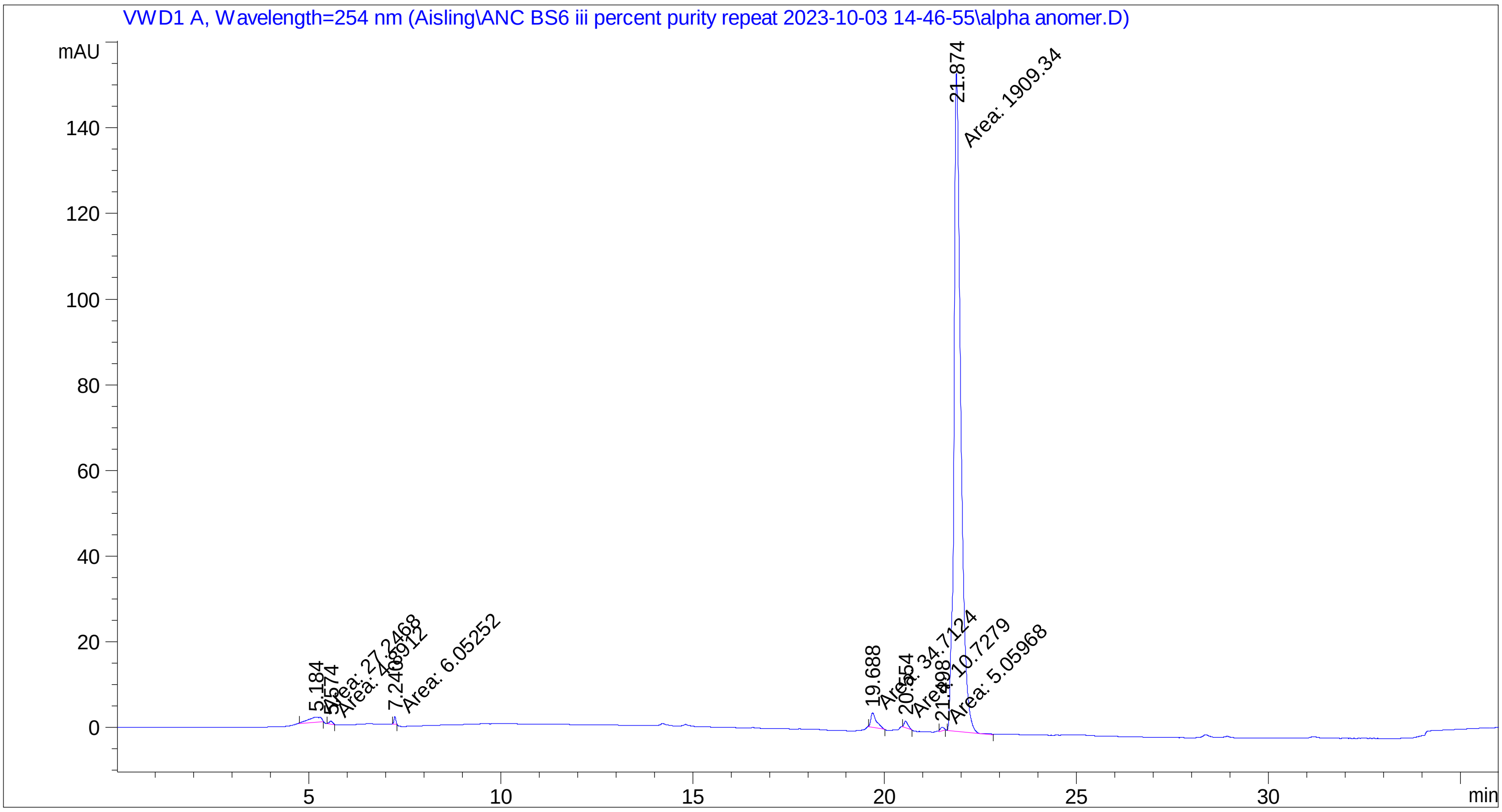


=====================================================================

Fraction Information

=====================================================================

No Fractions found.

=====================================================================

=====================================================================

=====================================================================

Area Percent Report

=====================================================================

Sorted By : Signal
Multiplier : 1.0000
Dilution : 1.0000
Use Multiplier & Dilution Factor with ISTDs

Signal 1: VWD1 A, Wavelength=254 nm

| Peak # | RetTime [min] | Type | Width [min] | Area [mAU*s] | Height [mAU] | Area % |
|---|---|---|---|---|---|---|
| 1 | 5.184 | MM | 0.3947 | 27.24684 | 1.15067 | 1.3637 |
| 2 | 5.574 | MM | 0.1148 | 4.89120 | 7.10091e-1 | 0.2448 |
| 3 | 7.240 | MM | 0.0557 | 6.05252 | 1.81123 | 0.3029 |
| 4 | 19.688 | MM | 0.1701 | 34.71244 | 3.40032 | 1.7373 |
| 5 | 20.554 | MM | 0.1191 | 10.72786 | 1.50101 | 0.5369 |
| 6 | 21.498 | MM | 0.1049 | 5.05968 | 8.03798e-1 | 0.2532 |
| 7 | 21.874 | MM | 0.2072 | 1909.33606 | 153.56323 | 95.5611 |
| Totals : | | | | 1998.02659 | 162.94035 | |

=====================================================================

*** End of Report ***

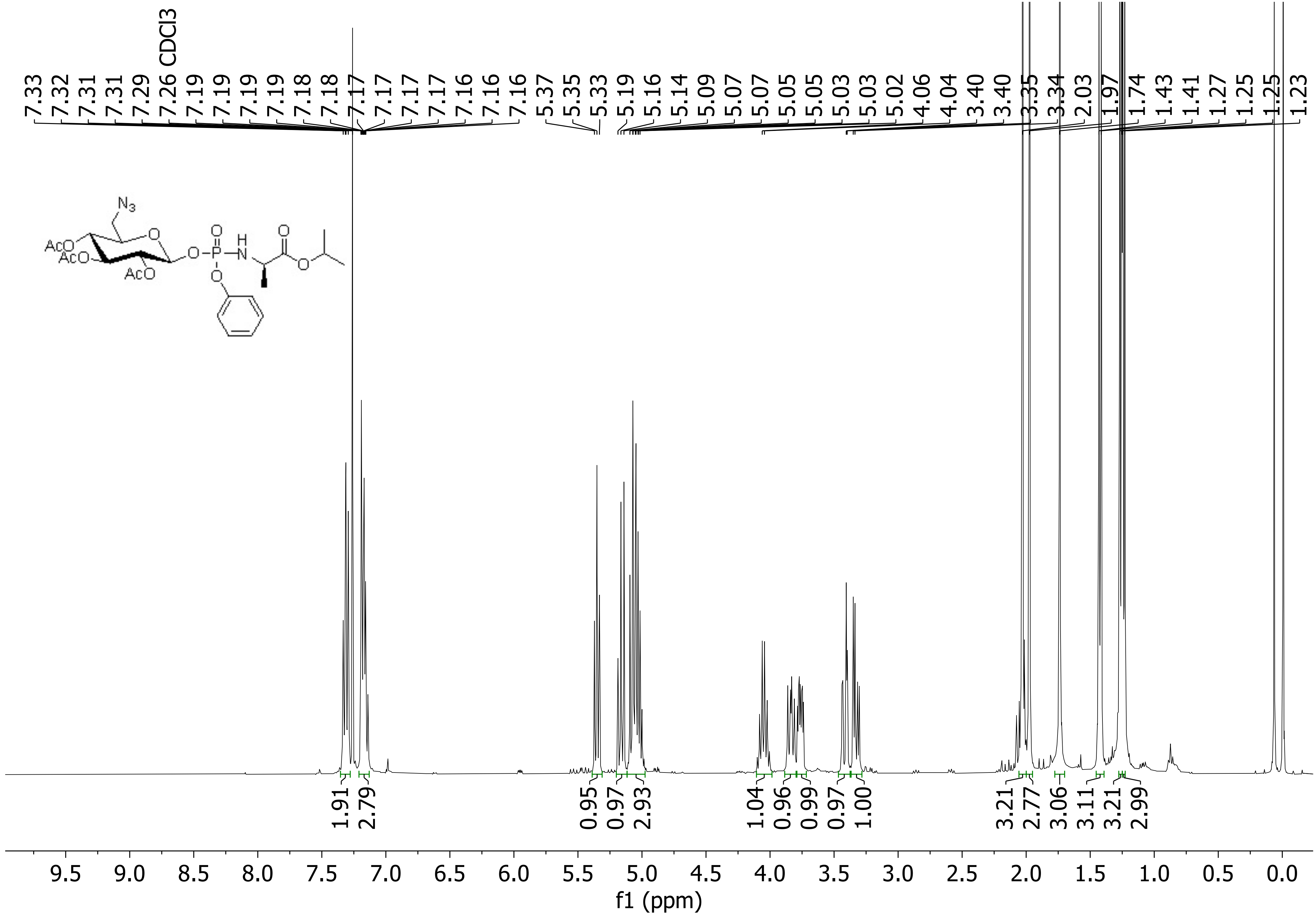

CDCl3
7.33 7.32 7.31 7.31 7.29 7.26 7.19 7.19 7.19 7.19 7.18 7.18 7.17 7.17 7.17 7.17 7.16 7.16
5.37 5.35 5.33 5.19 5.16 5.14 5.09 5.07 5.07 5.05 5.05 5.03 5.03 5.02
4.06 4.04 3.40 3.40 3.35 3.34
2.03 1.97 1.74 1.43 1.41 1.27 1.25 1.25 1.23
N3
AcO
AcO
AcO
O
H
N
1.91 2.79 0.95 0.97 2.93 1.04 0.96 0.99 0.97 1.00 3.21 2.77 3.06 3.11 3.21 2.99
9.5 9.0 8.5 8.0 7.5 7.0 6.5 6.0 5.5 5.0 4.5 4.0 3.5 3.0 2.5 2.0 1.5 1.0 0.5 0.0
f1 (ppm)

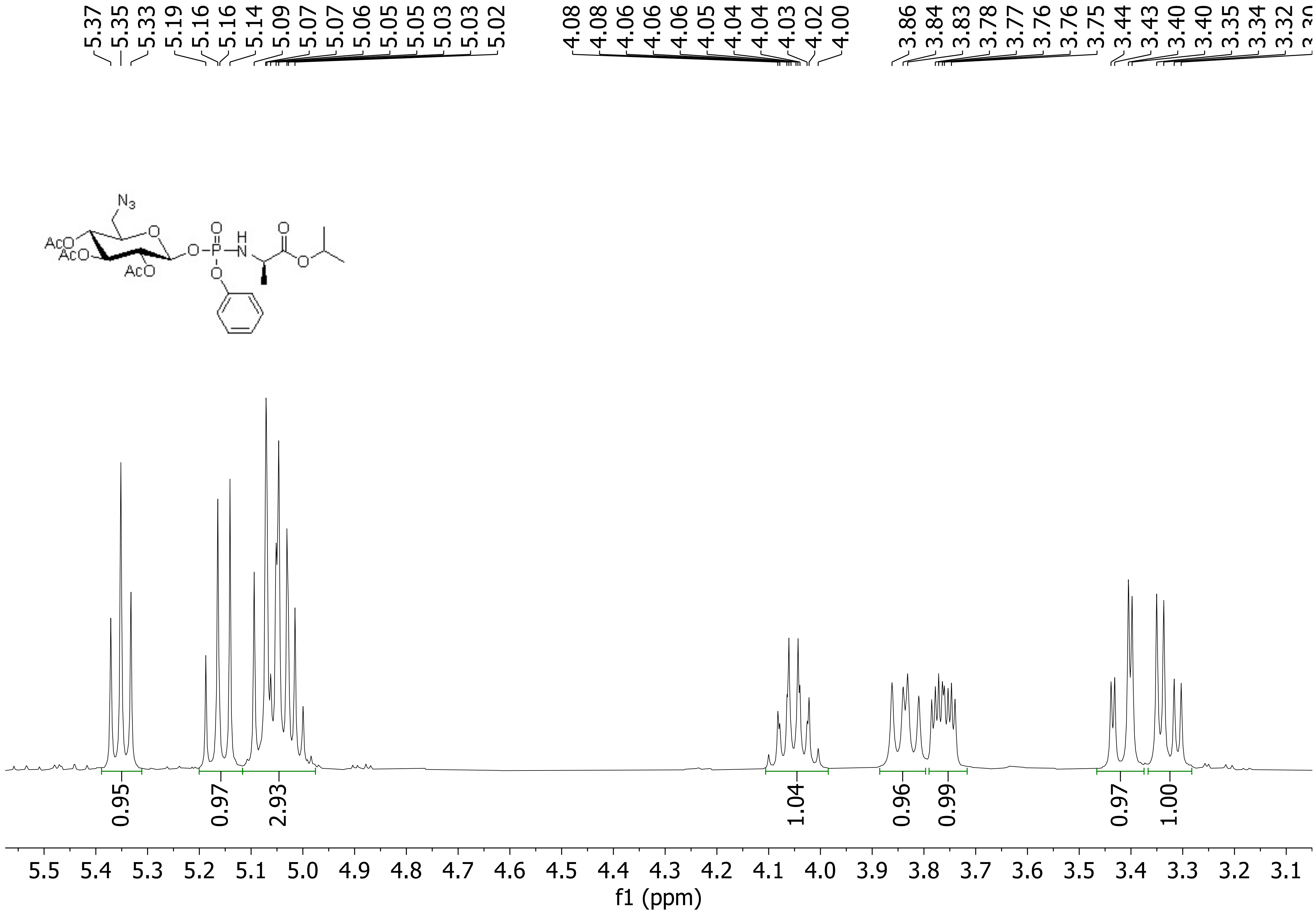
5.37 5.35 5.33 5.19 5.16 5.16 5.14 5.09 5.07 5.07 5.06 5.05 5.05 5.03 5.03 5.02
4.08 4.08 4.06 4.06 4.06 4.05 4.04 4.04 4.03 4.02 4.00
3.86 3.84 3.83 3.78 3.77 3.76 3.76 3.75 3.44 3.43 3.40 3.40 3.35 3.34 3.32
$N_3$
O
AcO
AcO
AcO
O
P
H
N
O
O
O
0.95
0.97
2.93
1.04
0.96
0.99
0.97
1.00
5.5 5.4 5.3 5.2 5.1 5.0 4.9 4.8 4.7 4.6 4.5 4.4 4.3 4.2 4.1 4.0 3.9 3.8 3.7 3.6 3.5 3.4 3.3 3.2 3.1
f1 (ppm)

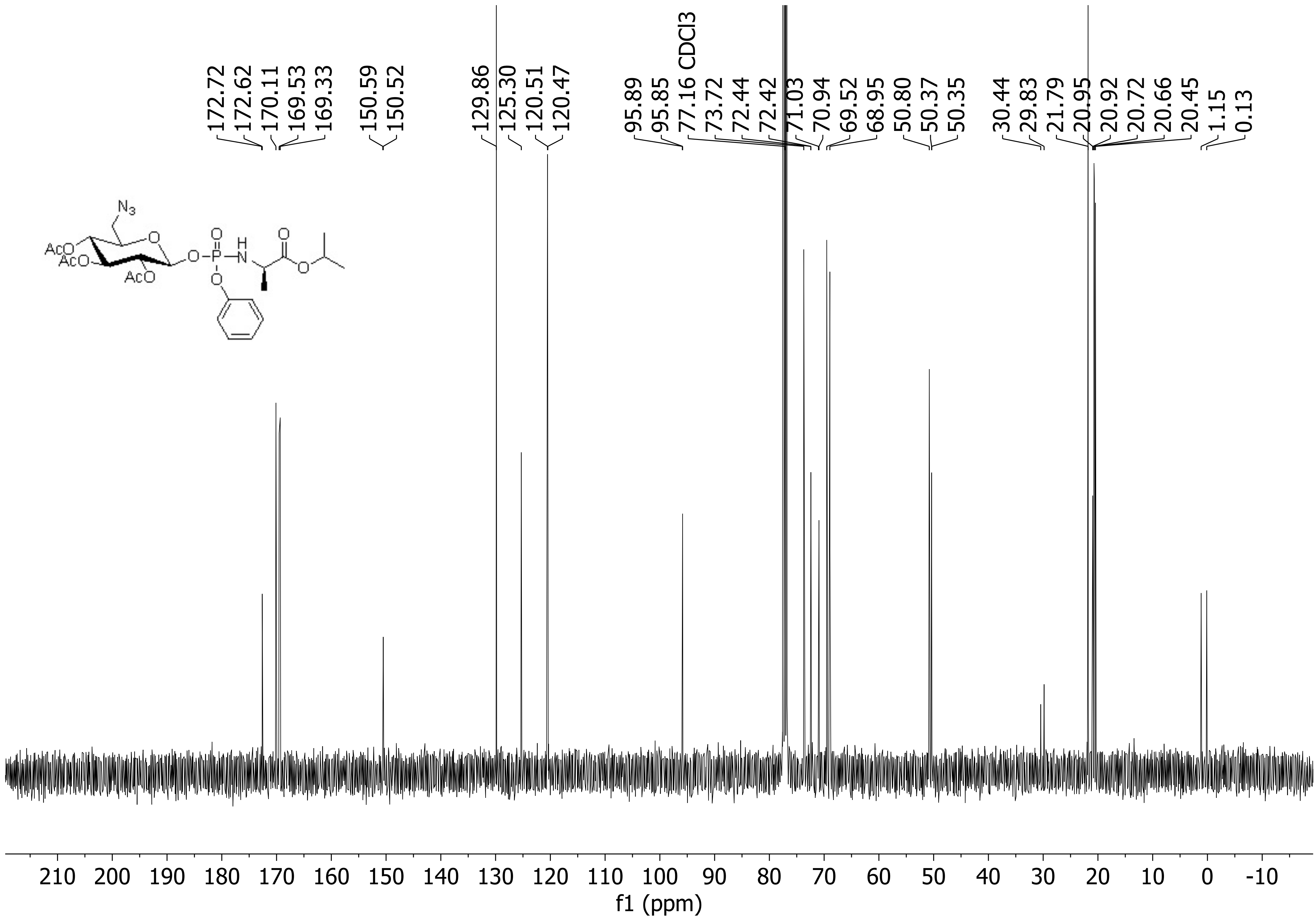

172.72
172.62
170.11
169.53
169.33
150.59
150.52
129.86
125.30
120.51
120.47
95.89
95.85
77.16 CDCl3
73.72
72.44
72.42
71.03
70.94
69.52
68.95
50.80
50.37
50.35
30.44
29.83
21.79
20.95
20.92
20.72
20.66
20.45
1.15
0.13
210 200 190 180 170 160 150 140 130 120 110 100 90 80 70 60 50 40 30 20 10 0 -10
f1 (ppm)

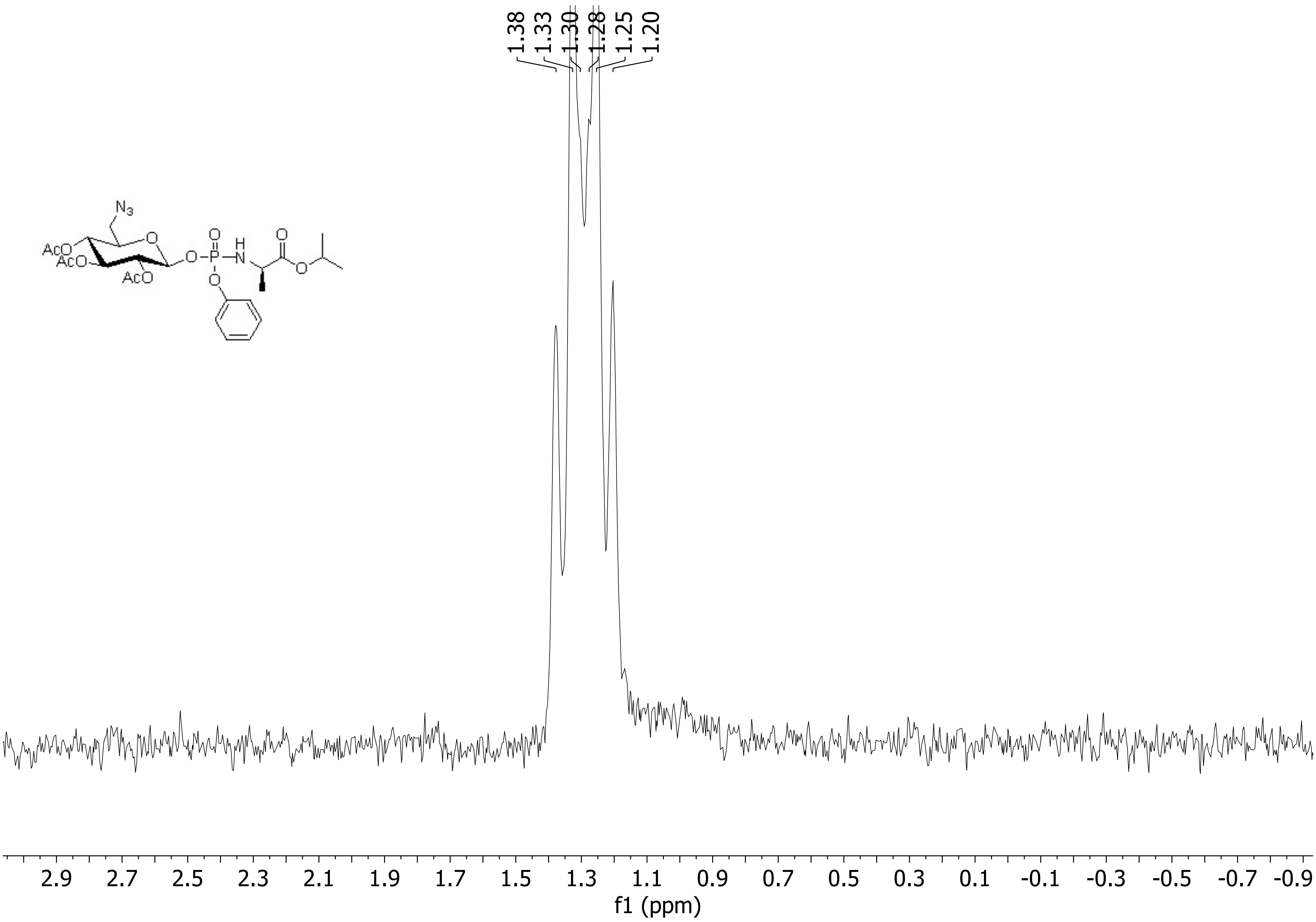
1.38
1.33
1.30
1.28
1.25
1.20
N3
AcO
AcO
AcO
O
O
P
H
N
O
O
O
2.9
2.7
2.5
2.3
2.1
1.9
1.7
1.5
1.3
1.1
0.9
0.7
0.5
0.3
0.1
-0.1
-0.3
-0.5
-0.7
-0.9
f1 (ppm)

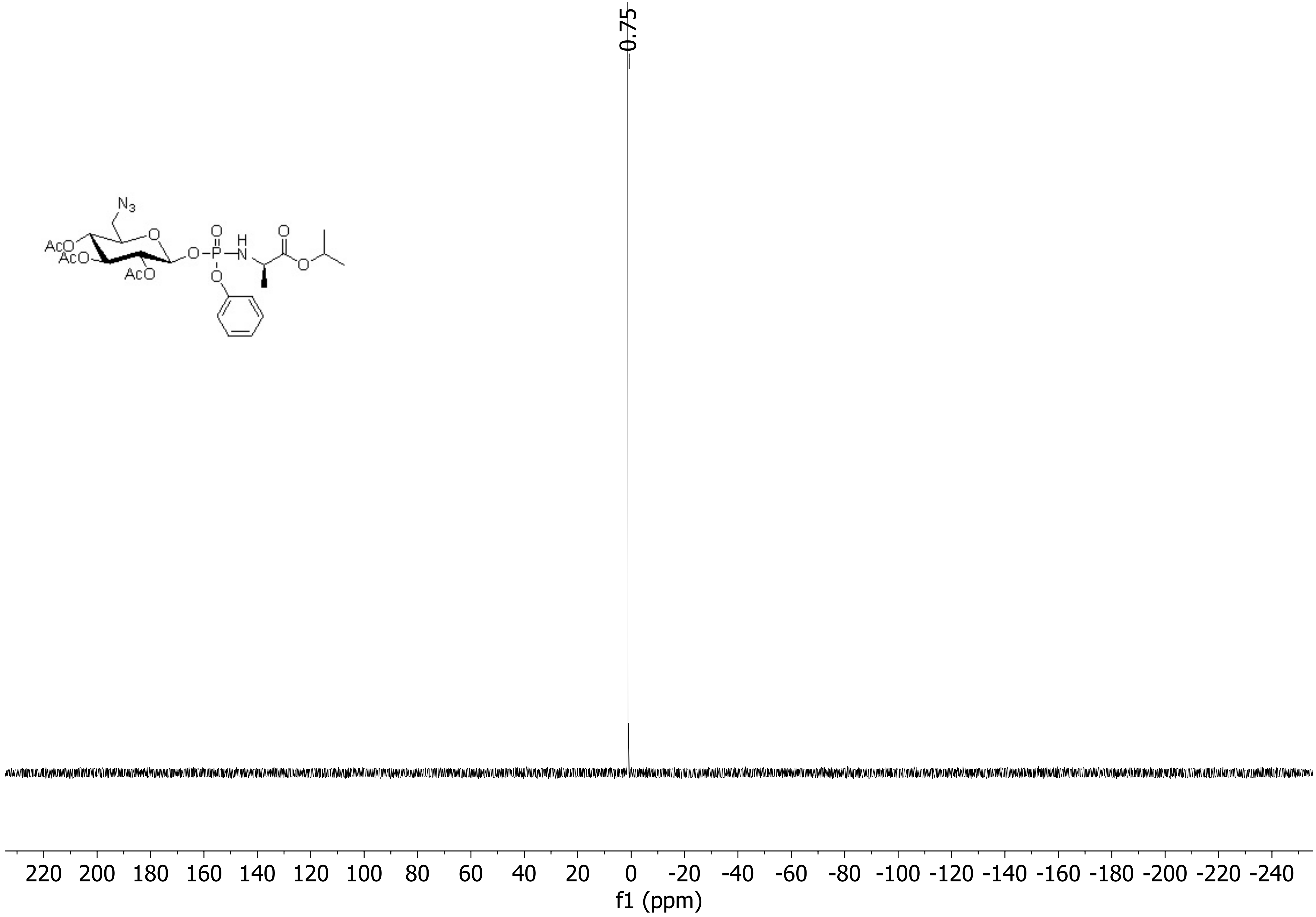

-0.75
N3
AcO
AcO
AcO
O
O
P
O
H
N
O
O
O
220 200 180 160 140 120 100 80 60 40 20 0 -20 -40 -60 -80 -100 -120 -140 -160 -180 -200 -220 -240
f1 (ppm)

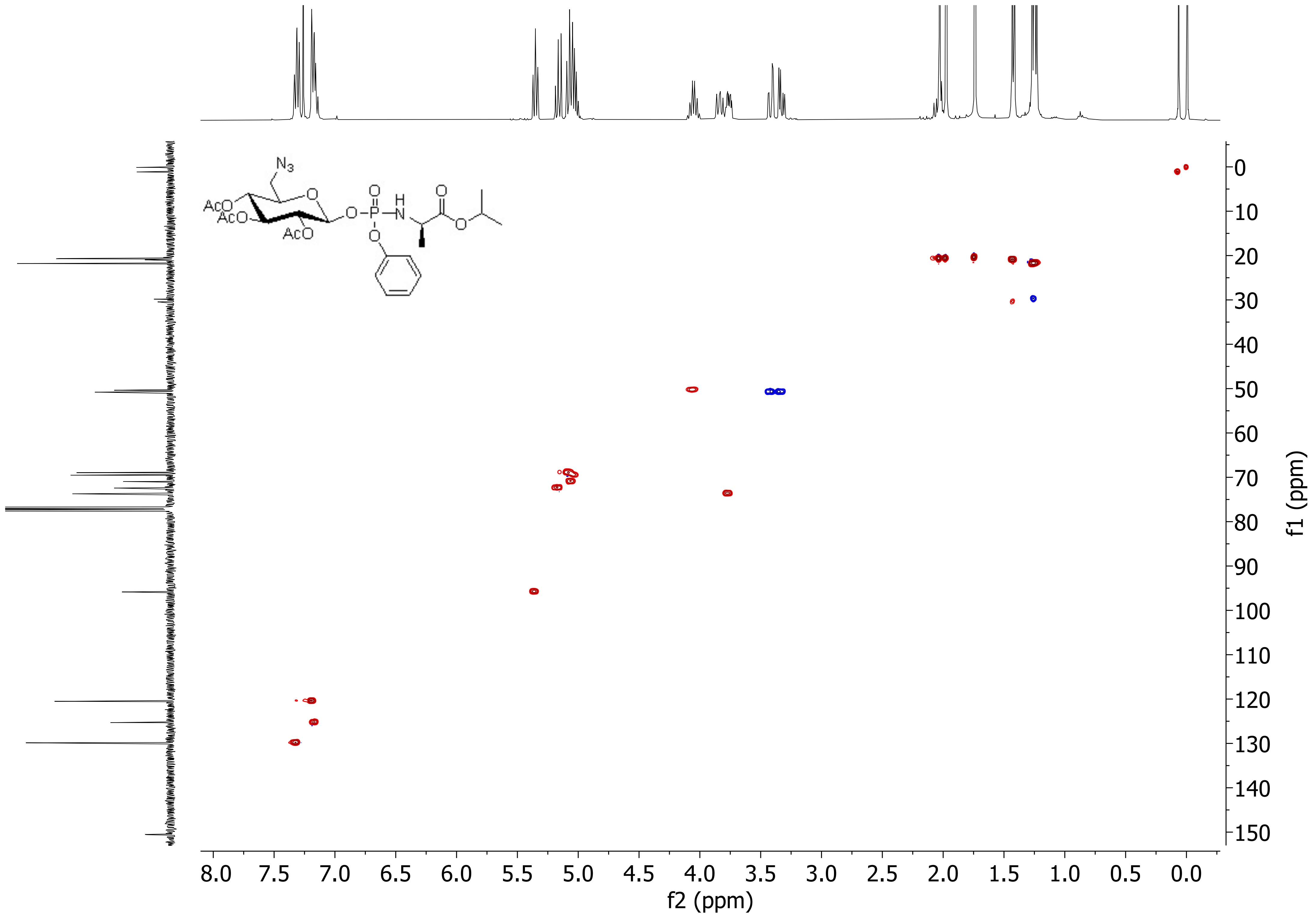
N3
AcO
AcO
AcO
O
P
H
N
f2 (ppm)
f1 (ppm)
8.0
7.5
7.0
6.5
6.0
5.5
5.0
4.5
4.0
3.5
3.0
2.5
2.0
1.5
1.0
0.5
0.0
0
10
20
30
40
50
60
70
80
90
100
110
120
130
140
150

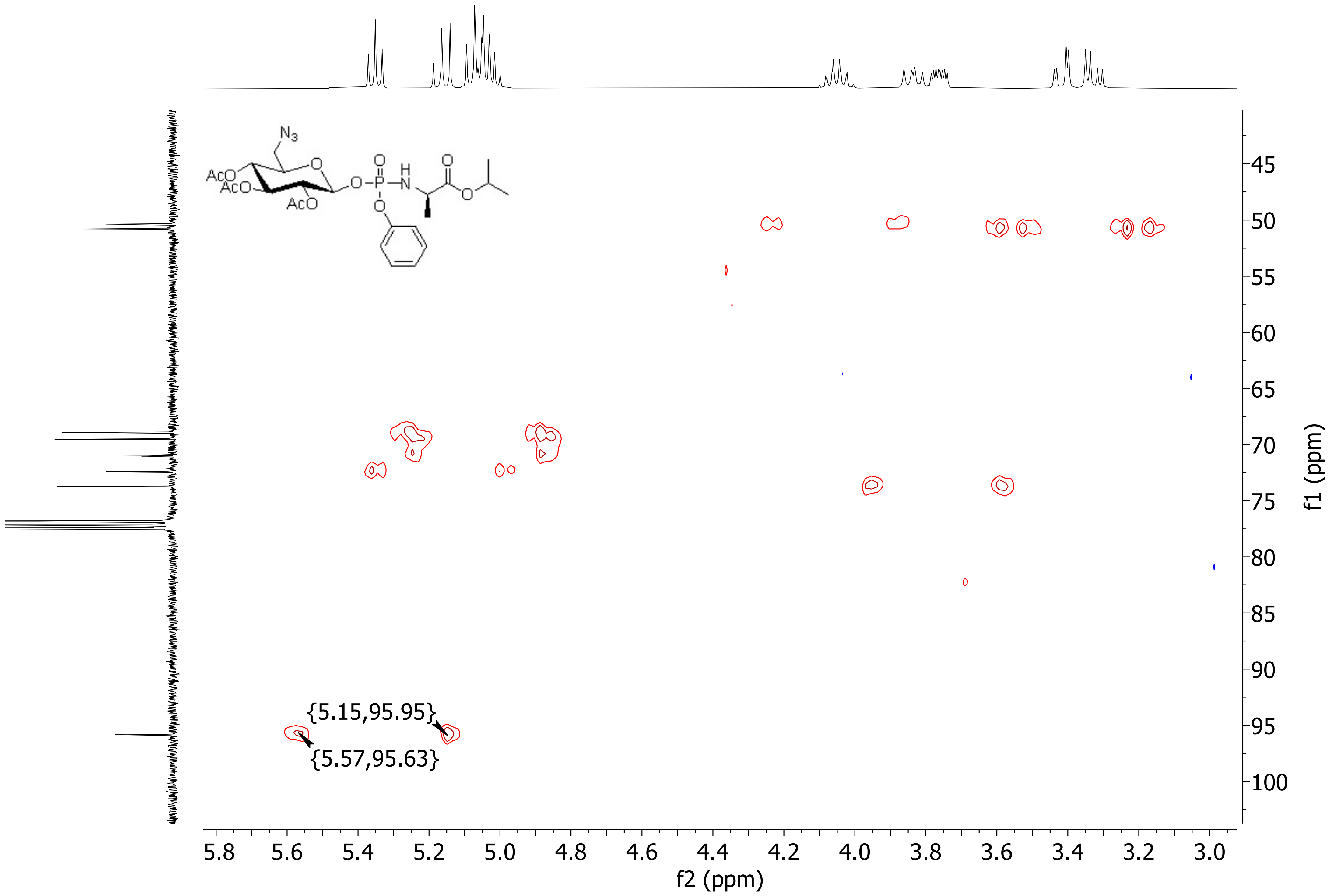
N3
AcO
AcO
AcO
O
O-P-N
H
O
O
{5.15,95.95}
{5.57,95.63}
f2 (ppm)
f1 (ppm)

ANC BS 5 F3 MW=600? C24H33N4O12P
(DCM)/MeOH+NH4OAc

NMSF, Swansea University
LTQ Orbitrap XL

Gavin Miller
01/03/2023 10:58:09

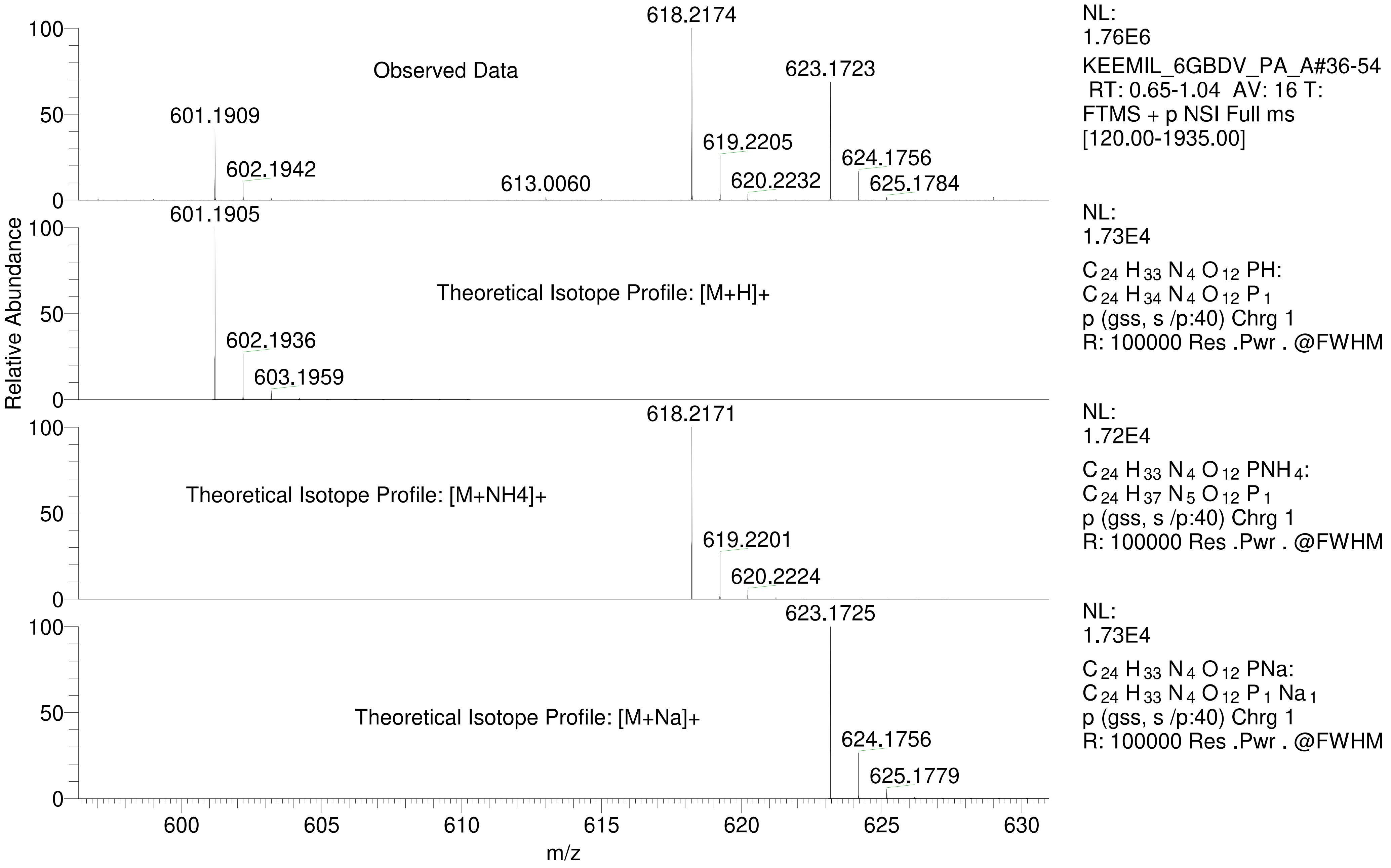

| | | | | |
|---|---|---|---|---|
| **Isotope:** | **Min. .. Max.** | | | |
| **14 N** | **0....16** | | | |
| **16 O** | **0....20** | | | |
| **12 C** | **0....100** | | | |
| **1 H** | **0....120** | | | |
| **23 Na** | **1....1** | | | |
| **31 P** | **0....2** | | | |
| **Tolerance Window:** | **+- 5.00 ppm** | | | |
| **Db/Ring Equiv:** | **-10.. 500** | | **N-Rule:** | **Do not use** |
| **Fits:** | **500** | | **Charge:** | **1** |

| **Mass** | **Theoretical Mass** | **Delta [ppm]** | **RDB** | **Composition** |
|---|---|---|---|---|
| 601.1909 | 601.1908 | 0.1 | 6.0 | $C_{20} H_{37} O_{12} N_5 P_2$ |
| | 601.1908 | 0.1 | 11.5 | $C_{19} H_{31} O_7 N_{12} P_2$ |
| | 601.1910 | -0.2 | 3.5 | $C_9 H_{30} O_{13} N_{16} P_1$ |
| | 601.1910 | -0.2 | -2.0 | $C_{10} H_{36} O_{18} N_9 P_1$ |
| | 601.1911 | -0.3 | 32.5 | $C_{43} H_{25} O_2 N_2$ |
| | 601.1907 | 0.3 | 2.5 | $C_{14} H_{33} O_{18} N_8$ |
| | 601.1907 | 0.3 | 8.0 | $C_{13} H_{27} O_{13} N_{15}$ |
| | 601.1905 | 0.6 | 10.5 | $C_{24} H_{34} O_{12} N_4 P_1$ |
| | 601.1905 | 0.6 | 16.0 | $C_{23} H_{28} O_7 N_{11} P_1$ |
| | 601.1914 | -0.8 | -6.5 | $C_6 H_{39} O_{18} N_{10} P_2$ |
| | 601.1914 | -0.8 | 28.0 | $C_{39} H_{28} O_2 N_3 P_1$ |
| | 601.1903 | 0.9 | 18.5 | $C_{34} H_{35} O_6 P_2$ |
| | 601.1903 | 0.9 | 24.0 | $C_{33} H_{29} O_1 N_7 P_2$ |
| | 601.1916 | -1.1 | 25.5 | $C_{28} H_{21} O_3 N_{14}$ |
| | 601.1916 | -1.1 | 20.0 | $C_{29} H_{27} O_8 N_7$ |
| | 601.1916 | -1.1 | 14.5 | $C_{30} H_{33} O_{13}$ |
| | 601.1902 | 1.1 | 15.0 | $C_{28} H_{31} O_{12} N_3$ |
| | 601.1902 | 1.1 | 20.5 | $C_{27} H_{25} O_7 N_{10}$ |
| | 601.1917 | -1.3 | 23.5 | $C_{35} H_{31} O_2 N_4 P_2$ |
| | 601.1900 | 1.5 | 28.5 | $C_{37} H_{26} O_1 N_6 P_1$ |
| | 601.1900 | 1.5 | -6.0 | $C_4 H_{37} O_{17} N_{13} P_2$ |
| | 601.1919 | -1.6 | 21.0 | $C_{24} H_{24} O_3 N_{15} P_1$ |
| | 601.1919 | -1.6 | 15.5 | $C_{25} H_{30} O_8 N_8 P_1$ |
| | 601.1919 | -1.6 | 10.0 | $C_{26} H_{36} O_{13} N_1 P_1$ |
| | 601.1921 | -1.9 | 7.5 | $C_{15} H_{29} O_{14} N_{12}$ |
| | 601.1921 | -2.0 | 2.0 | $C_{16} H_{35} O_{19} N_5$ |
| | 601.1897 | 2.0 | 33.0 | $C_{41} H_{23} O_1 N_5$ |
| | 601.1897 | 2.0 | -1.5 | $C_8 H_{34} O_{17} N_{12} P_1$ |
| | 601.1922 | -2.1 | 16.5 | $C_{20} H_{27} O_3 N_{16} P_2$ |
| | 601.1922 | -2.1 | 11.0 | $C_{21} H_{33} O_8 N_9 P_2$ |
| | 601.1922 | -2.1 | 5.5 | $C_{22} H_{39} O_{13} N_2 P_2$ |

==================================================================
Acq. Operator : SYSTEM    Seq. Line : 3
Sample Operator : SYSTEM
Acq. Instrument : Prep LC    Location : P2-A-03
Injection Date : 10/3/2023 12:50:43 PM    Inj : 1
Inj Volume : 50.000 µl
Method : C:\Users\Public\Documents\ChemStation\1\Data\Aisling\ANC BS6 iii percent purity  2023-10-03 11-36-25\ANC-BS-6iii Percent Purity.M (Sequence Method)
Last changed : 10/3/2023 11:33:37 AM by SYSTEM
Method Info : Polaris 5 C15-A 250 x10mm SN 593740

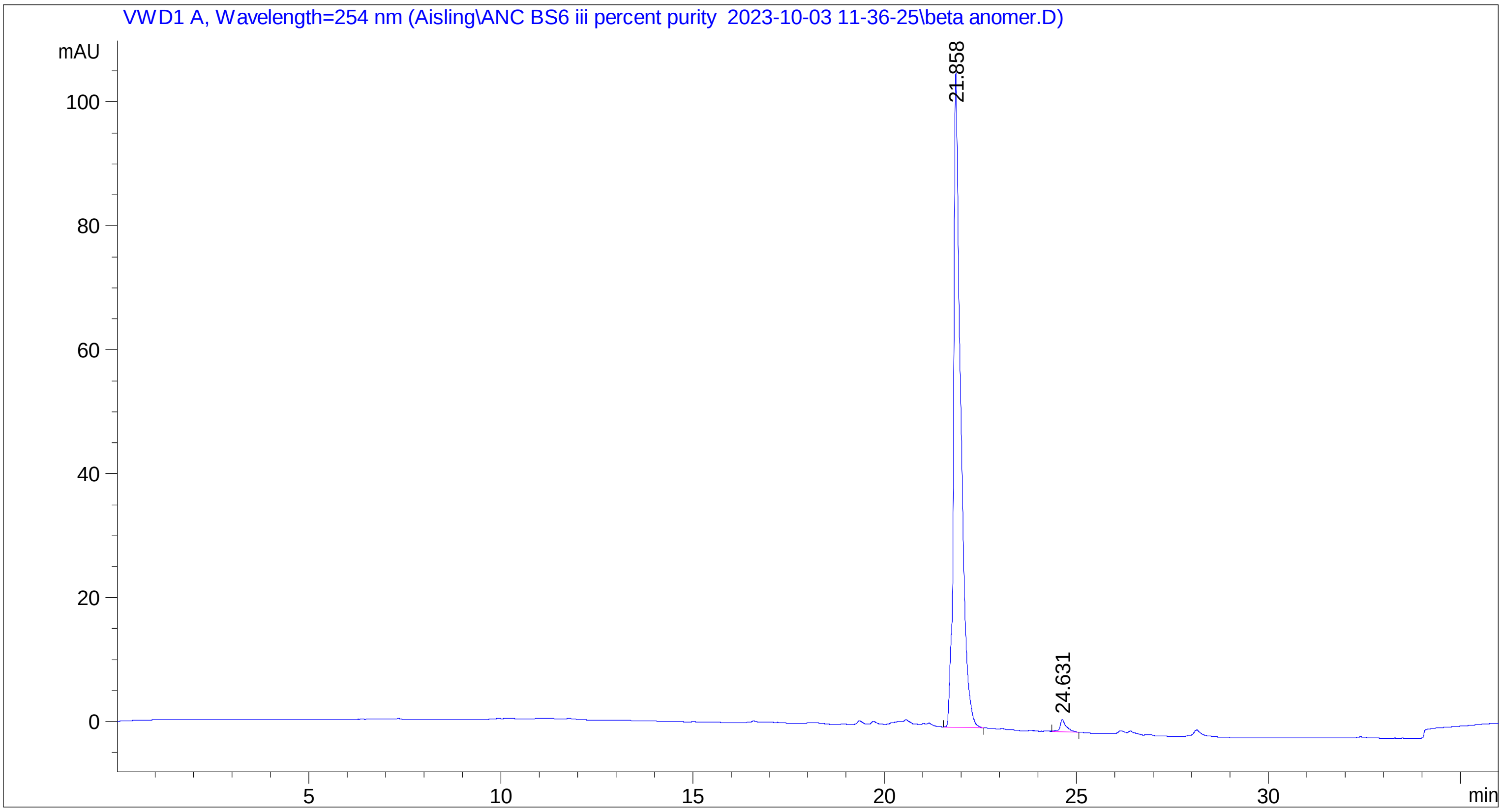


==================================================================
Fraction Information
==================================================================
No Fractions found.
==================================================================
==================================================================
==================================================================
Area Percent Report
==================================================================

Sorted By : Signal
Multiplier : 1.0000
Dilution : 1.0000
Use Multiplier & Dilution Factor with ISTDs

Signal 1: VWD1 A, Wavelength=254 nm

| Peak # | RetTime [min] | Type | Width [min] | Area [mAU*s] | Height [mAU] | Area % |
|---|---|---|---|---|---|---|
| 1 | 21.858 | BB | 0.1794 | 1359.55212 | 105.49213 | 98.4411 |
| 2 | 24.631 | BB | 0.1543 | 21.52952 | 1.94319 | 1.5589 |
| Totals : | | | | 1381.08164 | 107.43532 | |

=====================================================================

*** End of Report ***
